# Regulatory grammar design of tissue-specific enhancers across species

**DOI:** 10.64898/2026.11.17.767007

**Authors:** Liwen Yao, Mai Ngoc Tran, Alexandra S. Ruffus, Le Xiong, Aniket Jana, Sanaa Bhuleshkar, Ronald A. Conlon, Takuya F. Sakaguchi, Liangqi Xie

## Abstract

Enhancers encode tissue-specific gene expression through the combinatorial identity, number, and arrangement of transcription factor motifs, collectively known as regulatory grammar. However, existing deep-learning methods directly optimize nucleotide sequences, leaving enhancer regulatory grammar implicit. Here we introduce REgulatory Grammar Architecture via reinforcement Learning (REGAL), a framework that explicitly programs enhancer grammar under sequence-to-function oracles. REGAL generated tissue-specific enhancers across species using the same placement architecture with interchangeable oracles and motif vocabularies. Experimental validation and perturbation revealed that tissue specificity requires coordinated target-tissue activation and off-target repression, depends on both motif composition and spatial organization, and employs distinct regulatory strategies across tissues. REGAL-designed enhancers drove predicted lineage-specific expression in transgenic mouse embryos, outperformed a nucleotide-level optimizer in human cells, and matched or exceeded leading designs in zebrafish and *Drosophila* at notably shorter length. Together, these findings establish regulatory grammar as a programmable and experimentally testable substrate underlying enhancer design and specificity.

## Introduction

Enhancers control gene expression in specific cell types, tissues, and developmental stages through combinatorial binding of sequence-specific transcription factors (TFs). Enhancer activity is encoded by the identity, copy number, spacing, orientation, and organization of TF binding motifs, features that collectively constitute the enhancer’s regulatory grammar^1–3^. Leveraging this grammar would move enhancer biology from reading natural regulatory sequences to programming regulatory logic, enabling the rational design of compact, cell-type-specific enhancers^3,4^, with valuable applications in gene therapy to minimize off-target toxicity^5^ and maximize delivery payload capacity^6^.

Emerging deep-learning models can accurately predict enhancer activity from DNA sequence^7,8^. Coupling these predictive models with sequence optimization has produced tissue-specific enhancers in *Drosophila melanogaster*^9,10^, human cell lines^4,11^, and transgenic mouse embryos^12^. These approaches, however, optimize enhancers as nucleotide sequences, leaving the regulatory grammar that specifies enhancer function latent and inferred retrospectively by attribution or motif scanning. A more fundamental question therefore remains: can regulatory grammar itself be the substrate of design? Just as human language conveys meaning through words arranged according to syntax, cells decode enhancer function through TF motifs organized into regulatory grammar^13^. Designing explicitly in this grammar space, rather than at the nucleotide level, could constrain optimization to biologically meaningful variables, compress regulatory logic into compact elements, and make the basis of enhancer function directly explainable. However, whether this representation can support effective, efficient, and interpretable enhancer design remains unknown (**Fig. 1a**).

**Fig. 1.**
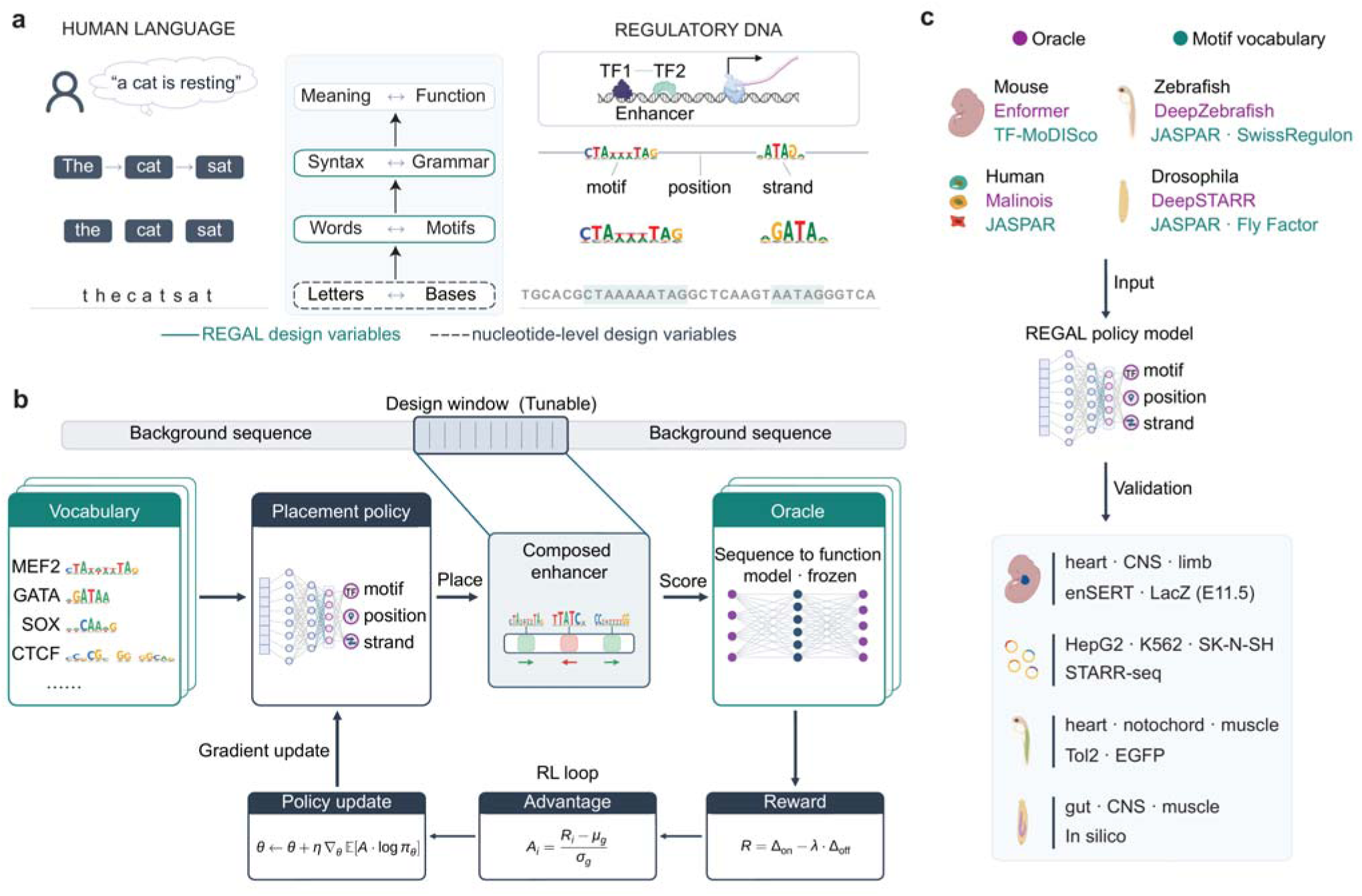
REGAL framework for cross-species regulatory grammar design. **a**, Conceptual analogy between human language and regulatory DNA. Meaning corresponds to enhancer function, syntax to regulatory grammar, words to transcription factor (TF) motifs, and letters to nucleotide bases. Teal outlines mark the levels REGAL designs — motif identity, order, orientation, and spacing — whereas the dashed outline indicates bases optimized by conventional nucleotide-level methods. **b,** The REGAL architecture. An autoregressive placement policy selects motif identity, position, and strand within a tunable window in a fixed background sequence. A frozen sequence-to-function oracle evaluates and scores the composed enhancer, and group-relative policy optimization updates the policy using rewards and group-standardized advantages. Equations are schematic. **c,** REGAL designs were evaluated experimentally in mouse embryos, human cell lines and zebrafish, and *in silico* in Drosophila. Δ, the oracle-predicted activity of the composed enhancer minus that of the background sequence; λ, the off-target penalty coefficient; Ri and Ai, the reward and group-standardized advantage of sampled design i; μg and σg, the mean and standard deviation of rewards within its sampling group; θ, policy parameters; η, learning rate; πθ, the policy; E11.5, embryonic day 11.5; EGFP, enhanced green fluorescent protein; STARR-seq, self-transcribing active regulatory region sequencing; CNS, central nervous system.

Here we introduce REGAL (REgulatory Grammar Architecture via reinforcement Learning), which uses a sequence-to-function oracle to directly optimize TF motif composition and spatial organization underlying TF binding and cooperativity^1–3^. Grammar-level enhancer design presents a vast combinatorial problem, assembling discrete choices of motif identity, position and orientation across multiple motifs without predefined rules for their optimal arrangements. REGAL addresses this challenge and formulates enhancer design as a reward-driven optimization through reinforcement learning, which enables a policy to search this combinatorial grammar space according to explicitly defined regulatory objectives while keeping the motif vocabulary, placement policy, predictive oracle and design objective modular and interchangeable (**Fig. 1b**).

Using an oracle trained on transgenic mouse enhancer annotations, REGAL generated enhancers that were both tissue-active and tissue-discriminative. Contrastive design revealed that lineage specificity requires both lineage activation and off-target suppression, whereas motif perturbation demonstrated both motif composition and spatial arrangement are key to enhancer function. The contribution of individual grammatical feature differed by lineage, with cardiac specificity relied more strongly on bifunctional motif content and neural specificity on combinatorial motif arrangement. REGAL-designed enhancers drove the intended-lineage expression in transgenic mouse embryos. Holding the placement policy architecture fixed while exchanging only the oracle and motif vocabulary, REGAL outperformed a nucleotide-level optimizer targeting the same oracle in human cell lines and matched or exceeded leading design methods in zebrafish and *Drosophila* while producing substantially shorter enhancers (**Fig. 1c**). Together, these results establish regulatory grammar as an explicit, programmable, and experimentally testable object and reveal fundamental principles governing enhancer regulatory specificity.

## Results

### REGAL: a modular framework for grammar-level enhancer design

REGAL is a modular reinforcement-learning framework that designs enhancers by composing TF motifs in regulatory grammar space rather than directly optimizing nucleotide sequences (**Fig. 1b**). REGAL consists of three modular components. The motif vocabulary defines the regulatory alphabet. The placement policy is an autoregressive Transformer^14^ decoder that sequentially assembles motifs onto a background sequence by specifying motif identity, position, and strand until termination or reaching a preset limit (**Extended Data Fig. 1a**). The sequence-to-function oracle evaluates each design and returns a scalar reward summarizing either predicted target-tissue activity alone or a contrastive combination of on- and off-target activities. The placement policy is updated using group-relative policy optimization (GRPO), which compares the rewards of candidate grammatical designs sampled from the same background relative to their group mean^15^.

The motif vocabulary, oracle, and reward objective can each be exchanged independently, whereas the placement policy is the only trained component with its architecture unchanged across tasks and species. This modular design allows REGAL to generalize across diverse biological systems, thereby functioning not as a task-specific model but as a flexible framework for regulatory grammar-level enhancer design (**Fig. 1c**).

### Benchmarking the design oracle and deriving an oracle-coupled motif vocabulary

To benchmark REGAL for enhancer design, we first established a tissue-specific sequence-to-function oracle and corresponding motif vocabulary. Using Enformer^7^, DNABERT-2^16^, HyenaDNA^17^, and DeepSTARR^8^ on VISTA^18^ enhancers, which have been experimentally validated in transgenic mouse embryos. Enformer achieved the highest discriminative performance, with a mean area under the receiver operating characteristic (AUROC) of 0.77, compared with 0.54–0.65 for the other models (**Fig. 2a, Extended Data Figs. 1b and 2a,b, Supplementary Tables 1 and 2**). Enformer predicted activity changes correlated with experimentally measured effects from an independent *in vivo* enhancer-mutagenesis dataset^19^ (r = 0.61; **Fig. 2b, Extended Data Fig. 2c**), and genome-wide high-scoring regions were strongly enriched in tissue-matched ChromHMM enhancer states^20^ (**Extended Data Fig. 2d**). We therefore selected Enformer as the design oracle.

**Fig. 2.**
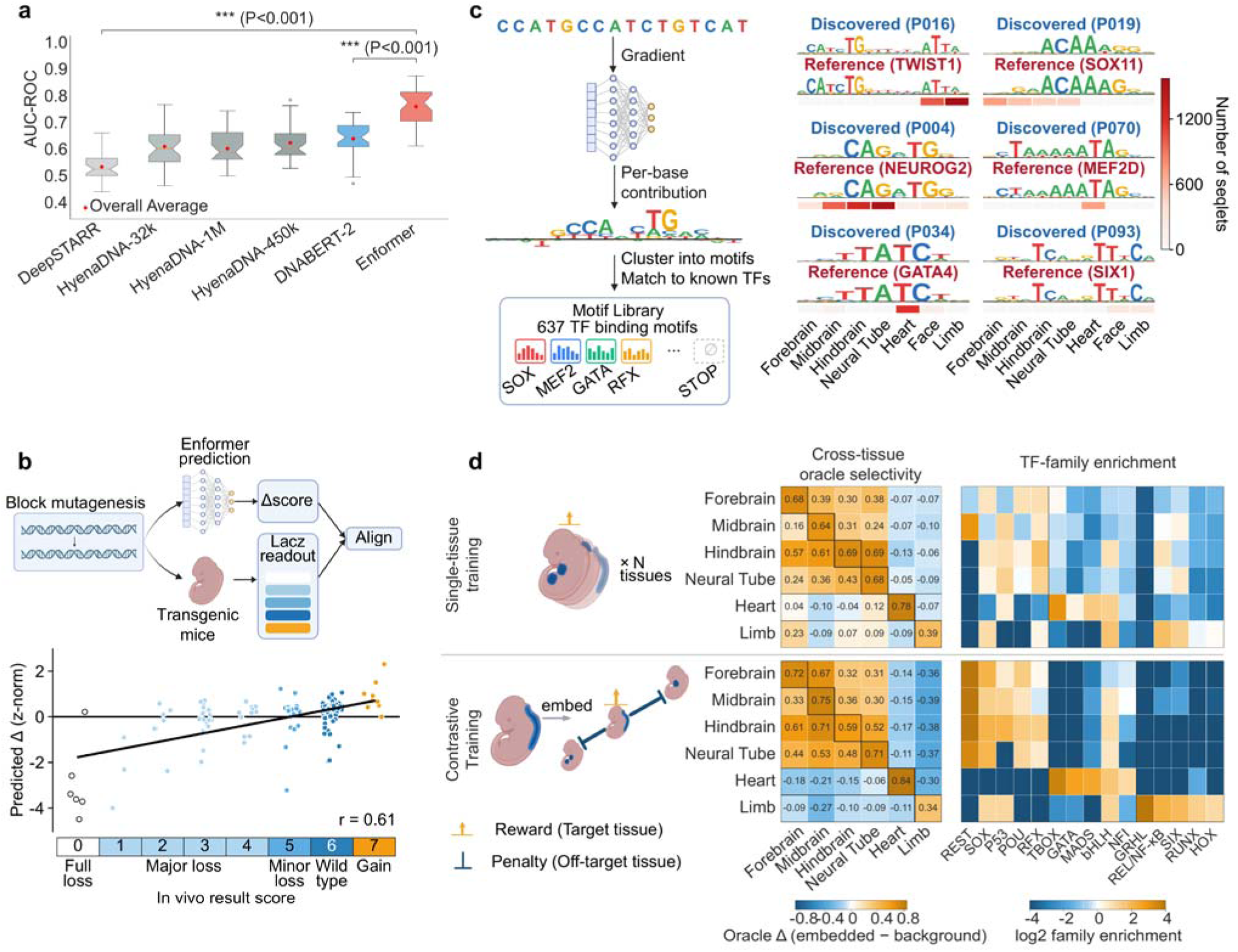
Oracle benchmarking and motif-guided tissue-specific mouse enhancer design. **a**, Performance of six oracles on VISTA enhancer classification (held-out mean area under the receiver operating characteristic curve (AUROC)). n = 35 per model (seven E11.5 tissues × five seeds). Boxes show the median and interquartile range (IQR), whiskers extend to 1.5 × IQR, diamonds indicate means, and circles indicate outliers. Enformer outperformed DeepSTARR and DNABERT-2 in two-sided independent-samples t-tests (***P < 0.001). **b,** Independent validation of Enformer using 167 block mutations across seven developmental enhancers assayed *in vivo* in transgenic mouse embryos. Predicted mutational effects were normalized within enhancer and compared with the measured functional scores. In the lower scatter plot, the black line shows the least-squares fit (Pearson r = 0.61, P < 0.001). **c,** Construction of the mouse motif vocabulary. Integrated-gradients attribution profiles from the seven tissue oracles were clustered by TF-MoDISco to identify 637 motifs, which were matched to known TF motifs using Tomtom. Representative discovered motifs and their reference matches are shown together with their tissue-specific seqlet contributions. **d,** Cross-tissue activity and TF-family usage learned by single-tissue and contrastive REGAL policies. Heatmaps show the mean oracle response across tissues (left) and TF-family enrichment among designed enhancers (right). Contrastive optimization increased target-versus-off-target separation and produced tissue-specific shifts in motif-family usage. Values are summarized across 24 independently trained policies per tissue.

Because REGAL composes enhancers from TF motifs rather than nucleotide sequences, we performed attribution analysis to derive an oracle-coupled motif vocabulary. VISTA fine-tuning endowed the oracle with tissue-specific motif sensitivity. Integrated gradients revealed attribution patterns positive for tissue-matched cognate TF motifs but neutral or negative elsewhere (**Extended Data Fig. 2e,f**)^21^. Applying TF-MoDISco to these attributions identified 637 unified motifs comprising both activating (326) and repressing (311) regulatory elements (**Fig. 2c**)^22^. Of these, 521 (81.8%) matched known TF binding motifs by extensive database query^23–24^, whereas the remaining 116 (18.2%) formed previously unannotated clusters (**Supplementary Data 1**).

### REGAL recapitulates mouse developmental enhancer grammars

We first applied REGAL to design enhancers for six embryonic day 11.5 (E11.5) mouse tissues: forebrain, midbrain, hindbrain, neural tube, heart, and limb. The single-tissue objective rewarded target-tissue activity alone, whereas the contrastive objective also penalized activity in non-target tissues. Under the contrastive objective, the predicted target-tissue oracle activity (Δ) increased rapidly and plateaued within preset optimization iterations in all six tissues, indicating convergence (**Extended Data Fig. 3a**).

Across tissues, nearly all single-tissue REGAL designs readily separated from motif-count-matched random sequences (AUROC 0.855–0.993) and increased predicted target-tissue activity, whereas random sequences produced little change (**Extended Data Fig. 3b**). Across 24 independent optimization runs, policies converged to similar reward levels (**Extended Data Fig. 3c**) while generating distinct enhancer sequence repertoires (**Extended Data Fig. 3d**). Thus, REGAL converged reproducibly without collapsing sequence diversity.

We next asked whether REGAL revealed developmental regulatory grammar. Although every design policy optimized the same 637-motif vocabulary under an target-tissue-only reward and received no explicit TF–tissue supervision, the resulting designs became enriched for lineage-appropriate TF families (**Fig. 2d**). Heart designs preferentially incorporated canonical cardiac regulators, including HAND1^25^, MEF2, and TBX5; neural designs enriched factors including SOX2, REST and ATOH1^26^; limb designs enriched motifs associated with skeletal and appendicular regulators, including SOX9, SIX1, and posterior HOXD^27^. Thus, tissue activity alone was sufficient to recover recognizable lineage-associated regulatory programs.

Introducing the contrastive objective further refined the grammar. Heart designs became concentrated on canonical cardiac regulator motifs, including T-box, GATA, and MEF2-family motifs^28^, whereas limb designs further gained TWIST 1 and enriched HOX motifs^29^. Neural lineages preferentially incorporated REST and CTCF motifs^30^, consistent with roles of transcriptional suppression^31^ and higher-order chromatin organization^32^ in reinforcing tissue specificity. Correspondingly, cross-tissue oracle scoring recapitulated expected lineage relationships: neural enhancers cross-activated the other neural tissues (mean Δ +0.39) while suppressing heart and limb (deepening to −0.26), whereas heart enhancers selectively activated heart (+0.78) with minimal off-target activity (−0.01) (**Fig. 2d**). The contrastive objective increased predicted tissue specificity across all six tissues while largely preserving target-tissue activity, with only modest reductions in hindbrain and limb (−15.2% and −12.2%; **Extended Data Fig. 3e,f**). These results suggest that optimization in the regulatory grammar space recapitulates developmental enhancer grammars.

### REGAL designs tissue-discriminative enhancers and reveals lineage-specific regulatory grammar

To evaluate enhancer activity and specificity, we performed assay for transposase-accessible chromatin using self-transcribing active regulatory region sequencing (ATAC-STARR-seq)^33^ in mouse N2a (neural), HL-1 (cardiac), and embryonic stem cells (mESCs) (**Extended Data Fig. 4a–c**). Lineage-specific elements, defined as active in one line but inactive in the other two, were significantly enriched among the top 5,000 predicted regions for the corresponding neural and cardiac tissues relative to random regions (**Extended Data Fig. 4d**), supporting N2a and HL-1 as appropriate neural- and cardiac-lineage reporter contexts.

We next synthesized a 9,500-oligo library containing 8,000 REGAL-designed enhancers (1,000 single-tissue and 1,000 contrastive designs for each of four target tissues) plus 1,500 controls (**Supplementary Data 2**) and measured their activity by STARR-seq^34^ in both N2a and HL-1 cells (**Fig. 3a, Extended Data Fig. 5a,b**). Activities were normalized to lineage-matched natural-enhancer and designed-null controls, and specificity was quantified as the ratio of on-to off-target activity (see **Methods**).

**Fig. 3.**
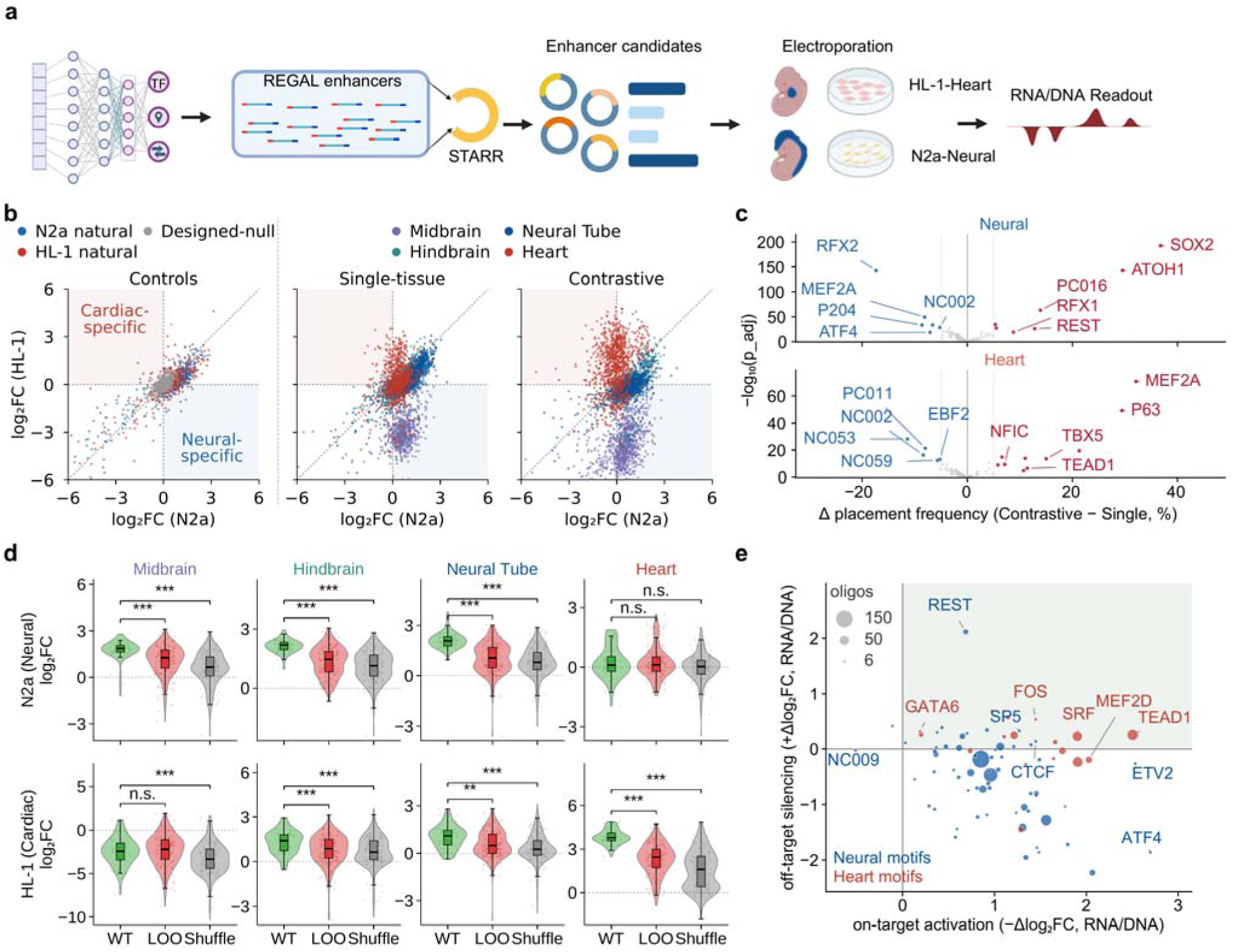
REGAL designs cell-discriminative mouse enhancers with motif-dependent activity. **a**, STARR-seq validation of REGAL-designed enhancers in N2a (neural) and HL-1 (cardiac) cells. Enhancer activity was quantified as reporter RNA normalized by input DNA. Three paired libraries were generated per cell line. **b,** Enhancer activity in N2a and HL-1 cells, relative to the median activity of designed-null controls, for natural and designed-null controls (left) and single-tissue and contrastive REGAL designs (right). For each cell line, the fraction of active designs was compared between single-tissue and contrastive designs using a two-sided Fisher’s exact test (both P < 10□□□). **c,** Difference in motif usage between contrastive and single-tissue designs for pooled neural enhancers and heart enhancers. Significance was assessed using two-sided Fisher’s exact tests with Benjamini–Hochberg correction; significant motifs were highlighted. **d,** Functional validation of motif composition and spatial arrangement by leave-one-out (LOO) motif deletion and arrangement shuffling. Two-sided Mann–Whitney U tests compared each perturbation versus WT. ***P < 0.001; **P < 0.01; n.s., not significant. Violin plots show the distributions, with boxes indicating the median and IQR; whiskers, 1.5 × IQR. **e,** Contribution of motifs to tissue specificity derived from the LOO perturbation series. Each point represents a motif positioned by its mean contribution to on-target activation and off-target silencing; point size reflects the number of supporting oligos. The shaded quadrant identifies motifs with dual activating and silencing functions. WT, wild-type REGAL design.

Single-tissue REGAL designs generated active enhancers in the intended lineages (44.0% of midbrain, 41.5% of hindbrain, and 58.5% of neural tube designs were active in N2a, whereas 24.2% of heart designs in HL-1), comparable (0.8–1.1 fold) to matched natural enhancers. However, these designs exhibited substantial off-target activity: 28.6% neural tube designs and 14.5% hindbrain designs were active in HL-1, and heart designs showed only a modest preference for cardiac over neural cells (24.2% versus 16.4%). Thus, maximizing target-lineage activity alone was insufficient to achieve desired tissue discrimination (**Fig. 3b**).

Incorporating a contrastive objective markedly improved specificity across all four target tissues (**Fig. 3b**). Midbrain, neural tube, and hindbrain designs achieved 130-fold (26.0% in N2a versus 0.2% in HL-1), 7.6-fold (44.6% versus 5.9%), and 5.8-fold (49.6% versus 8.6%) selectivity, respectively, compared with 9.6-fold, 2.0-fold, and 2.9-fold for matched single-tissue designs. Heart designs similarly improved from 1.5-fold to 6.3-fold selectivity, while on-target activity increased from 24.2% to 60.8% (**Extended Data Fig. 5c**). Off-target activity fell to or below the designed-null baseline, whereas on-target activity remained comparable to natural enhancers and, for heart designs, reached 3.6-fold higher activity than the natural cardiac control. These findings from grammar-level design indicate that enhancer specificity is governed not only by activating on-target programs but also by suppressing off-target activity.

### REGAL reveals tissue-discriminative enhancer grammar

To determine how contrastive optimization improves tissue specificity, we compared motif usage learned by single-tissue and contrastive designs. Contrastive optimization consistently rewired motif composition in a lineage-specific manner. Neural designs more frequently incorporated neural regulator motifs including SOX2, ATOH1, and RFX1, while depleting cardiac-associated MEF2-family and RFX2 motifs. REST^30^ was also enriched across pooled neural designs, particularly in midbrain enhancers (**Fig. 3c**).

To test whether these learned enhancer grammars are functionally required, we generated a matched STARR-seq library comprising wild-type enhancers, leave-one-out (LOO) variants lacking single motifs, and variants with identical motif composition but randomized positions and orientations (shuffled) (**Extended Data Fig. 5d,e; Supplementary Data 3**). Both motif composition and spatial arrangement proved required for enhancer activity. Pooled across the neural designs, motif ablation reduced the fraction of active enhancers from 98.0% to 66.5%, whereas position-shuffling reduced activity to 53.9%. Among heart designs, both perturbations similarly reduced enhancer activity. Positional shuffling impaired enhancer activity despite preserving motif content, demonstrating an independent role of motif spatial organization. Consistent with a lineage-specific mechanism, motif ablation significantly reduced heart enhancer activity in HL-1 cells but had no significant effect in N2a cells (**Fig. 3d**).

The motif-ablation library further enabled quantification of the functional contribution of individual motifs. The strongest on-target dependencies recovered were lineage regulators, including TEAD1 and SRF^25^ in heart enhancers, and SOX2 and ATOH1^26^ in neural enhancers. Notably, nearly half of the heart motifs (8 of 18, 44%) were bifunctional, simultaneously promoting cardiac activity while suppressing neural activity. For example, removing TEAD1, SRF, or MEF2A decreased enhancer activity in HL-1 cells but increased activity in N2a cells, whereas REST exhibited the reciprocal behavior, supporting neural activity while suppressing cardiac activity^30^ (**Fig. 3e, Extended Data Fig. 5f**). In contrast, only 7 of 60 motifs (12%) in neural designs were bifunctional. These results suggest distinct grammatical strategies for enhancer specificity in different tissues: cardiac enhancers frequently use bifunctional motifs that couple on-target activation with off-target repression, whereas neural enhancers rely more on combinatorial motif composition and spatial organization.

### REGAL-designed enhancers drive tissue-specific expression in transgenic mouse embryos

We next tested whether REGAL-designed enhancers function *in vivo*. Before experimental testing, we randomly selected nine designs from the top 5% of each target-tissue pool ranked by Enformer-predicted activity: three for heart (HT1, HT2, HT3), three for limb (LB7, LB8, LB9), and one each for midbrain (MB4), hindbrain (HB5), and forebrain (FB6), representing three central nervous system (CNS) subregions. Each enhancer was site-specifically integrated into the mouse genome by enSERT and assayed for *lacZ* reporter activity by whole-mount X-gal staining in E11.5 embryos^19,35,36^ (**Figs. 1c and 4a, Supplementary Data 4**). Of 273 imaged embryos, 186 had PCR-confirmed transgene integration and were included for analysis. All nine REGAL-designed enhancers drove reporter expression in their intended lineage (9/9), with 153 of 186 embryos (82.3%) showing target-tissue activity; the remaining 33 embryos were reporter-negative. No embryo showed majority-supported off-target staining outside the intended lineage (0/186; **Fig. 4b**).

**Fig. 4.**
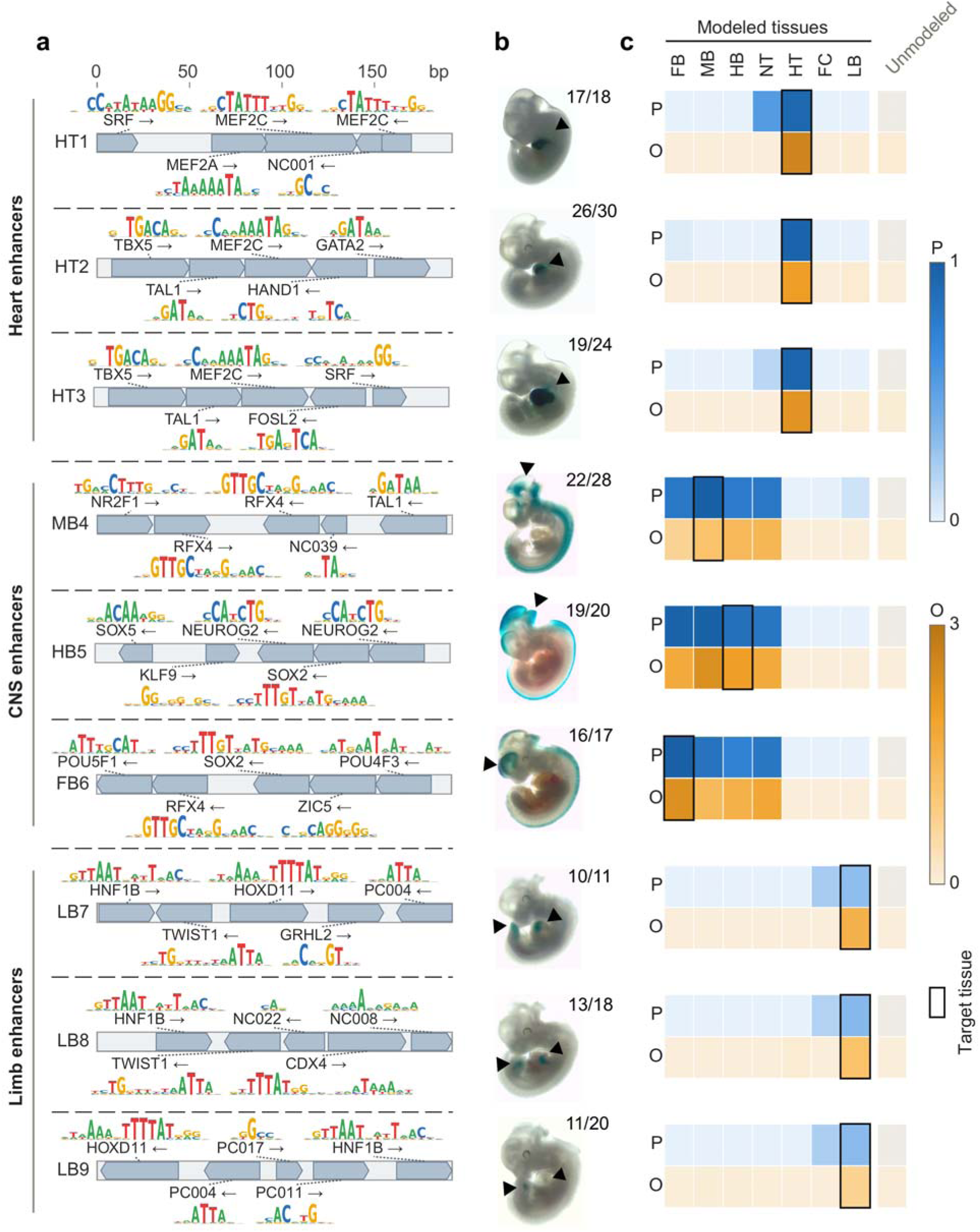
*In vivo* validation of REGAL-designed enhancers in transgenic mouse embryos. **a,** Motif grammar maps of nine 192-bp REGAL-designed enhancers targeting heart, CNS or limb. Pentagons indicate motif position and orientation, and sequence logos show motif information content. Labels prefixed NC or PC are de novo vocabulary motifs without a Tomtom match to a known transcription factor. **b,** Representative X-gal-stained (*lacZ* reporter) E11.5 embryos. Black arrowheads indicate target tissue staining. Numbers indicate the fraction of analyzed embryos showing target-tissue staining for each design. **c,** Enformer-predicted and observed activities across the modeled tissues. Blue indicates predicted activity and amber indicates observed activity; outlined boxes mark the target tissue. Across the 63 design–tissue combinations, predicted and observed activities were strongly correlated (Pearson r = 0.909, P = 8.3 × 10□²□). Of 273 imaged embryos, 186 with PCR-confirmed transgene integration were analyzed; 153 of 186 were scored as target positive, and no design showed majority-supported off-target staining. FB, forebrain; MB, midbrain; HB, hindbrain; NT, neural tube; HT, heart; FC, face; LB, limb.

To quantify enhancer activity objectively, three independent assessors blinded to construct identity scored X-gal staining intensity on a 0–3 scale across a 15-structure anatomical panel (**Fig. 4c**). REGAL-designed enhancers produced robust on-target reporter expression (mean score, 1.66) with negligible off-target activity (mean score, 0.008; no off-target structure above 0.093). Across all 63 design-by-tissue combinations, predicted and observed profiles correlated strongly (Pearson r = 0.909). Heart and limb designs drove sharply restricted expression within their target tissues, whereas the three neural designs reproducibly activated expression across all four CNS subregions, consistent with their shared developmental programs and predicted CNS activity by the oracle. Together, these results demonstrate that REGAL-designed enhancer grammars encode robust and predictable lineage-restricted regulatory activity *in vivo*.

### REGAL programs cell-type-specific human enhancers and outperforms nucleotide-level design

To test whether REGAL generalizes beyond mouse and to directly compare regulatory grammar with established nucleotide-level design, we retained the placement-policy architecture and replaced only its oracle (Malinois^4^) and motif vocabulary with human TF motifs (**Fig. 1c**). We targeted three human cell lines, K562, HepG2, and SK-N-SH. Under the contrastive objective, all three policies converged on cell-type-specific motif repertoires while increasing predicted activity in the target cell and suppressing off-target activity toward baseline (**Extended Data Fig. 6a**)^37^.

We next asked whether REGAL learned combinatorial regulatory grammar rather than merely accumulating individual high-scoring motifs. Motif pairs co-placed by REGAL showed significantly greater oracle-predicted synergy than random control pairs drawn from the same motif vocabulary in all three cell lines (**Fig. 5a**). Moreover, when constrained to place a fixed motif pair and optimize its spacing and orientation, REGAL recovered motif distances closely matching oracle-preferred spacing across 30 motif pairs (median absolute difference 3.5 bp, r = 0.75; **Fig. 5b**). These results indicate that REGAL captures cooperative motif interactions and their spatial preferences.

**Fig. 5.**
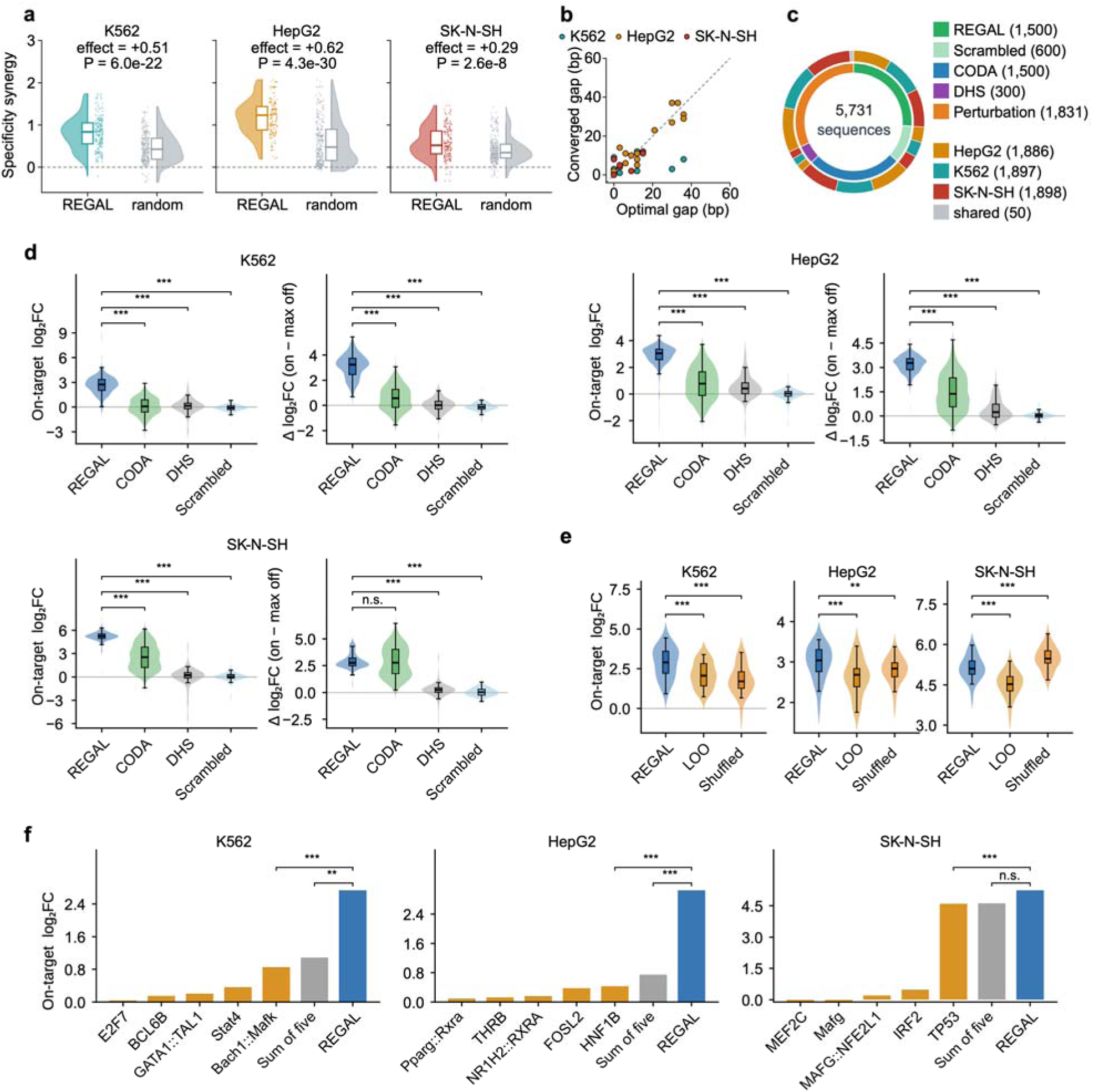
Motif-grammar design and validation of REGAL enhancers in human cells. **a**, Oracle-predicted specificity synergy of REGAL-co-placed versus random pairs in K562, HepG2 and SK-N-SH. Comparisons were performed using two-sided Mann–Whitney U tests. **b,** Recovery of oracle-preferred motif spacing by forced-pair REGAL optimization. Each point compares the oracle-optimal spacing with the spacing learned by the policy; the dashed line indicates equality (Pearson r = 0.75, P = 2.3 × 10). **c,** Composition of the 5,731-oligo human STARR-seq library. The inner ring indicates sequence class and the outer ring the target cell line. **d,** STARR-seq measured on-target log FC and specificity of REGAL enhancers compared with CODA designs, natural DHS elements and scrambled controls in K562, HepG2 and SK-N-SH. Specificity was defined as the on-target log_2_FC minus the maximum off-target log_2_FC. Comparisons with REGAL were performed using two-sided Mann–Whitney U tests. **e,** Paired validation of REGAL grammar by leave-one-out (LOO) motif deletion and positional shuffling. Perturbed oligos were compared with their matched intact REGAL parents using two-sided Wilcoxon signed-rank tests. **f**, On-target log_2_FC of the five constituent motifs tested individually, the sum of their individual log_2_FC values, and the corresponding intact REGAL enhancer in K562, HepG2 and SK-N-SH cells. Boxes in a, d and e show the median and IQR, with whiskers extending to 1.5 × IQR. ***P < 0.001; **P < 0.01; n.s., not significant. DHS, DNase I hypersensitive site.

We then benchmarked REGAL with two published enhancer-design frameworks: CODA^4^, a nucleotide-level sequence optimizer and Guided Optimization of Cis-Regulatory Elements (GO-CRE)^38^, both targeting the same Malinois oracle. *In silico*, REGAL designs achieved higher predicted activity and specificity than CODA, and higher predicted activity than GO-CRE in all three cell lines and higher specificity in K562 and HepG2 (**Extended Data Fig. 6b,c**). Because optimization against predictive models can drive designed sequences away from natural sequence distributions^39^, we examined global sequence composition such as CpG density associated with regulatory activity and environment^40^. REGAL designs showed substantially less CpG enrichment than CODA and GO-CRE across all three cell types and approached natural DNase I hypersensitive site (DHS) levels in K562 and HepG2 (**Extended Data** Fig. 7a).

To experimentally compare designs, we synthesized a 5,731-oligo library comprising REGAL and CODA enhancers, natural DHS elements^4^, scrambled controls, and motif-perturbation variants and measured activity by STARR-seq in all three lines (**Fig. 5c, Extended Data Fig. 7b,c, Supplementary Data 5**). REGAL consistently outperformed CODA, producing substantially stronger enhancer activity in K562 (median STARR-seq log₂ fold change (log₂FC), 2.74 versus 0.06), HepG2 (3.05 versus 0.78), and SK-N-SH (5.24 versus 2.52) (**Fig. 5d**). Nearly all REGAL enhancers were active in their cognate lineage (95.7% in K562, 96.8% in HepG2 and 99.8% in SK-N-SH), whereas K562 and HepG2 designs exhibited minimal activity in non-cognate lines (0.7% and 1.3%), demonstrating robust cell-type specificity (**Fig. 5d**, **Extended Data Fig. 7c**). SK-N-SH designs retained measurable activity in HepG2 and showed specificity gap comparable to that of CODA (medians 2.76 versus 2.77) despite their much higher cognate activity. Because REGAL and CODA spanned overlapping ranges of predicted activity in K562 and HepG2, we compared measured activity with matched predicted scores. REGAL designs consistently yielded higher experimental activity than CODA, indicating greater prediction-to-measurement fidelity. Natural DHS elements showed a similar advantage over CODA, while REGAL further exceeded natural DHS in K562 (**Extended Data Fig. 7d**). Measured CODA and DHS activity correlated with previously reported lentiviral massively parallel reporter assay (lentiMPRA) measurements (Spearman 0.63–0.75; **Extended Data Fig. 7e**)^41^, supporting cross-assay reproducibility. Thus, REGAL programmed experimentally validated cell-type-active and -discriminative human enhancers, outperforming nucleotide-level design.

We then tested whether enhancer activity depended on the designed regulatory grammar by performing perturbation experiments. Grammar dependence differed markedly among cell types. Deleting individual motifs reduced activity in all three cell lines, whereas shuffling motif positions while preserving motif composition and background impaired activity in K562 and HepG2 but not in SK-N-SH (**Fig. 5e**). In K562 and HepG2, the strongest individual motif contributed only 31.1% and 14.2% of intact enhancer activity, respectively, while intact enhancers exhibited 3.1-fold and 5.0-fold higher activity than expected from independent motif action (**Fig. 5f**). These results show that REGAL designs derive activity not only from motif content but also from cooperative interactions among spatially arranged motifs.

By contrast, SK-N-SH enhancers did not significantly exceed the expectation from independent motifs, and TP53 alone contributed 87.6% of intact enhancer activity, consistent with a TP53-dominated architecture with weaker dependence on cooperative motif interactions (**Fig. 5f**). This TP53-dominated architecture of SK-N-SH designs correlated with experimentally observed leakiness in HepG2 (**Fig. 5d**) and prompted us to test whether REGAL’s explicit motif vocabulary could be exploited to diagnose and correct off-target activity. We therefore used oracle-based motif-family to identify dominant motif contributors, removed them from the motif vocabulary, and retrained the placement policy after each round (**Extended Data Fig. 8a**). After iterative vocabulary curation, the contribution of the leading TF family to predicted off-target activity decreased from 73.3% to 15.5%. When calibrated to the measured STARR-seq response scale (see Methods), HepG2 off-target activity decreased from log₂FC 2.48 to 0.60 while SK-N-SH activity was largely retained (5.29 to 4.61) (**Extended Data Fig. 8b**). Vocabulary curation also altered the specificity attribution: the single motif domination effect was largely eliminated and the strongest single motif contributed only 0.24 to the specificity score, whereas the complete enhancer achieved a score of 4.01, indicating that specificity emerged from combinatorial motif interactions (**Extended Data Fig. 8c,d**). Taken together, removing dominant motif families did not abolish enhancer design but redirected REGAL toward alternative combinatorial grammars, establishing the regulatory alphabet itself as a programmable and interpretable control point for cell-type-specific enhancer design.

### REGAL generalizes enhancer grammar design across vertebrate and invertebrate genomes

To test the portability of REGAL design across species, we next exchanged only the oracle and motif vocabulary for zebrafish and *Drosophila*, benchmarking against the state-of-the-art design framework for each system.

In zebrafish, we coupled REGAL’s placement policy to the DeepZebrafish single-cell chromatin accessibility oracle used by CREsted, and to a zebrafish TF motif vocabulary^42^. REGAL converged on tissue-specific regulatory grammars for cardiac muscle, endothelium, fast muscle, and slow muscle that recovered canonical lineage regulator motifs, including MEF2 and GATA for cardiac muscle, MYOG for muscle^43^, and FLI1 for endothelium (**Fig. 6a**)^44^. Scoring 200 designs per method across four cell types on the shared DeepZebrafish oracle, REGAL matched or exceeded CREsted designs in predicted on-target accessibility and specificity (**Fig. 6b,c**). On-target accessibility was higher in endothelium, fast and slow muscle (median predicted accessibility 31.0, 40.3 and 36.9 versus 17.7, 16.3 and 18.5) and comparable in cardiac muscle (19.1 versus 19.9), and specificity followed the same pattern. REGAL achieved this with substantially shorter enhancers than CREsted designs (192 bp versus 500 bp). We next tested nine REGAL-designed enhancers *in vivo* by transient reporter assays in zebrafish embryos (**Fig. 1c**), three each for cardiac muscle, somatic muscle (designed against the fast- and slow-muscle outputs) and notochord, an additional target for which no published designs are available for benchmarking. All nine reporters were specifically active in their target cell type in injected zebrafish embryos (9/9 constructs; **Fig. 6d,e**; **Extended Data Fig. 9a,b**; **Supplementary Data 6**).

**Fig. 6.**
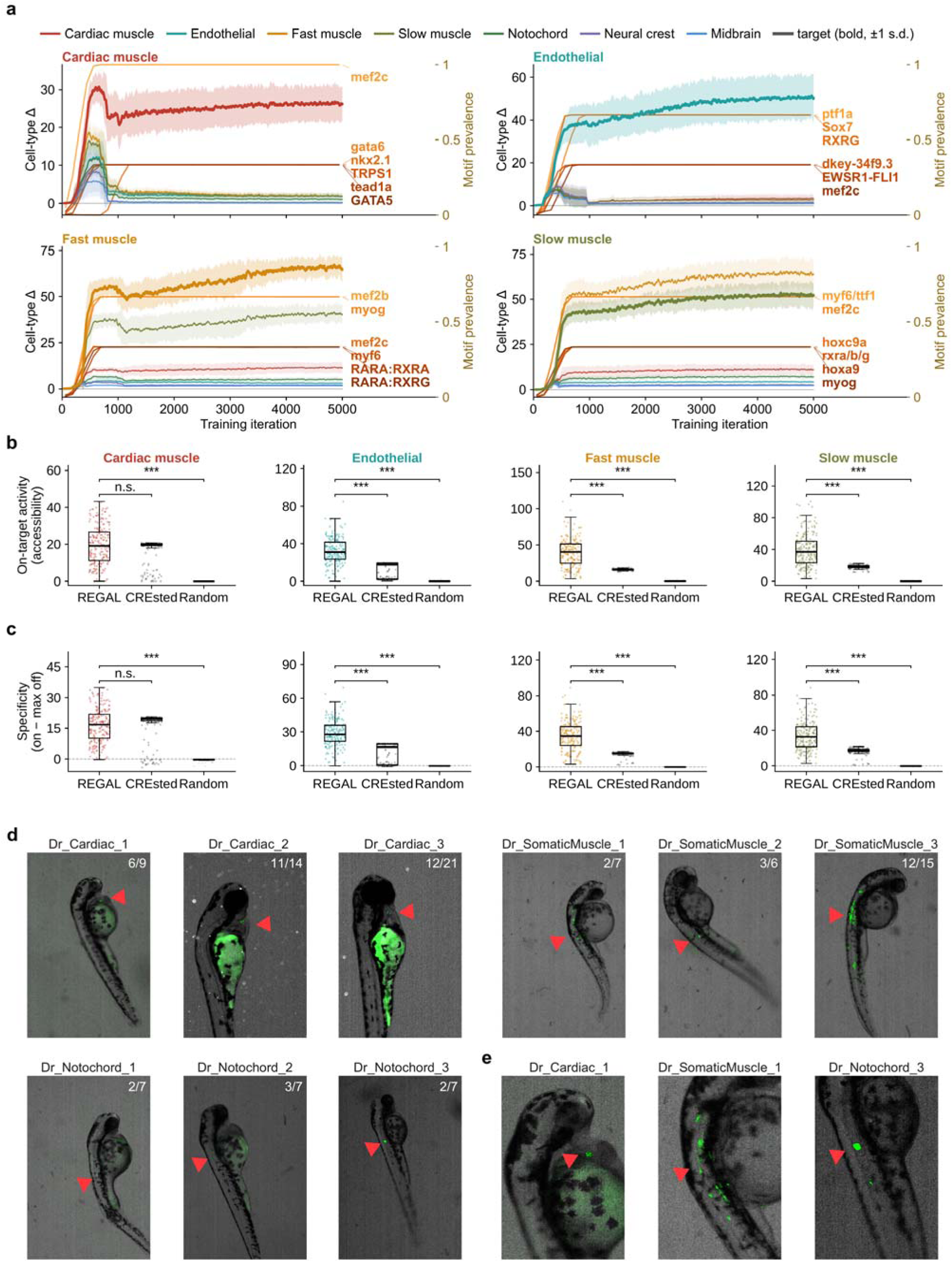
REGAL designs tissue-specific enhancers in zebrafish. **a**, Training for cardiac, endothelial, fast- and slow-muscle enhancer design using the DeepZebrafish oracle. Thick lines show the target-cell output and thin lines the off-target outputs (mean ± standard deviation across three independent seeds). Orange curves show the prevalence of representative motifs enriched during training. **b,c,** DeepZebrafish-predicted on-target accessibility (b) and specificity (c) of REGAL designs compared with CREsted designs and random sequences. Specificity was defined as target-cell accessibility minus the strongest off-target accessibility, with fast- and slow-muscle outputs excluded from one another’s off-target set. Boxes show the median and IQR, whiskers extend to 1.5 × IQR, and points represent individual sequences. Comparisons with REGAL were assessed using two-sided Mann–Whitney U tests; ***P < 0.001; n.s., not significant. **d,** Representative 48-hpf zebrafish embryos expressing a green fluorescent reporter driven by REGAL-designed enhancers. Red arrowheads indicate reporter expression in the intended target tissue, and numbers indicate the fraction of scored embryos showing target-tissue activity. Tiles are labeled Dr_<cell type>_<NUMBER> (for example, Dr_Cardiac_1; Dr, *Danio rerio*). **e,** Enlarged views for cardiac 1, somatic-muscle 1, and notochord 3. hpf, hours post-fertilization.

We then exchanged the oracle with DeepSTARR (embryo)^10^ and the motif vocabulary with a *Drosophila* motif collection (**Fig. 1c**). REGAL again converged on tissue-specific grammars enriched for canonical developmental regulators, including Serpent (srp) for gut^45^, Mef2^46^ together with the NK-homeobox factor Tinman (tin)^47^ for muscle, and Tramtrack (ttk) for CNS (**Extended Data Fig. 10a**). Benchmarked *in silico* using the shared DeepSTARR oracle, 192-bp REGAL designs (three per tissue) matched or exceeded the 501-bp DeepSTARR-designed enhancers in on-target activity and predicted tissue specificity across CNS, gut and muscle. Both design methods outperformed natural enhancers and random sequences (**Extended Data Figs. 9c,d and 10b,c; Supplementary Data 7**).

Together with the mouse and human results, these findings establish REGAL as a portable and species-agnostic framework for programmable design and engineering of compact, tissue-specific enhancers across vertebrate and invertebrate genomes.

## Discussion

Existing deep learning-based enhancer design has largely optimized nucleotide sequences^4,9–12^, leaving regulatory grammar as a latent property interpreted post hoc. REGAL instead makes regulatory grammar itself the substrate of design, representing each enhancer as an explicit combination of TF motifs and their spatial organization. This representation makes the design elements interpretable, experimentally testable and revisable, allowing functional measurements to be mapped back to specific grammatical choices. By separating the motif vocabulary, grammar-writing policy and the predictive oracle, REGAL provides a flexible and modular architecture that can be adapted across biological systems by exchanging regulatory alphabets and sequence-to-function models.

REGAL revealed that tissue specificity is a composite regulatory property. First, tissue specificity requires joint optimization of target-lineage activation and off-target repression. Second, enhancer function depends on both motif composition and spatial organization. Third, different lineages achieve specificity through distinct grammatical strategies: cardiac enhancers frequently exploit bifunctional motifs, whereas neural enhancers rely more on distributed motif composition and arrangement. Thus, tissue-specific enhancer function does not appear to arise from a universal grammar but from distinct combinations of activating, repressive, and cooperative interactions.

The effectiveness of REGAL further establishes regulatory grammar as a functional search space for enhancer design. Rather than accumulating individually active motifs, REGAL optimizes combinations of motif identity and organization to identify effective regulatory configurations. In K562 and HepG2, intact REGAL enhancers exceeded the activity expected from independent motif action. These results suggest that enhancer function depended on motif cooperativity and that grammar-level optimization can uncover combinatorial solutions that would be missed by selecting motifs on the basis of their individual effects.

Grammar-level design also showed strong prediction-to-validation fidelity. At matched predicted activity in K562 and HepG2, REGAL designs produced greater measured STARR-seq activity than nucleotide-level designs, indicating that predicted gains translated more effectively into experimental activity. This fidelity extended across experimental systems, with all selected designs tested in mouse and zebrafish embryos producing activity in their intended lineages. REGAL designs also showed substantially less CpG enrichment than nucleotide-level designs and remained closer to natural sequence composition for constraining biologically meaningful regulatory features. Importantly, these constraints remain editable: when a dominant TF family motif contributed to off-target activity in SK-N-SH designs, removing that motif family from the vocabulary redirected optimization toward alternative combinatorial grammars while largely preserving target activity and improving specificity. Thus, an explicit grammar provides a biologically structured design space that can improve both the reliability and controllability of enhancer design.

REGAL has several limitations. First, its performance depends on the reliability and resolution of the predictive oracle. For example, neural designs spanned all four CNS subregions, consistent with the oracle’s coarse predictions, and cell types or states not represented reliably by the oracle may remain out of reach by REGAL. Second, REGAL may capture only part of the regulatory grammar. Although our analyses support pairwise cooperativity and preferred spacing, higher-order motif interactions remain incompletely characterized, requiring extensive multiway perturbations. Third, motif-vocabulary coverage and fidelity bound performance. Although an explicit regulatory alphabet can be inspected, expanded, or restricted, motif redundancy and incomplete annotation are likely to constrain REGAL designs.

The compact size and tissue specificity of REGAL designs may be advantageous for therapeutic applications. At 192 bp, these enhancers preserve vector payload capacity^6^. REGAL’s contrast optimization penalizes undesired off-target activity, potentially reducing ectopic expression and associated toxicity. More broadly, because each design is represented as an explicit set of motifs and spatial organization, experimental measurements can be mapped back to specific grammatical features and used to refine subsequent designs, providing a route toward iterative design–build–test–learn cycles^48^. By making regulatory grammar rather than nucleotide sequence the design substrate, REGAL shifts enhancer engineering from sequence optimization toward programming regulatory logic, providing a framework to build, interrogate, and refine tissue-specific gene regulation.

## Methods

### 1. VISTA dataset and prediction task

We obtained developmental enhancer annotations from the VISTA Enhancer Browser^18^ (accessed September 11, 2024), which reports reproducible E11.5 lacZ activity in transgenic mouse embryos. The dataset comprised 3,955 unique mm10 sequences annotated across seven tissues: forebrain, midbrain, hindbrain, neural tube, heart, embryonic facial prominence (face), and limb. We retained elements identified by mouse or human comparative genomics. For each tissue, sequences scored as active were treated as positives and all assayed but inactive elements as tissue-matched negatives. Per-tissue annotation counts are provided in **Supplementary Table 1**. Prediction was formulated as seven independent binary classification tasks. Sequences were split into training, validation, and test sets at an 8:1:1 ratio, stratified to preserve tissue-specific activity distributions.

### 2. Sequence-to-function oracle

#### 2.1 Architecture benchmark

To select the architecture best suited to serve as the design oracle, we benchmarked six models spanning four architectural families: a 1D convolutional network (DeepSTARR)^8^, a convolutional neural network–transformer hybrid (Enformer)^7^, a BERT-style transformer with byte-pair-encoding tokenization (DNABERT-2)^16^, and three HyenaDNA long-convolution variants differing only in pretraining context (32K, 450K, 1M)^17^.

Each model was trained — fine-tuned from a pretrained checkpoint, or from randomly initialized weights in the case of DeepSTARR — and evaluated on the seven per-tissue binary classification tasks; pretrained-checkpoint sources and per-model configurations are given in **Supplementary Information**. All models were trained under a standardized protocol with a shared classification head (**Supplementary Information**). Discriminative performance was quantified as AUROC, with the area under the precision–recall curve reported as a secondary metric given the pronounced class imbalance, and was averaged over five random seeds per tissue and then across the seven tissues to give an overall mean AUROC.

#### 2.2 Enformer oracle fine-tuning

On the basis of the architecture benchmark, we selected Enformer as the design oracle and fine-tuned it at the 1,536-bp input length (12 × 128-bp output bins) used for downstream design and scoring. We trained one oracle per tissue, giving seven independent binary classifiers. Each oracle comprised the pretrained Enformer backbone fine-tuned end-to-end^7^ followed by a fixed classification head: Enformer trunk → adaptive average pooling → multilayer perceptron → two-class linear head. Weights were loaded strictly from per-tissue validation-best checkpoints. The tissue-activity score was the softmax probability of the active class for tissue t, evaluated on the full 1,536-bp composite sequence. Fine-tuning used AdamW (learning rate 1×10⁻⁴) with cosine annealing and warm restarts^49^. Validation, held-out test evaluation, and specified post hoc rescoring analyses averaged logits over the forward and reverse-complement strands before softmax (**Supplementary Information**). Per-tissue production-oracle AUROC is given in **Supplementary Table 3**.

#### 2.3 Genome-wide enhancer scanning

Each tissue oracle was applied genome-wide in 1,536-bp windows with a 200-bp step. Windows exceeding the tissue-specific classification threshold were merged into contiguous regions, which were ranked by their maximum predicted probability (**Supplementary Table 3**). The top 5,000 regions per tissue were retained as predicted enhancers (**Supplementary Data 8**), with each region represented in downstream analyses by its highest-scoring 1,536-bp window.

### 3. Oracle validation

The tissue-specific oracles were evaluated using three independent datasets not used for training: enhancer-associated chromatin states, in vivo saturation-mutagenesis effects, and lineage-matched ATAC-STARR-seq activity.

#### 3.1 ChromHMM chromatin-state enrichment

The top 5,000 predicted regions per tissue were intersected with matched E11.5 ENCODE ChromHMM annotations (**Supplementary Table 4**)^20^. Each region was assigned its predominant chromatin state, and enrichment of enhancer-associated states (EnhG, EnhPois, Enh, EnhLo, and EnhPr) was assessed relative to 5,000 randomly repositioned control regions using two-sided Fisher’s exact tests with Benjamini–Hochberg correction^50^.

#### 3.2 *In vivo* saturation mutagenesis

Oracle predictions were compared with an independent transgenic mouse saturation-mutagenesis dataset^19^ comprising 167 block substitutions across seven limb, heart, and neural enhancers. Each variant was scored with the oracle corresponding to its primary tissue, and the predicted mutational effect was defined as the change in enhancer probability relative to the wild-type sequence. Predicted effects were normalized within enhancer and compared with the measured in vivo functional scores by Pearson correlation and Kruskal–Wallis testing across functional categories.

#### 3.3 Enrichment in ATAC-STARR-seq active elements

The top 5,000 predicted regions per tissue were compared with endogenous ATAC-STARR-seq active elements from N2a, HL-1, and mESC. Cell-type-specific hits were defined as overlaps with active elements from one cell line but not the other two. Enrichment was assessed against chromosome- and GC-matched random genomic regions using two-sided Fisher’s exact tests with Benjamini–Hochberg correction. Lineage preference was further tested for neural predictions in N2a versus HL-1 and heart predictions in HL-1 versus N2a.

### 4. REGAL framework

REGAL designs tissue-specific enhancers by oracle-guided placement of TF motifs into a genomic background, optimized with a GRPO loop.

#### 4.1 Framework overview

REGAL consists of a fixed motif vocabulary, a trainable autoregressive Transformer placement policy, and a frozen sequence-to-function oracle. A separate frozen AlphaGenome encoder provides genomic-context embeddings to the policy. Only the placement policy is trained; all other components remain fixed. Candidate enhancers were generated within a 1,536-bp genomic background containing a central 192-bp design window. The policy sequentially selected motif identity, position, and orientation, with up to five motifs placed per sequence while preserving the surrounding genomic context. At each training iteration, genomic backgrounds and their precomputed AlphaGenome^51^ embeddings were sampled, the policy generated candidate motif configurations, and the resulting sequences were scored by the oracle. Rewards were calculated from changes in oracle output and used to update the placement policy by GRPO^15^. Campaign-specific vocabularies, oracles, background pools, targets, and scoring windows are summarized in **Supplementary Table 5**.

#### 4.2 Genomic context encoding with AlphaGenome embeddings

The genomic context of each background was encoded by a frozen AlphaGenome model^51^ used as a feature extractor. For each background, the 1,536-bp sequence was symmetrically N-padded to AlphaGenome’s 2,048-bp minimum input (256 bp per side) and one-hot encoded (A,C,G,T → standard basis; ambiguous bases → uniform 0.25). The model’s intermediate per-position representation, embeddings_1bp (1,536 channels), was read rather than the final accessibility output head; the 256-bp pad was then cropped from each side to recover a context feature map of shape [1,536, 1,536] (positions × features) aligned to the input.

#### 4.3 Motif placement policy network

The placement policy was a context-conditioned autoregressive Transformer decoder^14^ with six layers, eight attention heads, a model dimension of 768, and a feed-forward dimension of 3,072. The decoder attended to frozen AlphaGenome genomic-context features and sequentially generated up to five motif placements. At each step, the policy selected motif identity, position, and orientation through separate action heads. Position choices were conditioned on the selected motif and masked to prevent out-of-bounds or overlapping placements, and orientation was chosen as forward or reverse-complement. Each completed placement was appended to the decoding history, and generation terminated upon a STOP action or after five motifs.

#### 4.4 Reward design

For each candidate sequence, the oracle reward was defined from the change in predicted activity relative to its matched genomic background. The contrastive objective rewarded increased activity in the target tissue while penalizing activity in designated off-target tissues, with tissue-specific weights given in **Supplementary Table 6**. Additional regularization terms discouraged motif overlap and promoted motif-family diversity during training. Rewards were standardized and clipped before GRPO optimization.

#### 4.5 Policy optimization with GRPO

The policy was trained with GRPO^15^, implemented as a group-baseline REINFORCE objective^52^. Each iteration sampled batch_size = 16 background sequences and, for each, generated samples_per_prompt = 4 candidate configurations. The 64 rewards were reshaped into 16 groups of 4 sharing a background, and a group-relative advantage was computed within each group, 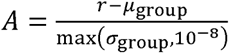 . The loss was 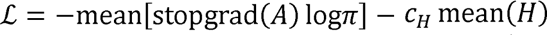, with the entropy coefficient c_H = 0.01 and H the per-candidate decoding entropy; log-probabilities and entropy came from a single on-policy forward pass. Sampling temperature was initialized at 1.5 and decayed multiplicatively by 0.995 each iteration, with a floor of 0.5. Parameters were optimized with AdamW^49^, with gradients clipped to a global norm and the learning rate following a warm-up and cosine-decay schedule. Policies were trained from scratch with a fixed random seed and deterministic cuDNN. Per-campaign iteration overrides and the seed lists are tabulated in **Supplementary Table 6**.

### 5. Design campaigns

The REGAL framework was instantiated for four design campaigns: mouse *in vivo* enhancers and three cross-species settings (*Drosophila*, human cell lines, and zebrafish). Each campaign fixes the same four swappable slots — motif vocabulary, reward oracle, background pool, and target set — together with its objective and a small set of optimizer overrides. These slots are tabulated in **Supplementary Table 5 and Supplementary Table 6**. Each oracle read a model-specific window of the composite sequence (**Supplementary Table 5**).

#### 5.1 Mouse (*in vivo*, VISTA)

Motif vocabulary was derived de novo by applying TF-MoDISco-lite (v2.3.2)^22^ to integrated gradients attribution scores from the seven tissue-specific Enformer oracles. This yielded 637 motifs, including 326 with positive and 311 with negative attribution. Each motif was represented by a position weight matrix (PWM) and rendered during design as its consensus sequence. Motifs were additionally annotated to TF identities and grouped into 28 TF families. The reward oracle was the per-tissue Enformer classifier, reading the full 1,536-bp composite (**Supplementary Table 5**). Backgrounds were real VISTA-negative^18^ sequences for the matching tissue. Two objectives were run for six target tissues—forebrain, midbrain, hindbrain, neural tube, heart, and limb—a single-tissue objective and a contrastive objective (**Supplementary Table 6**).

#### 5.2 Drosophila

Motifs were drawn from the jaspar_ffs_v1^53,54^. The reward oracle was DeepSTARR_embryo^10^, a PyTorch port numerically matched to the original Keras DeepSTARR model. A separate single-output convolutional model (four convolution–batch-norm–max-pool blocks, two dense layers, and a sigmoid activity head) was used per embryonic tissue, and each tissue score was the mean over an AUROC-filtered fold ensemble — cross-validation folds with AUROC ≥ 0.60 and NaN-rate ≤ 0.05. The oracle read only the central 1,001-bp window (**Supplementary Table 5**); within each fold the activity probability was standardized against that fold’s inactive-background statistics, and the fold z-scores were averaged to give tissue-activity score. Backgrounds were tissue-inactive DeepSTARR genomic tiles (1,001-bp cores padded to 1,536 bp with GC-matched flanks). The design targets were three embryonic tissues — CNS, gut, and muscle; brain and epidermis are oracle heads used as off-targets (**Supplementary Table 6**).

#### 5.3 Human cell lines

Motifs were drawn from the jaspar_vertebrates_2024^53^ (**Supplementary Table 5**). The reward oracle was Malinois^4^, loaded from the published model artifact. Malinois emits continuous massively parallel reporter assay (MPRA) log₂FC for K562, HepG2, and SK-N-SH simultaneously (output channels 0, 1 and 2, respectively). The oracle read the central 200-bp core, padded to the model’s 600-bp input with the fixed MPRA adapter flanks and scored as the mean of its forward and reverse-complement passes (**Supplementary Table 5**); tissue-activity score was the raw log₂FC for the target cell line. Backgrounds were low-activity MPRA oligos per cell line (|log₂FC| < 0.1 and standard error < 0.3) taken from the Gosai et al. MPRA library^4^, recentered and extended to 1,536 bp on GRCh38 with ENCODE-blacklisted^55^ and N-rich windows removed. One policy was trained per cell line, with the two other cell lines as mutually competing off-targets (**Supplementary Table 6**).

#### 5.4 Zebrafish

Motifs were drawn from the zebrafish_swissregulon_jaspar_merged_2026 vocabulary^53,56^ (**Supplementary Table 5**). The reward oracle was DeepZebrafish^42^, a multi-task accessibility model trained on zebrafish embryonic single-cell ATAC-seq data^57^. The objective was a 639-head contrastive reward, 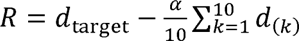, where 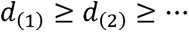 denote the deltas over the penalized off-target head pool sorted in descending order; “neutral” same-group heads were excluded from this pool — any head with a non-zero target vector, e.g. both the 48- and 72-h post-fertilization (hpf) versions of the target cell type and closely related subtypes. The training used α = 1.0 for endothelial and slow muscle, and α = 2.0 for cardiac muscle, fast muscle and notochord. α was held at 0 for iterations 1–500, linearly ramped to its final value between iterations 500 and 1,000, and held thereafter (**Supplementary Table 6**). Backgrounds were per-cell-type, per-timepoint 1,536-bp pools. Five cell types entered the reported analyses — cardiac muscle, endothelial, fast muscle, slow muscle, and notochord (**Supplementary Table 6**).

### 6. Experimental validation

#### 6.1 Cell culture

Mouse enhancer validation used E14Tg2a mESCs (ATCC CRL-1821), Neuro-2a cells (N2a; ATCC CCL-131), and HL-1 cardiomyocytes (Sigma-Aldrich SCC065). Human validation used HepG2 (ATCC HB-8065), K562 (ATCC CCL-243), and SK-N-SH (ATCC HTB-11) cells. Cells were maintained at 37 °C and 5% CO₂ under supplier-recommended conditions. mESCs were cultured on 0.2% gelatin-coated plates in Knockout DMEM supplemented with 10% ESC-qualified fetal bovine serum (FBS), 1 mM GlutaMAX, 0.1 mM MEM non-essential amino acids, 0.1 mM 2-mercaptoethanol, penicillin–streptomycin, leukemia inhibitory factor, and the 2i inhibitors CHIR99021 and PD0325901, and were passaged every 2 days. N2a cells were maintained in EMEM with 10% FBS and penicillin–streptomycin and passaged every 2–3 days at 80–90% confluency. HL-1 cardiomyocytes were cultured on 0.02% gelatin-coated flasks in Claycomb Medium supplemented with 10% FBS, 0.1 mM norepinephrine, 2 mM L-glutamine, and penicillin–streptomycin, with daily medium changes. HepG2 and SK-N-SH cells were maintained in EMEM with 10% FBS and penicillin–streptomycin and passaged every 3–4 days at 70–90% confluency. K562 cells were cultured in RPMI-1640 with 10% FBS and penicillin–streptomycin and maintained at 1 × 10⁵–1 × 10⁶ cells ml⁻¹.

#### 6.2 Endogenous ATAC-STARR-seq enhancer maps

Whole-genome ATAC-STARR-seq^33^ was performed in N2a, HL-1, and mESC with three paired input-DNA and reporter-RNA replicates per cell line. Libraries were sequenced paired-end on an Illumina NovaSeq X Plus and base calls were demultiplexed per library with bcl-convert (v4.2.4). Reads were adapter- and quality-trimmed with Trim Galore^58^ (v0.6.10; Cutadapt v4.9^59^, default parameters) and aligned with Bowtie2^60^ (v2.3.4.1; --sensitive -X 500 --no-discordant --no-mixed). Alignments were filtered to mapping quality ≥ 30, and mitochondrial reads and ENCODE mm10 blacklist (v2)^55^ regions were removed (SAMtools v1.14^61^). Input (DNA) reads were deduplicated with Picard MarkDuplicates (v2.18.14)^62^, whereas reporter (RNA) reads were retained without deduplication to preserve transcript abundance. Reporter activity was determined with DESeq2^63^ by comparing RNA with input DNA. Regions with false discovery rate ≤ 0.1 and positive log₂(RNA/DNA) were classified as active, whereas significantly depleted regions were classified as silent. Adjacent significant regions were merged. Counts-per-million-normalized input and reporter coverage tracks were generated with deepTools^64^.

#### 6.3 STARR-seq shared reporter-assay protocol

Synthetic oligo pools or endogenous accessible fragments were cloned into the hSTARR-seq_ORI^34,65^ vector and propagated in MegaX DH10B cells. Plasmid libraries were introduced into the corresponding cell lines by Neon electroporation, and cells were harvested 24 h later for paired reporter-RNA and input-plasmid-DNA isolation. Poly(A)+ RNA was purified, DNase-treated, and reverse-transcribed using a reporter-specific primer. Reporter cDNA and input DNA were amplified with indexed sequencing primers. Oligonucleotides used for library preparation are listed in **Supplementary Table 7**.

#### 6.4 Designed STARR-seq library

Designed enhancers were synthesized as 192-bp inserts flanked by cloning adapters. REGAL designs were used directly, whereas 200-bp CODA and natural DHS sequences were trimmed symmetrically to 192 bp to meet synthesis constraints. Re-scoring confirmed that this trimming preserved CODA predictions (Pearson r = 0.971–0.992 across the three human cell lines).

Mouse libraries were assayed in N2a and HL-1 cells, and the pooled human library in K562, HepG2, and SK-N-SH cells, with three replicates per cell line. Cell-line-specific Neon electroporation settings are provided in **Supplementary Table 7**.

Sequencing reads were aligned directly to the designed-oligo library and counted per oligo. Reporter activity was estimated with DESeq2 from paired RNA and DNA counts, with low-count oligos excluded. Size factors were estimated from designed-null controls to anchor log₂FC = 0 to the inactive baseline, and activity was tested against a 1.5-fold effect-size threshold. Oligos with adjusted P ≤ 0.05 and log₂FC > 0 were classified as active. For mouse designs, N2a served as the on-target line for neural enhancers and HL-1 for cardiac enhancers; selectivity was assessed from on-versus off-target activity. For human designs, specificity was defined as the cognate-cell log₂FC minus the maximum off-target log₂FC. Replicate concordance is reported in **Supplementary Tables 8–10**.

#### 6.5 Perturbation sublibrary design (LOO and position-shuffle)

Both series were built on a shared set of the top contrastive designs — the 50 most active GRPO_contrastive enhancers per tissue (midbrain, hindbrain, neural tube, heart). For the LOO series, each variant was generated by deleting one motif from the intact design while leaving all other motifs unchanged. For the position-shuffled variants, each parent generated ten variants in which all five motif identities were retained but their positions and strands were randomly reassigned.

#### 6.6 Transgenic mouse validation (enSERT)

Designs from the mouse campaign were ranked by the Enformer oracle prediction logit of the target tissue. Nine designs randomly drawn from the top 5% of their target-tissue pools were carried forward to transgenic testing: three targeting heart, three targeting limb, and one targeting each of midbrain, hindbrain and forebrain **Supplementary Data 4**.

Candidate enhancer sequences were synthesized by Twist Bioscience and cloned into a donor plasmid containing a minimal Shh promoter, a lacZ reporter, and H11-locus homology arms (Addgene, 139098) using NEBuilder HiFi DNA Assembly Master Mix (New England Biolabs, E2621)^35^. Donor plasmid sequence identity was verified by long-read sequencing.

Transgenic embryos were generated by the Case Transgenic and Targeting Facility (Case Western Reserve University) using site-directed transgene insertion at the H11 safe-harbor locus (enSERT)^35,36^. FVB males were bred with three-week-old superovulated FVB females to generate fertilized eggs. Lyophilized Cas9 protein (PNAbio) was resuspended in nuclease-free water at 1 µg µl⁻¹. The H11-targeting single-guide RNA (sgRNA, Synthego; spacer 5′-GCTGATGGAACAGGTAACAA-3′) was resuspended at 1 µg µl⁻¹ in nuclease-free 10 mM Tris-HCl (pH 8.0). An injection mixture of 80 ng µl⁻¹ Cas9 protein, 80 ng µl⁻¹ H11 sgRNA and 10 ng µl⁻¹ supercoiled donor plasmid in nuclease-free water was injected into the pronucleus of FVB fertilized eggs. Injected embryos were cultured in droplets of EmbryoMax Advanced KSOM (Millipore) under mineral oil in benchtop MINC incubators under a tri-gas atmosphere (5% CO₂, 5% O₂, 90% N₂) and transferred by oviduct transfer into pseudopregnant CD-1 recipients. The day after oviduct transfer was designated E1.5. Pregnant dams were euthanized at E11.5; uteri were dissected into phosphate-buffered saline (PBS) and individual embryos recovered.

Embryos were processed for lacZ staining. Briefly, embryos were washed in cold PBS, fixed in 4% paraformaldehyde for 30 min and washed three times in embryo wash buffer (2 mM MgCl₂, 0.02% NP-40, 0.01% sodium deoxycholate in PBS, pH 7.3). Embryos were stained overnight at room temperature in X-gal staining solution (4 mM potassium ferricyanide, 4 mM potassium ferrocyanide, 1 mg ml⁻¹ X-gal, 20 mM Tris pH 7.5 in embryo wash buffer) and washed three times in PBS. Genomic DNA was extracted from yolk sacs using DirectPCR lysis reagent (Viagen, 301-C) supplemented with proteinase K (final concentration 6 U ml⁻¹) and used to confirm site-directed integration of the transgene at the H11 locus by junction PCR (**Supplementary Table 7**). Two independent junction assays were performed: a 5′-junction PCR, with a forward primer annealing in H11 genomic sequence 5′ of the left homology arm and a reverse primer in the Shh promoter, and a 3′-junction PCR spanning the transgene–genome boundary at the right homology arm. Both reactions were performed with VeriFi polymerase (PCR Biosystems) following the manufacturer’s standard two-step protocol (35 cycles). Each amplicon spans a junction present only in the correctly targeted allele, and an embryo was scored as positive for H11 integration when either the 5′-or the 3′-junction PCR yielded a band of the expected size. All reported analyses used the 186 PCR-positive embryos. A further 22 embryos showed majority-supported target staining without a confirmed junction-PCR band, consistent with non-targeted integration, and 65 embryos showed neither; both groups were excluded. Stained embryos were imaged on a Zeiss Stemi 305 stereomicroscope equipped with a Zeiss Axiocam 208 color camera.

X-gal-stained embryos were scored blind to construct identity. For each embryo, a single standardized whole-mount image (left-lateral view; right-lateral or dorsal substituted when a left-lateral view was unavailable) was white-balance-normalized, orientation-standardized (head-up) and stripped of identifying metadata, then given a randomized sample code. The resulting anonymized set of 273 images (nine designs, 21–41 embryos each) was scored independently by three assessors. Each assessor rated X-gal signal intensity on a 0–3 ordinal scale (0, no detectable staining; 3, maximal) for each of 15 anatomical structures — the seven oracle-modeled tissues (forebrain, midbrain, hindbrain, neural tube, heart, face, and limb) and eight additional structures scored but not modeled (dorsal root ganglion, cranial nerve, trigeminal, branchial arch, nose, eye, ear, and tail); structures without visible staining were left unmarked and treated as 0. A structure was positive when at least two assessors assigned a non-zero score; lineage-off-target staining was defined outside the intended lineage, with all four CNS structures treated as on-lineage for neural designs. The profile-level analysis used the same 186 embryos; consensus was computed per embryo across assessors and then across embryos.

#### 6.7 Zebrafish enhancer reporter assays

Designs from the zebrafish campaign were ranked by the DeepZebrafish-predicted accessibility of the target cell type, scored as defined in that section. Nine designs randomly drawn from the top 5% of their target-cell-type pools were carried forward to transient reporter testing: three targeting cardiac muscle, three targeting notochord, and three targeting the somatic-muscle category (**Supplementary Data 6)**.

Zebrafish enhancer-reporter constructs were built in the Tol2-I-SceI-ZSP:EGFP;cryaa:mCherry-I-SceI-Tol2 backbone^66^ (Addgene plasmid #194518; http://n2t.net/addgene:194518; RRID:Addgene_194518). Designed enhancer sequences were cloned upstream of the ZSP minimal promoter driving enhanced green fluorescent protein (EGFP); the lens-specific cryaa:mCherry cassette served as a transgenesis marker. Nine enhancer–reporter constructs were generated (three designed 192-bp synthetic enhancers each for cardiac muscle, notochord and somatic muscle) in the backbone Tol2-I-SceI-ZSP:EGFP;cryaa:mCherry-I-SceI-Tol2 (Addgene, 194518; 7,208 bp)^66^. Each enhancer was ordered as a full double-stranded gene fragment (Twist Bioscience) flanked by 30-bp homology arms, so no PCR amplification was required; each 433-bp fragment comprised an XhoI-site homology arm, the 192-bp enhancer, the 181-bp zebrafish synthetic promoter minimal-promoter body (restored on the fragment) and an AfeI-site homology arm. The vector was linearized in rCutSmart buffer (New England Biolabs) at 37 °C with XhoI + AfeI, gel-purified (NucleoSpin Gel & PCR clean-up, Macherey-Nagel), and assembled with the fragment at a 1:2 molar ratio using NEBuilder HiFi (New England Biolabs; 50 °C, 60 min). 2.5 µl of the reaction was transformed into DH5α chemically competent cells (30 min on ice, 45 s heat shock at 42 °C, 2 min on ice, SOC recovery) and plated on ampicillin. Single colonies were grown in 10 ml LB and plasmids isolated with the QIAprep Spin Miniprep Kit (Qiagen). All plasmids were verified by whole-plasmid long-read sequencing (Azenta/Genewiz); a sequence-perfect clone (expected size 7,362 bp) was recovered for all nine designs.

Plasmid DNA and Tol2 transposase mRNA were co-injected (at 30 and 100 ng µl⁻¹, respectively) into one-cell-stage embryos of the wild-type AB strain, which were then raised at 28.5 °C until 48 hpf^67^. Successfully injected embryos were selected on the basis of red fluorescence in the eye. At 48 hpf, embryos were anesthetized with 0.02% tricaine (MS-222; ethyl 3-aminobenzoate methanesulfonate), mounted in 3% low-melting-point agarose (Invitrogen) in a FluoroDish (FD3510-100, World Precision Instruments), and imaged for EGFP fluorescence on a Leica M165 FC fluorescence stereomicroscope. Images were processed in Fiji; brightness and contrast were adjusted linearly and applied identically across all embryos of a given construct. All zebrafish breeding was approved by the Institutional Animal Care and Use Committee (IACUC) of the Cleveland Clinic (protocol 2812), and all experiments were performed on embryos younger than 3 days post-fertilization.

For each enhancer, reporter assays were performed in multiple embryos across three replicates (that is, different injection days and clutches). Cell-type-specific reporter-expression patterns were qualitatively assessed from whole-embryo fluorescence images by a single experienced zebrafish investigator (T.F.S.). Only the cell-type specificity of each enhancer was assessed, not the level of activity, which cannot be quantified because the assay does not control for enhancer-reporter copy number per cell (episomal or integrated).

### 7. Downstream computational analyses

#### 7.1 Motif annotation and clustering

The de novo mouse motif vocabulary was assigned putative TF identities by matching each PWM against a merged reference database with Tomtom^23^ (MEME Suite v5.4.1^68^). The reference database unified 5,305 PWMs from three sources: HOCOMOCO v13 (H13CORE; 1,611 matrices)^69^, a published pairwise TF-combination PWM collection (1,348 matrices)^24^, and the JASPAR CORE collection^53^ (latest version per matrix, all taxa; 2,346 matrices, retrieved via pyjaspar^70^).

#### 7.2 Single-tissue versus contrastive enrichment

Motif usage was compared between the single-tissue (GRPO_diversity) and contrastive (GRPO_contrastive) design arms using the recorded motif placements, and motif function was evaluated using the LOO STARR-seq series.

**Placed-motif enrichment.** For each design, placed motifs were mapped to standardized TF names, and each TF was counted at most once per design. The analysis included 4,000 single-tissue and 4,000 contrastive designs across midbrain, hindbrain, neural tube, and heart. Differences in motif prevalence between the two design arms were tested using two-sided Fisher’s exact tests within each tissue and in the pooled neural set. For genome-wide comparisons shown in the volcano plots, all observed TFs were tested and P values were adjusted by the Benjamini–Hochberg procedure. Effect size was defined as the difference in placement frequency between contrastive and single-tissue designs.

**Per-motif bifunctionality.** Motif function was quantified from the measured change in STARR-seq activity after individual motif deletion from contrastive designs. For each motif, the mean LOO effect was calculated separately in N2a and HL-1 cells. Motifs were classified as activators when deletion reduced activity (mean Δlog₂FC ≤ −0.2), silencers when deletion increased activity (mean Δlog₂FC ≥ 0.2), and neutral otherwise. Neural motifs were evaluated with N2a as the on-target and HL-1 as the off-target cell line, with the assignment reversed for cardiac motifs. Motifs represented by at least three LOO measurements in each cell line were classified as dual-function when they acted as an on-target activator and an off-target silencer.

#### 7.3 Motif cooperativity and spacing

To test whether REGAL learned motif pairings and spacing preferences encoded by the Malinois oracle, we analyzed motif pairs recovered from converged human-cell-line policies.

**Pairwise synergy.** Frequently co-placed motif pairs were compared with randomly paired motifs using position-controlled second-order epistasis, calculated from pair and single-motif oracle scores at matched positions. Synergy was evaluated across multiple spacings and strand configurations and summarized by the maximum epistatic effect.

**Distance preference.** For each co-placed pair, one motif was fixed at its REGAL-learned position while the partner was scanned across feasible edge-to-edge distances and strand orientations. The oracle-preferred spacing was defined as the distance that maximized position-controlled reward epistasis and was compared with the spacing observed in REGAL designs.

**Spacing recovery.** To test whether these spacing preferences could be learned de novo, we trained policies constrained to place a single predefined motif pair while optimizing only position and orientation under the same Malinois reward. The learned spacing was taken from the converged policy and compared with the oracle-optimal spacing using Pearson correlation and absolute gap error.

#### 7.4 *In silico* benchmarking (Malinois /CREsted / DeepSTARR)

**Human cell lines.** REGAL designs for K562, HepG2, and SK-N-SH were compared with CODA^4^ designs (500 per cell line) and GO-CRE^38^ designs, with natural DHS elements and randomized sequences included as references. REGAL, CODA, DHS, and control sequences were scored using the same Malinois oracle and scoring procedure used during REGAL training; GO-CRE designs were evaluated using their released Malinois scores. Because Malinois outputs are directly comparable across cell lines, activity and specificity were calculated directly from the raw predicted MPRA log₂FC values.

**Zebrafish.** REGAL was compared with the 200 CREsted^42^ *in silico*-evolved enhancers available for each of four cell types (cardiac muscle, endothelial, fast muscle, and slow muscle), using 200 REGAL designs per cell type. All sequences were rescored with the DeepZebrafish oracle using its native input format. Activity was the target-cell accessibility score, and specificity was calculated against the remaining off-target heads using the same exclusions defined for the design campaign.

***Drosophila.*** REGAL designs targeting CNS, gut, and muscle were compared with the corresponding synthetic enhancers reported by de Almeida et al.^10^, together with natural active enhancers and random sequences as reference sets. All sequences were rescored with the same DeepSTARR embryo oracle ensemble used for REGAL design. Activity and specificity were calculated from background-normalized oracle scores, with specificity defined relative to the strongest off-target tissue.

#### 7.5 Sequence composition of designed enhancers

Sequence composition was summarized as CpG observed/expected, n(CpG) × L / (n(C) × n(G)). Each value was referred to natural DHS sequences of the same GC content^4^ per cell line using 1-percentage-point GC bins widened symmetrically until they held at least 200 reference loci. The reported statistic is the share of a collection above the 95th percentile of its GC-matched reference, with Wilson 95% confidence intervals.

#### 7.6 Prediction-to-measurement comparison at matched predicted activity

Arms were stratified on 2-log₂FC-wide bins of predicted on-target activity; a bin entered the test when both arms held at least eight oligos, and the arms were contrasted by a van Elteren stratified Wilcoxon test with weights inversely proportional to bin size, 1/(n + 1). Natural DHS elements were carried through the identical procedure as a natural-sequence reference. SK-N-SH was not evaluated because its arms share only one qualifying bin.

#### 7.7 Iterative motif-vocabulary curation

Every per-run quantity below was measured over the final 100 training iterations. These readouts are in silico. Each round’s predicted off-target signal was attributed to TF families as a first-order product of usage and effect: for every motif, its share of placed slots multiplied by its single-motif HepG2 delta, negative values set to zero, summed within families and normalized to 100%. Single-motif deltas came from a selectivity map in which each vocabulary motif was placed alone at the center of the design region across 16 fixed backgrounds. The leading family was withdrawn from the vocabulary, the policy retrained from scratch under the otherwise unchanged configuration with three seeds (42, 123, and 2024), and the attribution recomputed on the new policy, giving 15 runs each summarized by its mean oracle delta. Five rounds took the vocabulary from 879 motifs to 876 (− p53, 3 motifs), 874 (− ETS::SREBF2, 2), 865 (− IRF, 9), and 820 (− AP-1, 45). Predicted scores were placed on the measured scale by look-up rather than by a fitted model: the mean predicted score of the background panel (0.33 for SK-N-SH, 0.38 for HepG2) converted a training delta into an absolute predicted score, which was interpolated into the binned dose–response of this study’s human STARR-seq library.

### 8. Statistics, ethics, and availability

#### 8.1 Computational resources

REGAL optimization and design were performed on single NVIDIA H100 GPUs using Python 3.10.13 and PyTorch 2.9.1 (CUDA 12.8). A standard 1,500-iteration policy run required approximately 0.96 GPU-hours and 60 GB peak GPU memory. Reproducible software environments are provided with the code.

#### 8.2 Statistics and reproducibility

Unless otherwise stated, statistical tests were two-sided, with the test, sample size and exact P value reported in the corresponding figure legend. Fisher’s exact tests were used for enrichment and categorical comparisons, DESeq2 Wald tests for reporter activity, Mann–Whitney U or Wilcoxon signed-rank tests for unpaired or paired group comparisons, respectively, and Pearson or Spearman correlations for association analyses as appropriate. Prediction-matched activity was compared using van Elteren stratified Wilcoxon tests. Multiple-testing correction, when applied, used the Benjamini–Hochberg procedure^50^.

Endogenous and designed ATAC-STARR-seq experiments were performed in three biological replicates per cell line. Design policies were trained from scratch with fixed random seeds; campaign-specific settings are provided in **Supplementary Table 6**. Mouse embryos were scored independently by three assessors blinded to construct identity, and zebrafish reporter patterns were evaluated across three independent injection experiments. Statistical analyses were performed in Python and R.

#### 8.3 Ethics statement

All animal experiments were approved by the relevant institutional IACUCs. Mouse experiments were conducted at Case Western Reserve University under protocol 2014-0081 using FVB embryos and CD-1 pseudopregnant recipients; embryos were collected at E11.5. Zebrafish experiments were approved by the Cleveland Clinic IACUC under protocol 2812 and used wild-type AB embryos younger than 3 days post-fertilization. Recombinant and synthetic nucleic acid work was approved by the Cleveland Clinic Institutional Biosafety Committee (IBC 2205) in accordance with NIH guidelines. Sample sizes followed established field standards and were not predetermined statistically.

#### 8.4 Data availability

Sequencing data from the ATAC-STARR-seq and STARR-seq experiments have been deposited in the Gene Expression Omnibus under accession number GSE338744; the record is private pending publication and a secure reviewer token is provided with this submission. VISTA Enhancer Browser data are available at https://enhancer.lbl.gov, and the ENCODE Portal accessions for the E11.5 mouse ChromHMM segmentations are listed in **Supplementary Table 4**. Previously published enhancer-design and reporter datasets reanalyzed here were obtained from refs^4,10,19,38,42^. Designed enhancer sequences and metadata are provided as **Supplementary Data 2–7** and the de novo mouse motif vocabulary as **Supplementary Data 1**; the human, *Drosophila*, and zebrafish vocabularies were assembled from public motif databases as specified in **Supplementary Table 5**. X-gal-stained embryo and zebrafish reporter images are available at Zenodo (https://doi.org/10.5281/zenodo.21925412). Source data are provided with this paper.

#### 8.5 Code availability

The code implementing REGAL will be released upon publication under a non-commercial academic license.

#### 8.6 Use of large language models

Large language models (Claude Opus; Anthropic) were used to assist with language editing to improve clarity and organization of the manuscript, as well as generating analysis and visualization code. All language model-assisted text and code were reviewed, edited, and verified by the authors, who take full responsibility for the content of this manuscript. All scientific concepts, analysis, interpretations, and conclusions were developed and verified by the authors.

## Supporting information

Supplementary_Data_1_mouse_motif_vocabulary

Supplementary_Data_2_invitro_STARR_oligo_pool

Supplementary_Data_3_ablation_oligo_pool

Supplementary_Data_4_invivo_mouse_enhancers

Supplementary_Data_5_human_STARR_oligo_pool

Supplementary_Data_6_zebrafish_invivo_enhancers

Supplementary_Data_7_drosophila_designs

Supplementary_Data_8_genome_wide_predicted_enhancers

## Author contributions

Liangqi Xie conceived the original research direction, supervised the study, and provided resources. Liangqi Xie and L.Y. acquired funding. L.Y. developed and validated the initial enhancer-prediction framework, then conceived and developed REGAL, designed computational methods, performed computational and statistical analysis, designed and performed the ATAC-STARR-seq, STARR-seq, and motif-perturbation experiments, and designed and generated enhancers and reporter constructs for *in vivo* mouse and zebrafish validation. R.A.C. oversaw generation of the transgenic mouse embryos. L.Y. recovered the mouse embryos and performed fixation, X-gal staining and imaging, and prepared re-coded, anonymized image sets for blinded scoring. M.N.T., S.B. and one additional assessor (see Acknowledgements) scored the images independently, each blinded to construct identity. M.N.T. also assisted with mouse embryo junction-PCR genotyping and STARR-seq library preparation. S.B. also assisted with STARR-seq library preparation. A.S.R. and T.F.S. performed zebrafish husbandry and embryo microinjection. L.Y. performed zebrafish imaging and image processing, and T.F.S. scored the zebrafish reporter-expression patterns. Le Xiong and A.J. contributed to experimental design and data interpretation. L.Y. prepared the figures and wrote the original manuscript. Liangqi Xie reviewed and edited the manuscript. L.Y. revised the manuscript with input from all authors. All authors approved the final manuscript.

## Acknowledgements

We thank Drs. Feixiong Cheng, Ming Hu, John Barnard, Michelle S. Longworth, Len A. Pennacchio, Axel Visel, and Zhenghe J. Wang for scientific discussions and input. We thank Sohum Shah for independent blinded scoring of X-gal-stained embryo images. We thank David LePage, Andrea Trumbauer and Rachel Mann of the Case Transgenic and Targeting Facility for generating transgenic mouse embryos. We thank Alexey Chizhik for assistance with figure illustration. We also thank the VISTA Enhancer Browser and the Lawrence Berkeley National Laboratory for making the enhancer annotation resource publicly available, and members of the Xie laboratory for helpful discussions. Computational resources were provided by an allocation from the Ohio Supercomputer Center. L.Y. thanks Dr. L.Z. for her steadfast support and encouragement from across the ocean.

## Additional information

Supplementary Information is available for this paper. Correspondence and requests for materials should be addressed to Liangqi Xie.

## Extended Data Figures

**Extended Data Fig. 1.**
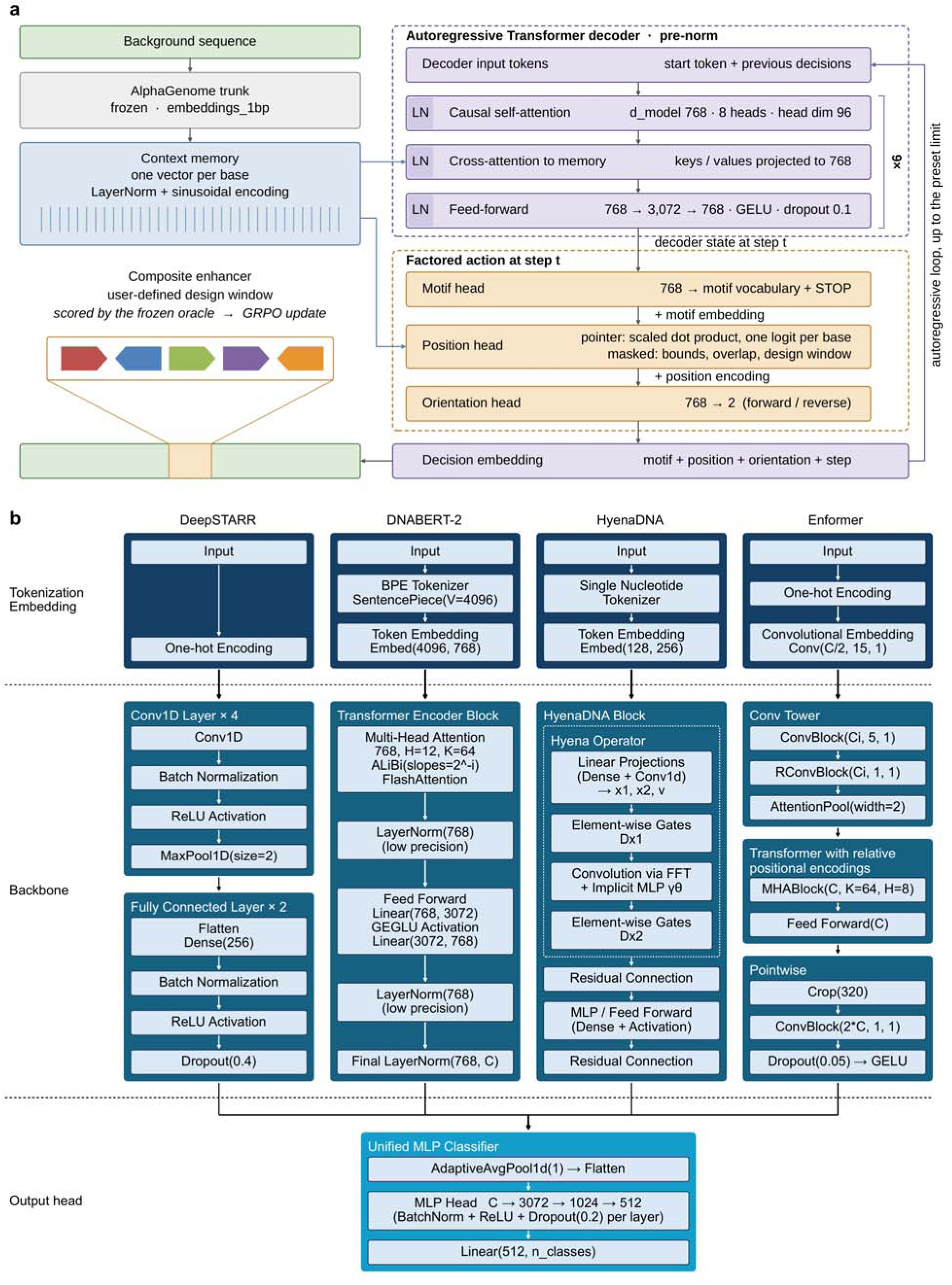
REGAL placement policy and design-oracle architectures. **a**, Architecture of the REGAL placement policy. A frozen AlphaGenome trunk converts the background sequence into base-resolution context features. A six-layer autoregressive Transformer cross-attends to these features and sequentially predicts motif identity, position, and orientation. Invalid, overlapping, or out-of-window positions are masked. The completed motif grammar is embedded in the user-defined design window and scored by the frozen oracle to provide rewards for GRPO. Colored arrowheads denote placements. **b,** Schematic of the four oracle architectures benchmarked for mouse enhancer prediction. DeepSTARR uses one-hot encoding and convolution; DNABERT-2 uses byte-pair encoding (BPE) and a Transformer backbone; HyenaDNA uses single-nucleotide tokens and Hyena blocks; and Enformer uses one-hot encoding, convolution, and a relative-position Transformer. Separately instantiated classifiers share one multilayer perceptron (MLP) architecture; DeepSTARR alone receives a flattened convolutional map. LN, layer normalization; GELU, Gaussian error linear unit; ReLU, rectified linear unit; MHA, multi-head attention; H, number of attention heads; K, key dimension; V, tokenizer vocabulary size; ALiBi, attention with linear biases; GEGLU, gated linear unit with GELU; FFT, fast Fourier transform; C, backbone feature width; n_classes, two activity classes.

**Extended Data Fig. 2.**
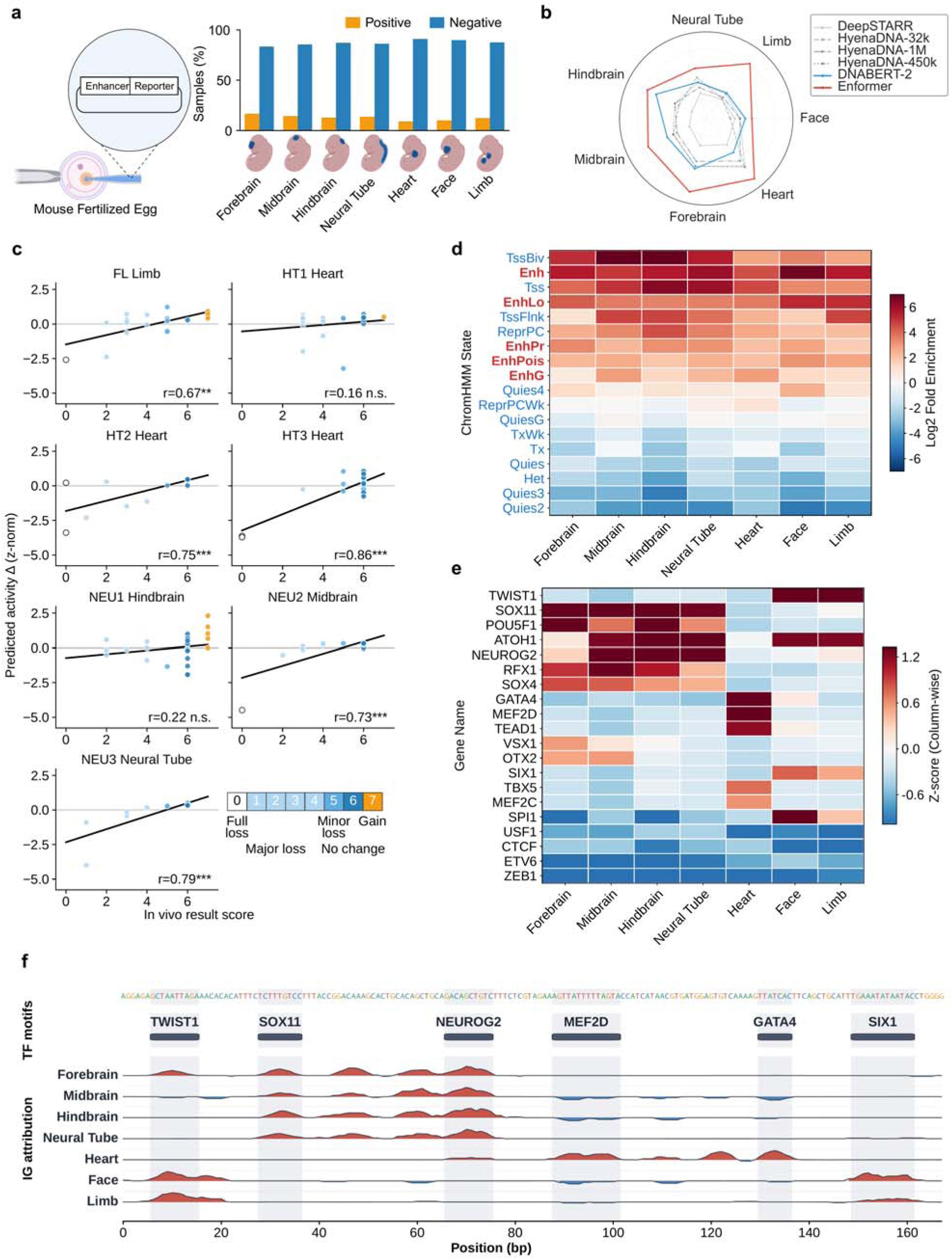
Benchmarking, validation and interpretation of VISTA enhancer oracles. **a**, Schematic of the VISTA reporter assay. Among 3,955 sequences, the numbers (n) scored positive for forebrain, midbrain, hindbrain, neural tube, heart, face, and limb were 658, 574, 507, 543, 353, 402 and 494, respectively (gold, positive; blue, negative). **b,** AUROC for six sequence-to-function oracles; vertices, five-seed means per model–tissue pair. **c,** Agreement between Enformer-predicted mutational effects and *in vivo* saturation-mutagenesis outcomes for 167 variants across seven enhancers. Black lines, least-squares fits; colors, labeled effect categories. *P < 0.05, **P < 0.01, ***P < 0.001; n.s., not significant. **d,** ChromHMM-state enrichment of the top 5,000 Enformer-predicted regions per tissue relative to random genomic regions. Values show log₂ fold enrichment; enhancer-associated states are highlighted in red. **e,** Column-standardized motif-attribution z-scores for the top 20 TF genes (140 values), with colors clipped at the 10th–90th percentiles. **f,** Integrated-gradients profiles across a representative motif-composed sequence (red, positive; blue, negative). Each track is normalized to its 99th absolute-signal percentile. ChromHMM, chromatin-state hidden Markov model; FL, limb element; HT1–HT3, heart elements; NEU1–NEU3, neural elements.

**Extended Data Fig. 3.**
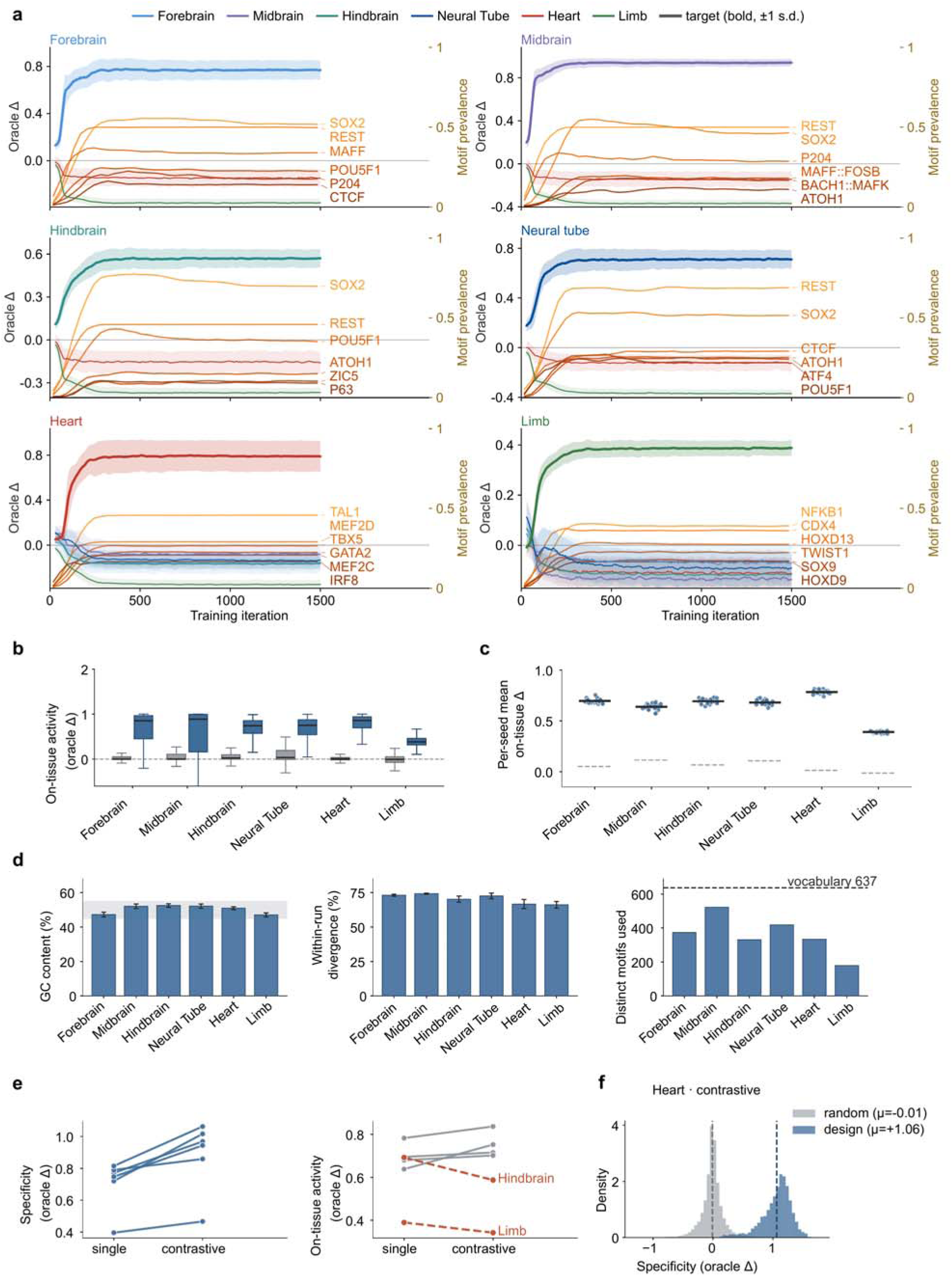
Convergence, reproducibility, and sequence quality of REGAL designs. **a**, Contrastive-training trajectories for six tissues. Lines show the mean oracle Δ, defined as the change in predicted activity relative to the matched background, across 24 independent seeds; target-tissue trajectories are shown as thick lines and off-target tissues as thin lines. Orange curves show the prevalence of representative motifs enriched during training. **b,** On-tissue oracle Δ for REGAL designs compared with motif-count-matched random sequences. Boxes show the median and IQR; whiskers extend to 1.5 × IQR. **c,** Reproducibility across 24 independently trained policies per tissue, shown as the mean on-target oracle response for each seed; solid lines indicate across-seed means and dashed lines the random-sequence baseline. **d,** Sequence composition and design diversity, quantified by GC content, within-run sequence divergence, and the number of distinct motifs used across designs. The gray band indicates 45–55% GC content and the dashed line the full 637-motif vocabulary. **e,** Effect of contrastive optimization on specificity (left) and on-target activity (right). Lines connect tissue-level means for single-tissue and contrastive designs; dashed red lines highlight tissues in which increased specificity was accompanied by reduced on-target activity. **f,** Distribution of predicted heart specificity for contrastive REGAL designs and random sequences; dashed lines indicate distribution means. Specificity was defined as target-tissue activation maximum max off-target activity. All values are *in silico* Enformer predictions.

**Extended Data Fig. 4.**
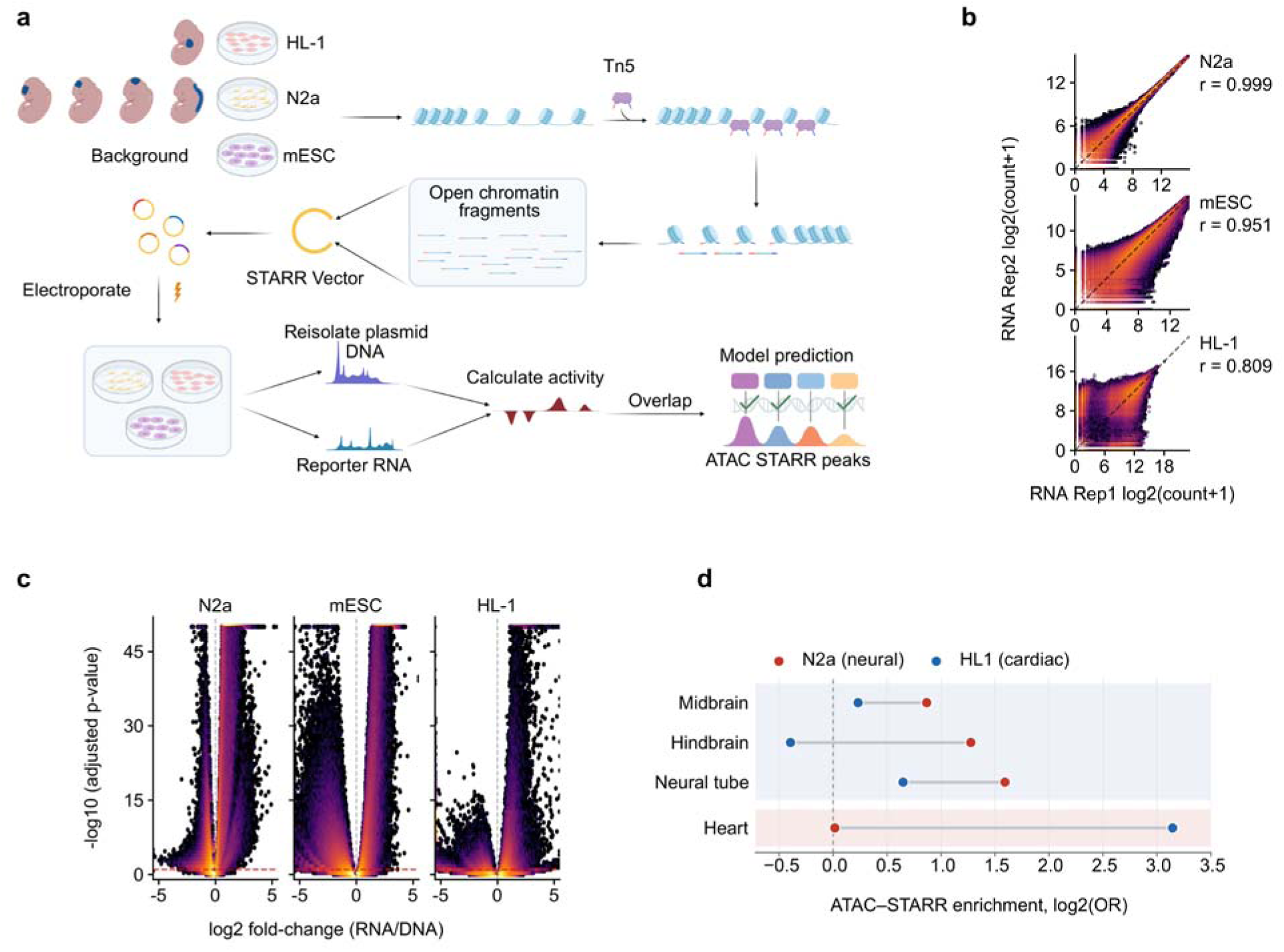
ATAC-STARR-seq reproducibility and enrichment of oracle-predicted regions. **a**, Schematic of the ATAC-STARR-seq assay in N2a (Neuro-2a), HL-1 (cardiomyocyte cell line), and mESCs (mouse embryonic stem cells). **b,** Reproducibility of reporter-RNA counts between two representative replicates for each cell line. The dashed line indicates the identity line, and point color represents local density. **c,** Differential reporter activity identified by DESeq2 from three paired DNA and RNA replicates per cell line. The dashed lines indicate log₂(RNA/DNA) = 0 and a Benjamini–Hochberg-adjusted P value of 0.1. Adjacent significant fragments were merged, yielding 25,791 active and 103,193 silent elements in N2a, 34,619 active and 56,751 silent elements in mESC, and 12,089 active and 147,442 silent elements in HL-1. **d,** Enrichment of the top 5,000 Enformer-predicted regions for overlap with cell-type-specific ATAC-STARR-seq activity. Neural-tissue predictions were preferentially enriched in N2a-active elements, whereas heart predictions were preferentially enriched in HL-1-active elements. Points show log₂ odds ratios relative to matched genomic controls. Enrichment was assessed using two-sided Fisher’s exact tests with Benjamini–Hochberg correction across tissue–cell-line comparisons.

**Extended Data Fig. 5.**
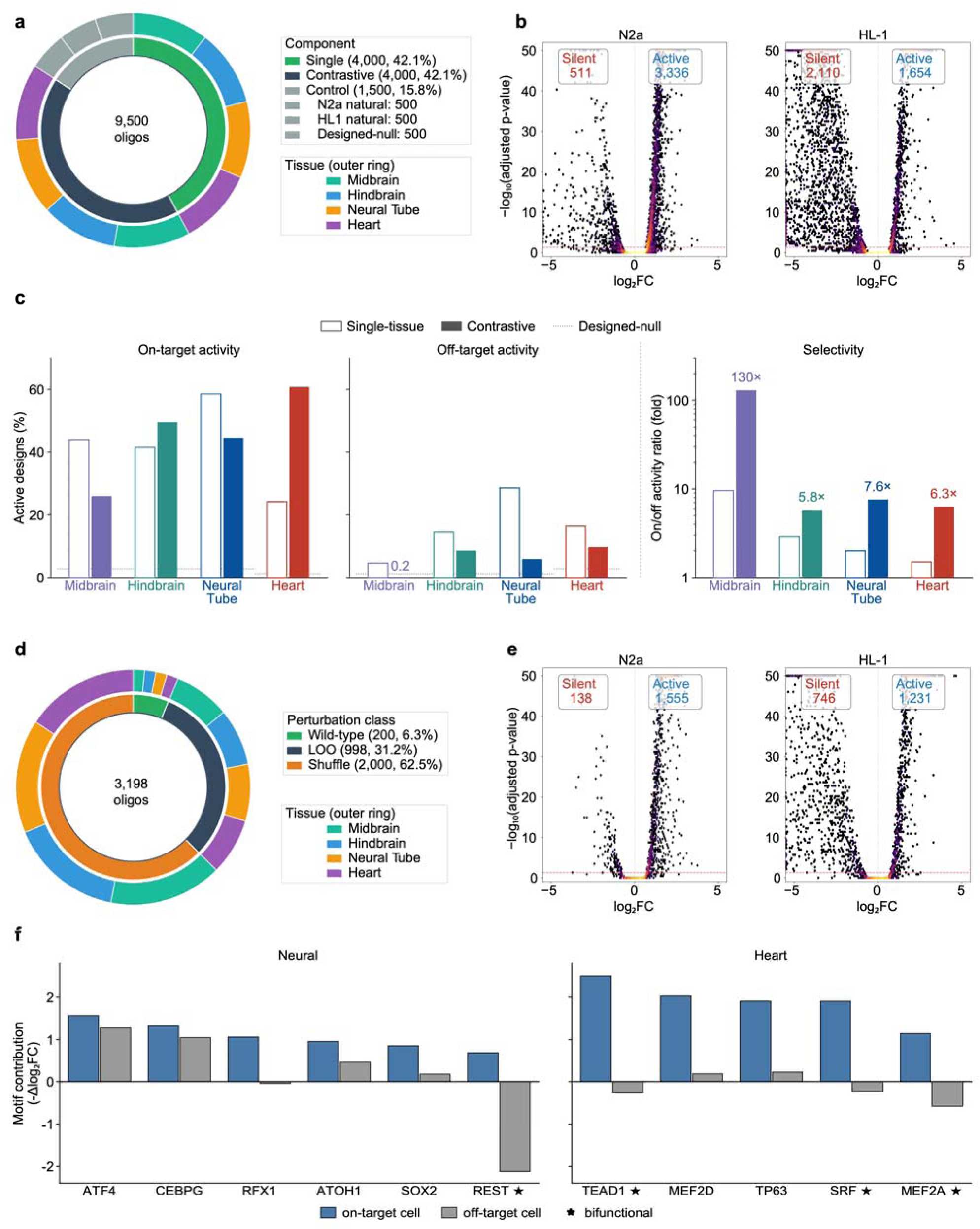
STARR-seq validation and perturbation analysis of REGAL designs. **a**, Composition of the 9,500-oligo synthetic-enhancer library, comprising 4,000 single-tissue designs, 4,000 contrastive designs, and 1,500 control sequences. Colors in the outer ring indicate the target tissue. **b,** DESeq2 volcano plots of enhancer activity in N2a and HL-1 cells. **c,** On- and off-target activity and selectivity of contrastive versus single-tissue designs. Left, designs active in the target cell line; middle, the same designs in the other line; right, the ratio of the two (log axis). Open bars, single-tissue; filled bars, contrastive; dotted lines, designed-null controls. **d,** Composition of the perturbation library: WT controls, LOO ablations and position shuffles. **e,** STARR-seq activity of the perturbation library in N2a and HL-1 cells, displayed as in b. In **b,e,** activity was determined from three paired DNA/RNA replicates using DESeq2, with adjusted P ≤ 0.05. **f,** Effects of individual motif deletion on neural and cardiac enhancer activity. Motif contribution was defined as the change in activity upon deletion relative to the WT control (−Δlog₂FC); positive values indicate activity-promoting effects, whereas negative values indicate repressive effects. Blue and gray bars show effects in the on- and off-target cell lines, respectively, and stars denote motifs with dual activating and repressive functions.

**Extended Data Fig. 6.**
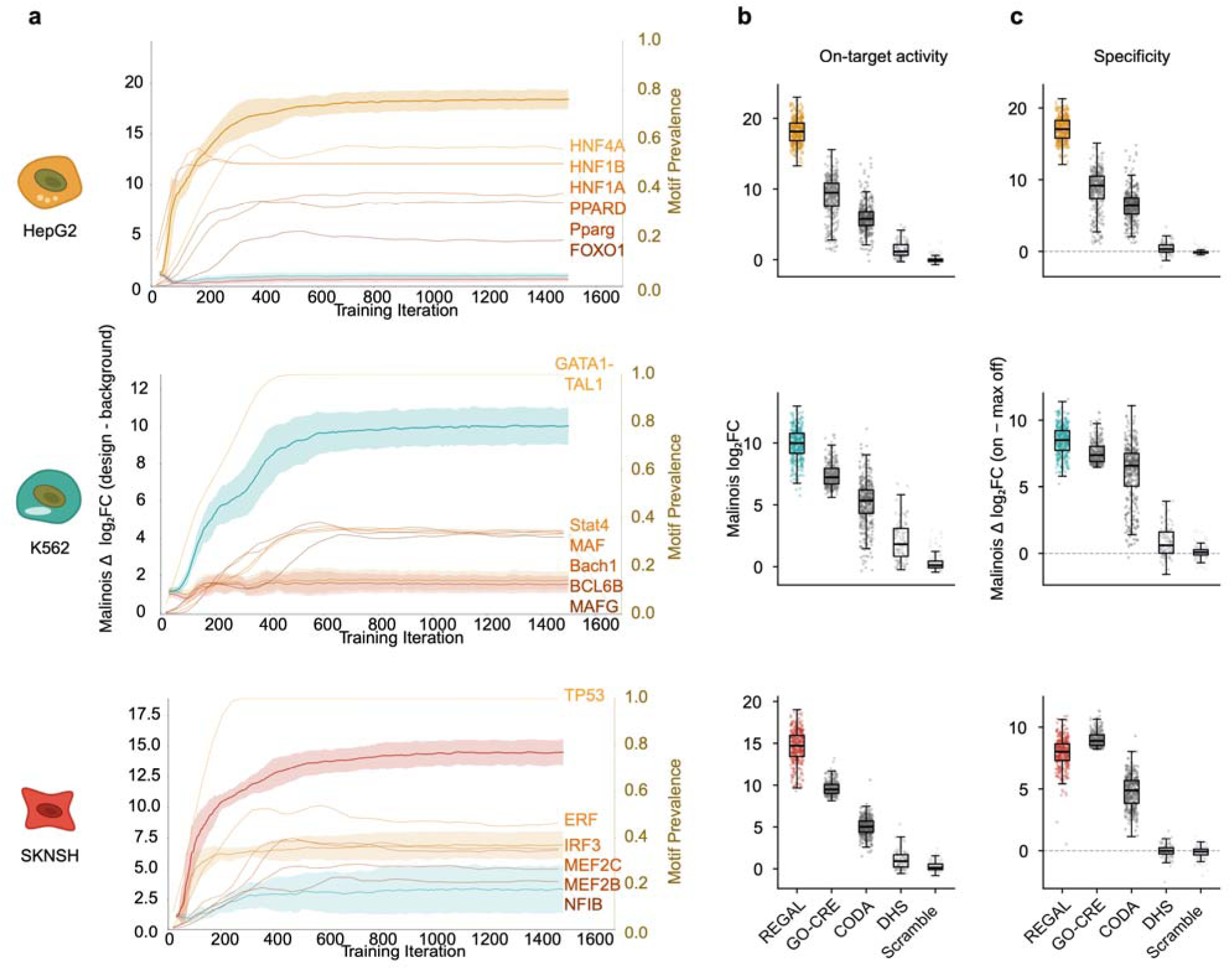
*In silico* performance of human cell-line-specific REGAL designs. **a,** Contrastive-training trajectories for HepG2, K562, and SK-N-SH. Thick lines show the target-cell Malinois oracle response and thin lines the off-target responses (mean ± standard deviation across eight independent training seeds). Orange curves show the prevalence of representative motifs enriched during training. **b,c,** Malinois-predicted log₂FC; b) and specificity (target minus maximum off-target log₂FC; c) for REGAL, Guided Optimization of Cis-Regulatory Elements (GO-CRE), CODA, DHS and dinucleotide-scrambled REGAL sequences. For GO-CRE, activity/specificity P values were below numerical precision in HepG2, 1 × 10⁻²³²/7 × 10⁻⁸⁵ in K562, and 7 × 10⁻²⁸⁷/1 × 10⁻¹²⁴ in SK-N-SH. Corresponding CODA activity/specificity P values were 6 × 10⁻¹⁶⁵/1 × 10⁻¹⁶⁴, 7 × 10⁻¹⁵⁶/9 × 10⁻⁸⁸, and 7 × 10⁻¹⁶⁵/5 × 10⁻¹⁴³ in HepG2, K562, and SK-N-SH, respectively.

**Extended Data Fig. 7.**
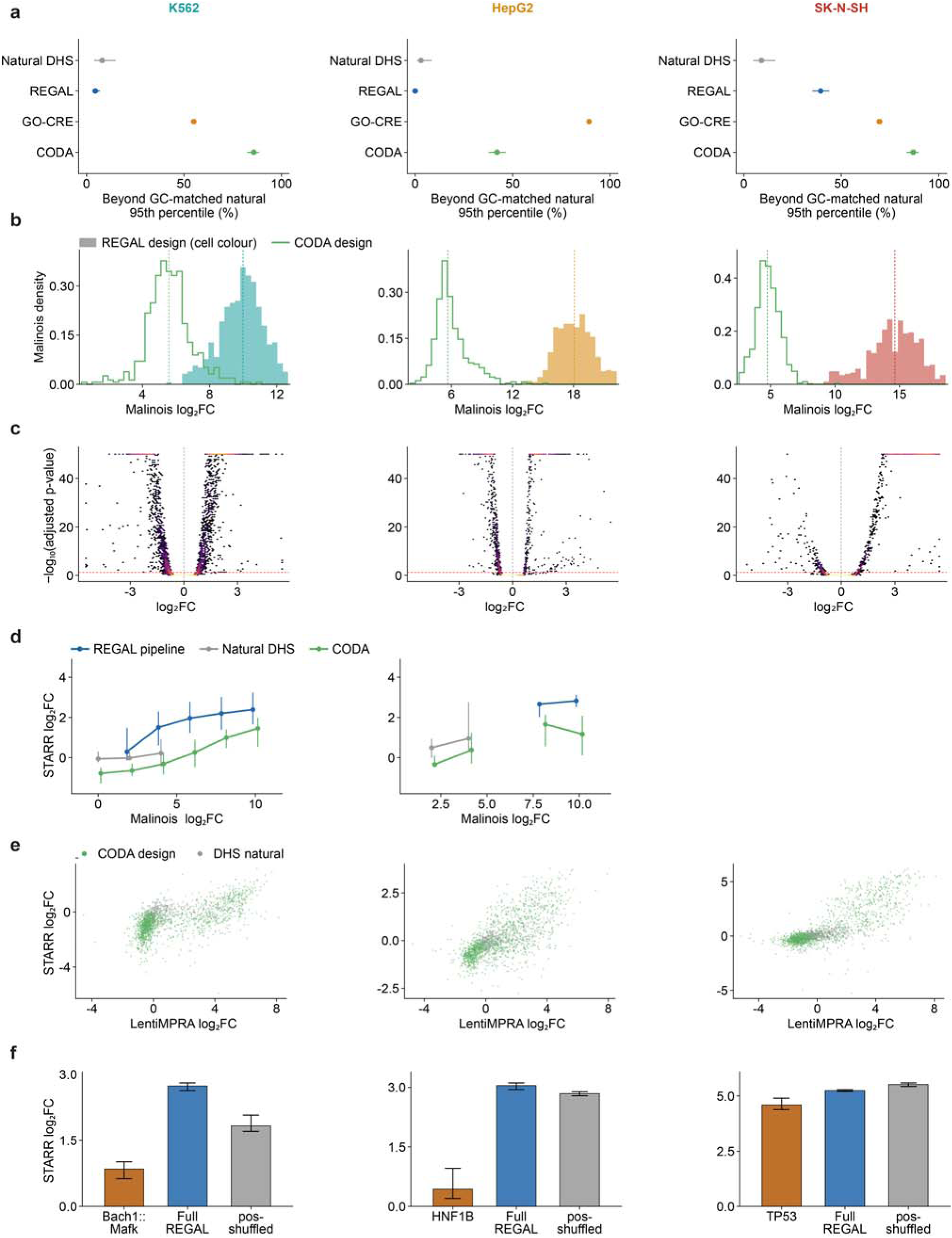
Sequence composition and experimental validation of human enhancer STARR-seq libraries. **a**, Fraction of each collection with CpG observed/expected ratios above the 95th percentile of GC-matched natural sequences. Points show the estimated fractions and horizontal bars indicate 95% confidence intervals. **b,** Malinois-predicted on-target activity distribution of the REGAL candidate pool and CODA designs. Dashed lines indicate group medians. **c,** STARR-seq activity for the human enhancer STARR-Seq library. **d,** Measured STARR-seq activity at matched ranges of Malinois-predicted activity in K562 and HepG2. Sequences were grouped into prediction bins, and points and whiskers show the median and IQR of measured activity within each bin. **e,** Concordance between STARR-seq activity and published lentiMPRA activity for CODA designs and natural DHS elements. Associations were assessed by Spearman correlation. **f,** Measured activity of the strongest individual motif associated with each cell type, the intact REGAL enhancer and its position-shuffled counterpart. Bars show medians and error bars indicate 95% bootstrap confidence intervals.

**Extended Data Fig. 8.**
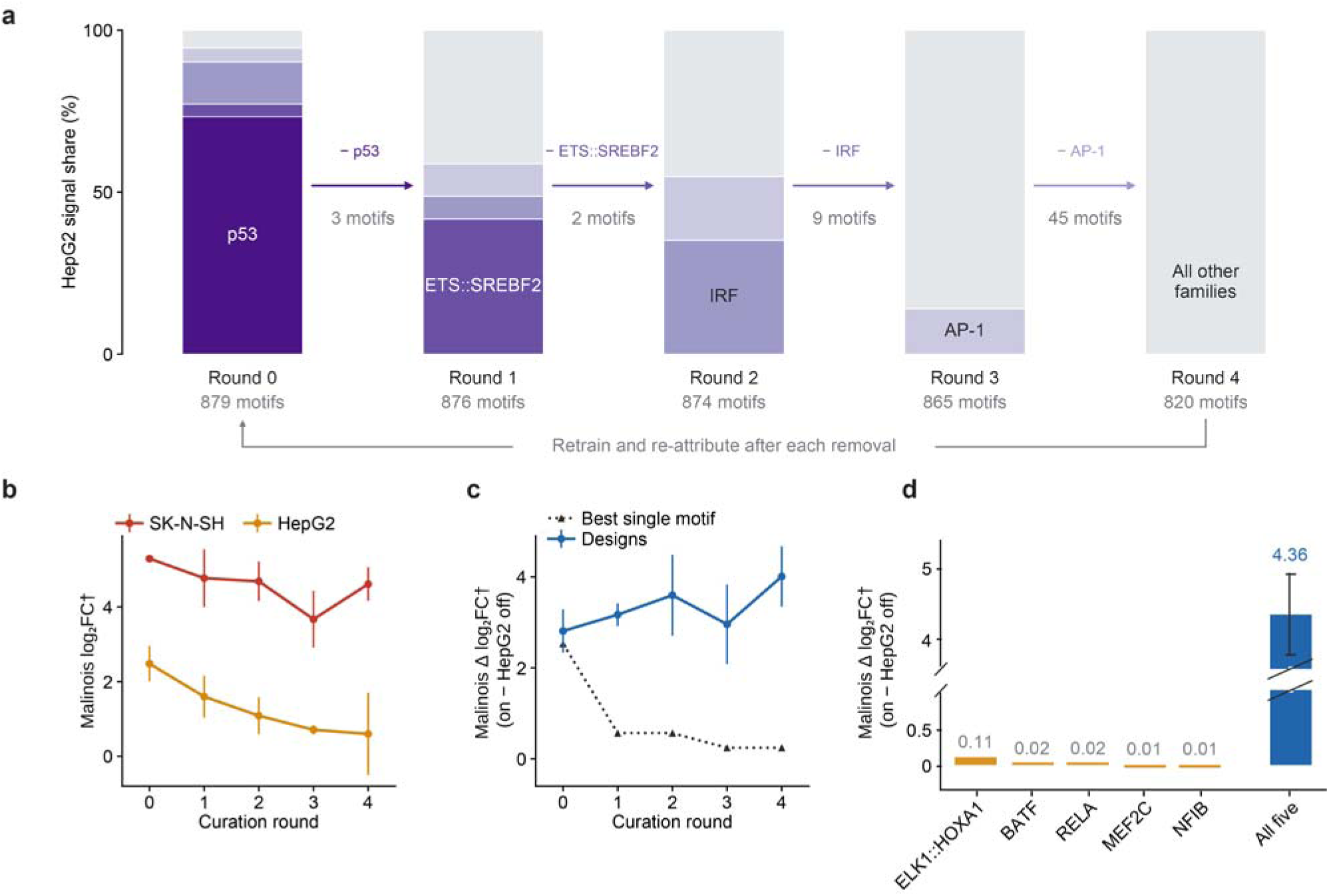
Iterative motif-vocabulary curation of the SK-N-SH design policy. **a**, Iterative attribution and curation of the motif vocabulary based on predicted off-target HepG2 activity. Stacked bars show the fraction of HepG2 signal attributed to TF families at each round. Arrows indicate the family removed before retraining, with the number of motifs removed shown below. **b,** Predicted SK-N-SH and HepG2 activity across successive curation rounds. Points show the mean across three independently trained seeds and error bars indicate ±1 standard deviation. **c,** Predicted specificity (SK-N-SH minus HepG2) of REGAL designs compared with the strongest individual motif remaining in each round’s vocabulary. **d,** Specificity of the five individual motifs comprising the most common motif set from the final curated policy compared with the complete five-motif design. The complete-design bar shows the mean ±1 standard deviation across 1,578 designs sharing this motif composition. Daggers denote Malinois-derived values transformed to the measured STARR-seq log₂FC scale as described in Methods 8.7.

**Extended Data Fig. 9.**
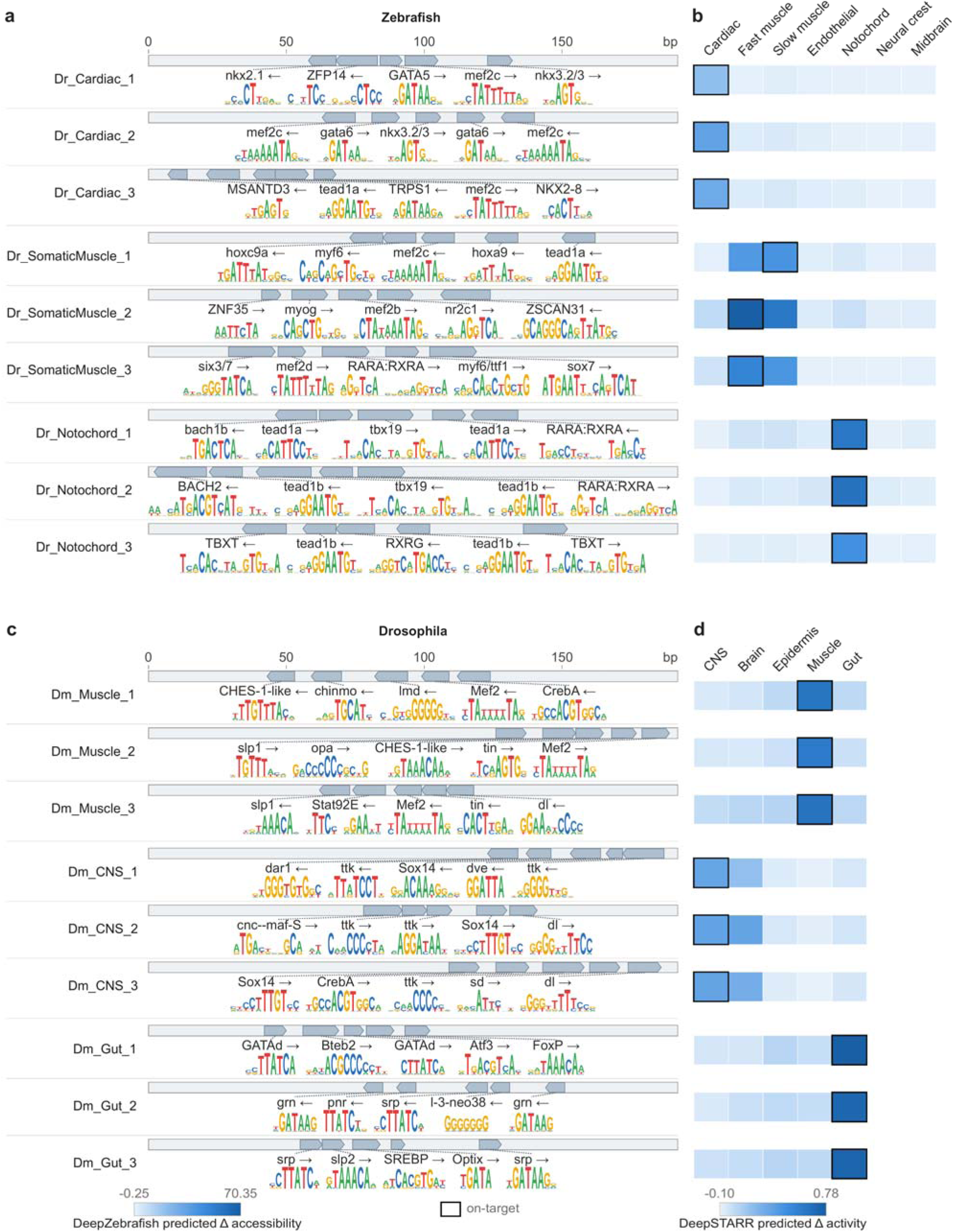
Motif grammars and cross-tissue predictions for zebrafish and *Drosophila* designs. **a**, Motif maps of nine zebrafish designs targeting cardiac muscle, somatic muscle, and notochord. Each design contains five motifs positioned within the 192-bp design window; arrowheads indicate motif orientation and sequence logos show motif identity. **b,** DeepZebrafish-predicted accessibility of the corresponding designs across seven cell-type outputs. Colors indicate the predicted change in accessibility relative to the matched background, and black outlines mark the intended target. **c,** Corresponding motif maps for nine *Drosophila* designs (three each targeting muscle, CNS and gut). **d,** DeepSTARR-predicted activity of the corresponding designs across five tissue outputs. Colors indicate the predicted change in activity relative to background, and black outlines mark the intended target.

**Extended Data Fig. 10.**
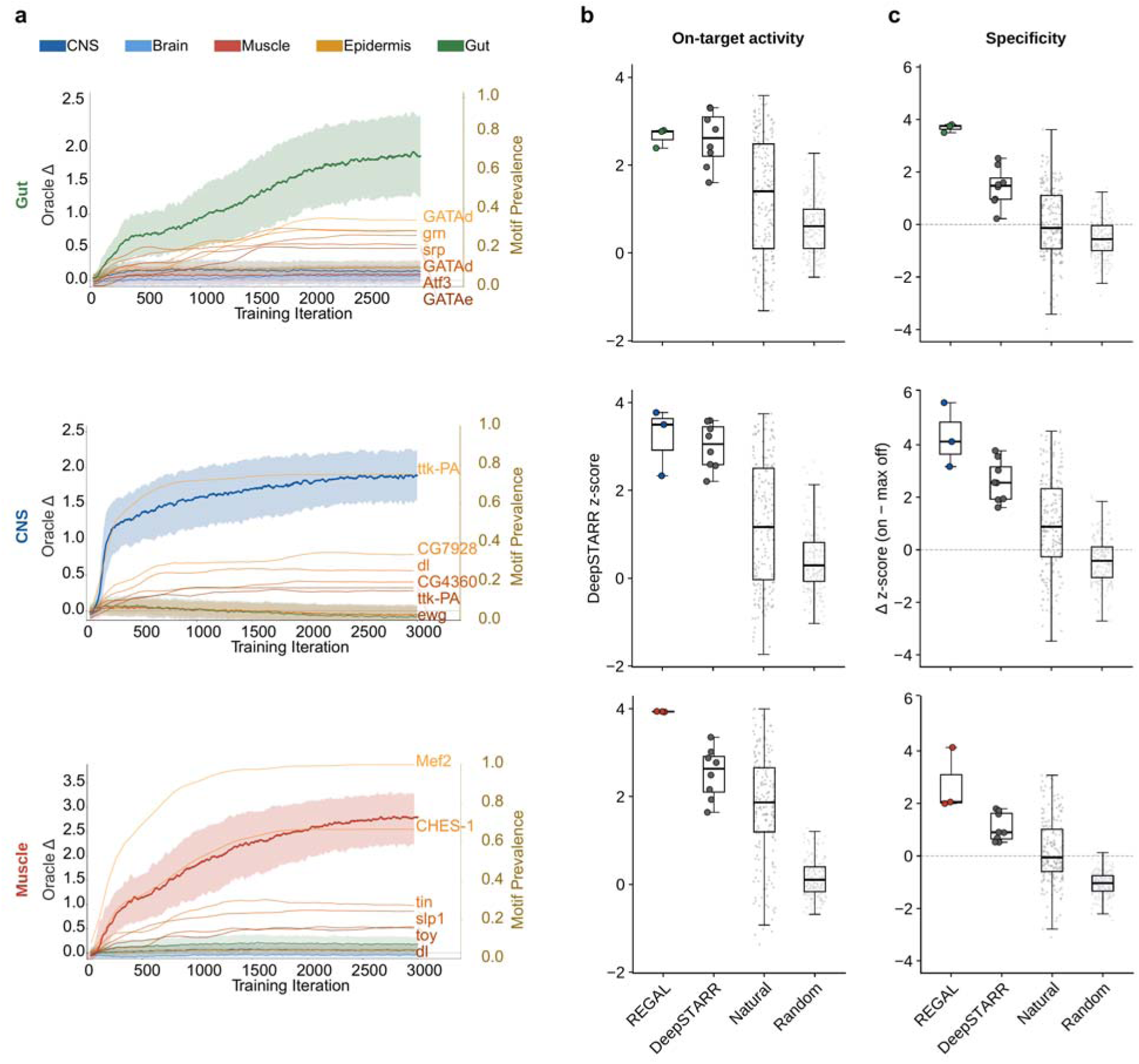
*Drosophila* training convergence and *in silico* benchmarking. **a,** REGAL policy training trajectories targeting gut, CNS and muscle. Thick lines show the target-tissue oracle response and thin lines the off-target responses (mean ± standard deviation across 24 independent training seeds). Orange curves show the prevalence of representative motifs enriched during training. **b,c,** DeepSTARR-predicted on-target activity (b) and specificity (c) of REGAL designs compared with published DeepSTARR designs, natural enhancers and random sequences. Activity is shown as the background-normalized target-tissue score, and specificity as the target score minus the strongest off-target score. The comparison included 3 REGAL, 8 published DeepSTARR, 200 natural and 200 random sequences per tissue. Boxes show the median and IQR, whiskers extend to 1.5 × IQR, and points represent individual sequences. The dashed line in c indicates zero specificity.

