## Supplementary_Information for "Regulatory grammar design of tissue-specific enhancers across species"

### Supplementary Methods

Section numbering follow the main text.

**2. Sequence-to-function oracle**

##### 2.1 Architecture benchmark Model configurations. Enformer was initialized from EleutherAI/enformer-official-rough with target_length=48, dropout 0.2, no organism-specific output heads, and use_tf_gamma=False. Its 48×3,072 embedding was adaptively average-pooled and passed through a 3,072→3,072→1,024→512→2 multilayer perceptron, with batch normalization, ReLU, and dropout 0.2 after each hidden layer.

DeepSTARR was trained from scratch. Its trunk comprised four same-padded Conv1d blocks with 256, 60, 60, and 120 filters and kernel widths 7, 3, 5, and 3, respectively. Each convolution was followed by batch normalization, ReLU, max pooling (width and stride 2), and dropout 0.4. The flattened 37,440-dimensional representation was passed through hidden layers of width 3,072, 1,024, and 512, each with batch normalization, ReLU, and dropout 0.2, followed by a two-logit output layer.

DNABERT-2 was initialized from zhihan1996/DNABERT-2-117M. Its pooled [CLS] embedding (768-dimensional) was passed through a 768→3,072→1,024→512→2 classifier, with batch normalization, ReLU, and dropout 0.2 after each hidden layer. The HyenaDNA models were initialized from the LongSafari hyenadna-small-32k-seqlen, hyenadna-medium-450k-seqlen, and hyenadna-large-1m-seqlen checkpoints. All three had model width 256; the 32K model had four layers, whereas the 450K and 1M models had eight. Their pooled embeddings were passed through a 256→3,072→1,024→512→2 classifier, with batch normalization, ReLU, and dropout 0.2 after each hidden layer. Thus, all four architectures shared the same 3,072→1,024→512 hidden-width classifier after architecture-specific pooling or flattening.

**Training and checkpoint selection.** All benchmark runs used two-class cross-entropy loss and were trained for 20 epochs. Class imbalance was handled by inverse-frequency sample weights supplied to a WeightedRandomSampler with replacement and num_samples equal to the training-set size; the loss itself was unweighted. Enformer, DeepSTARR, and HyenaDNA used batch size 32, and DNABERT-2 used batch size 16.

Enformer was fine-tuned end-to-end with AdamW, using learning rates of 1×10⁻⁶ for the pretrained backbone and 1×10⁻⁵ for the classifier and weight decay 1×10⁻⁴. A cosine-annealing-with-warm-restarts scheduler (T_0=5, T_mult=2, eta_min=1×10⁻⁶) was stepped after each optimization batch. The checkpoint with the highest validation area under the precision–recall curve (AUPRC) was retained, and validation and test logits were averaged over forward and reverse-complement inputs before softmax.

DNABERT-2 was fine-tuned end-to-end with AdamW at 1×10⁻⁵ and no explicitly specified weight decay. HyenaDNA used AdamW at 1×10⁻⁵ with weight decay 0.01, and DeepSTARR used Adam at 1×10⁻⁵ with default zero weight decay; no learning-rate scheduler was applied to these models. Their checkpoints were selected by validation area under the receiver operating characteristic curve (AUROC) and evaluated on the forward strand. HyenaDNA training used reverse-complement augmentation with probability 0.5; DNABERT-2 and DeepSTARR did not.

**Evaluation.** AUROC was the primary discrimination metric. The precision–recall metric reported as AUPRC was calculated with scikit-learn average_precision_score. The positive-class score was the softmax probability assigned to class 1. Threshold-independent metrics drove the architecture comparison. For the secondary threshold-dependent metrics in Supplementary Table 2, Enformer used a probability threshold of 0.5, and DeepSTARR, DNABERT-2 and HyenaDNA used the test-set receiver operating characteristic (ROC) threshold minimizing Euclidean distance to the upper-left corner. For each tissue and model, the mean and 95% confidence interval across five seeds were calculated using the sample standard deviation, standard error, and the two-sided *t* critical value with 4 degrees of freedom. Overall performance was the equal-weight mean of the seven tissue means.

##### 2.2 Enformer oracle fine-tuning

On the basis of the architecture benchmark, Enformer was retrained as the production sequence-to-function oracle at the 1,536-bp input length used for downstream design and scoring. This span produces 12 output bins of 128 bp. One independent binary classifier was trained for each of the seven tissues.

**Architecture and inputs.** Each oracle used an Enformer backbone with model dimension 1,536, depth 11, eight attention heads, target_length=12, dropout 0.2, no organism-specific output heads, and use_tf_gamma=False. The backbone was initialized from matching names and tensor shapes in EleutherAI/enformer-official-rough; all backbone parameters remained trainable during fine-tuning. The 12×3,072 embedding was adaptively average-pooled across bins and passed through a 3,072→1,024→512→2 classifier, with batch normalization, ReLU, and dropout 0.2 after each hidden layer.

Candidate oracles were trained with five random seeds for 20 epochs using AdamW, inverse-frequency weighted sampling, and cross-entropy loss. Optimization used a learning rate of 1 × 10⁻⁴, weight decay of 1 × 10⁻⁵, and cosine annealing. For each seed, checkpoints were selected by validation AUPRC or AUROC. Production models used the AUPRC-best checkpoint for six tissues and the AUROC-best checkpoint for hindbrain; deployed checkpoints and held-out performance are reported in **Supplementary Table 3**.

Tissue activity was defined as the active-class probability from the tissue-specific binary classifier. Oracles scored the full 1,536-bp sequence. Validation and post hoc analyses averaged forward- and reverse-complement logits before softmax, whereas REGAL optimization used a single forward-orientation score. Oracle parameters remained fixed during design. Threshold-dependent test metrics used the ROC threshold maximizing Youden’s index and are reported in **Supplementary Table 3**.

### Supplementary Tables

**Supplementary Table 1 | Composition of the VISTA enhancer dataset used for oracle training and evaluation.** Per-tissue counts of enhancer-positive and enhancer-negative sequences across the training, validation and held-out test sets.

| **tissue** | **split** | **class** | **count** |
| --- | --- | --- | --- |
| Forebrain | train | positive | 516 |
| Forebrain | train | negative | 2,647 |
| Forebrain | val | positive | 74 |
| Forebrain | val | negative | 321 |
| Forebrain | test | positive | 68 |
| Forebrain | test | negative | 329 |
| Midbrain | train | positive | 456 |
| Midbrain | train | negative | 2,707 |
| Midbrain | val | positive | 60 |
| Midbrain | val | negative | 335 |
| Midbrain | test | positive | 58 |
| Midbrain | test | negative | 339 |
| Hindbrain | train | positive | 403 |
| Hindbrain | train | negative | 2,760 |
| Hindbrain | val | positive | 52 |
| Hindbrain | val | negative | 343 |
| Hindbrain | test | positive | 52 |
| Hindbrain | test | negative | 345 |
| Neural_Tube | train | positive | 444 |
| Neural_Tube | train | negative | 2,719 |
| Neural_Tube | val | positive | 55 |
| Neural_Tube | val | negative | 340 |
| Neural_Tube | test | positive | 44 |
| Neural_Tube | test | negative | 353 |
| Heart | train | positive | 278 |
| Heart | train | negative | 2,885 |
| Heart | val | positive | 37 |
| Heart | val | negative | 358 |
| Heart | test | positive | 38 |
| Heart | test | negative | 359 |
| Face | train | positive | 323 |
| Face | train | negative | 2,839 |
| Face | val | positive | 45 |
| Face | val | negative | 350 |
| Face | test | positive | 34 |
| Face | test | negative | 364 |
| Limb | train | positive | 402 |
| Limb | train | negative | 2,761 |
| Limb | val | positive | 52 |
| Limb | val | negative | 342 |
| Limb | test | positive | 40 |
| Limb | test | negative | 358 |

**Supplementary Table 2 | Held-out test-set performance of six sequence-to-function models across seven mouse tissues.** Classification performance of Enformer, DNABERT-2, DeepSTARR, and three HyenaDNA models (32K, 450K, and 1M pretraining context) on the seven VISTA tissue tasks.

| **Model** | **Tissue** | **Replicate** | **AUROC** | **AUPRC** | **Accuracy** | **F1** | **Precision** | **Recall** | **Specificity** |
| --- | --- | --- | --- | --- | --- | --- | --- | --- | --- |
| Enformer | Forebrain | 1 | 82.00 | 54.59 | 77.58 | 48.55 | 40.00 | 61.76 | 80.85 |
| Enformer | Forebrain | 2 | 83.31 | 54.38 | 77.58 | 51.89 | 41.03 | 70.59 | 79.03 |
| Enformer | Forebrain | 3 | 84.67 | 57.17 | 78.59 | 52.51 | 42.34 | 69.12 | 80.55 |
| Enformer | Forebrain | 4 | 83.32 | 55.26 | 83.12 | 52.48 | 50.68 | 54.41 | 89.06 |
| Enformer | Forebrain | 5 | 83.90 | 59.31 | 76.32 | 50.00 | 39.17 | 69.12 | 77.81 |
| Enformer | Forebrain | Mean (95% CI) | 83.44 (82.22–84.65) | 56.14 (53.55–58.73) | 78.64 (75.37–81.91) | 51.09 (48.92–53.26) | 42.64 (36.87–48.42) | 65.00 (56.49–73.51) | 81.46 (75.97–86.95) |
| Enformer | Midbrain | 1 | 76.43 | 46.24 | 78.59 | 45.86 | 36.36 | 62.07 | 81.42 |
| Enformer | Midbrain | 2 | 79.22 | 48.90 | 79.60 | 48.41 | 38.38 | 65.52 | 82.01 |
| Enformer | Midbrain | 3 | 78.55 | 44.91 | 76.57 | 44.31 | 33.94 | 63.79 | 78.76 |
| Enformer | Midbrain | 4 | 75.22 | 45.44 | 77.83 | 44.30 | 35.00 | 60.34 | 80.83 |
| Enformer | Midbrain | 5 | 76.73 | 43.19 | 78.84 | 42.47 | 35.23 | 53.45 | 83.19 |
| Enformer | Midbrain | Mean (95% CI) | 77.23 (75.20–79.26) | 45.74 (43.14–48.33) | 78.29 (76.86–79.71) | 45.07 (42.31–47.83) | 35.78 (33.69–37.88) | 61.03 (55.25–66.82) | 81.24 (79.21–83.27) |
| Enformer | Hindbrain | 1 | 74.91 | 47.94 | 79.85 | 42.03 | 33.72 | 55.77 | 83.48 |
| Enformer | Hindbrain | 2 | 80.54 | 51.03 | 79.60 | 46.36 | 35.35 | 67.31 | 81.45 |
| Enformer | Hindbrain | 3 | 81.21 | 47.81 | 81.36 | 47.14 | 37.50 | 63.46 | 84.06 |
| Enformer | Hindbrain | 4 | 77.13 | 43.54 | 83.12 | 39.64 | 37.29 | 42.31 | 89.28 |
| Enformer | Hindbrain | 5 | 74.47 | 45.94 | 81.61 | 37.61 | 33.85 | 42.31 | 87.54 |
| Enformer | Hindbrain | Mean (95% CI) | 77.65 (73.78–81.53) | 47.25 (43.82–50.68) | 81.11 (79.33–82.89) | 42.56 (37.41–47.70) | 35.54 (33.29–37.79) | 54.23 (39.77–68.70) | 85.16 (81.21–89.11) |
| Enformer | Neural_Tube | 1 | 67.05 | 22.89 | 71.28 | 28.75 | 19.83 | 52.27 | 73.65 |
| Enformer | Neural_Tube | 2 | 71.50 | 25.73 | 72.29 | 32.93 | 22.50 | 61.36 | 73.65 |
| Enformer | Neural_Tube | 3 | 69.58 | 24.68 | 71.03 | 30.30 | 20.66 | 56.82 | 72.80 |
| Enformer | Neural_Tube | 4 | 71.61 | 27.04 | 80.86 | 34.48 | 27.78 | 45.45 | 85.27 |
| Enformer | Neural_Tube | 5 | 68.45 | 22.59 | 78.09 | 23.01 | 18.84 | 29.55 | 84.14 |
| Enformer | Neural_Tube | Mean (95% CI) | 69.64 (67.19–72.08) | 24.59 (22.24–26.93) | 74.71 (69.15–80.27) | 29.89 (24.37–35.42) | 21.92 (17.53–26.32) | 49.09 (33.69–64.50) | 77.90 (70.17–85.64) |
| Enformer | Heart | 1 | 85.05 | 51.47 | 80.60 | 39.37 | 28.09 | 65.79 | 82.17 |
| Enformer | Heart | 2 | 82.33 | 45.41 | 74.56 | 37.27 | 24.39 | 78.95 | 74.09 |
| Enformer | Heart | 3 | 83.02 | 48.45 | 76.32 | 37.33 | 25.00 | 73.68 | 76.60 |
| Enformer | Heart | 4 | 83.71 | 49.82 | 80.10 | 41.48 | 28.87 | 73.68 | 80.78 |
| Enformer | Heart | 5 | 88.54 | 59.79 | 83.88 | 49.21 | 35.23 | 81.58 | 84.12 |
| Enformer | Heart | Mean (95% CI) | 84.53 (81.48–87.58) | 50.99 (44.29–57.69) | 79.09 (74.51–83.67) | 40.93 (34.80–47.06) | 28.31 (22.96–33.67) | 74.74 (67.21–82.26) | 79.55 (74.44–84.66) |
| Enformer | Face | 1 | 69.38 | 30.30 | 86.18 | 30.38 | 26.67 | 35.29 | 90.93 |
| Enformer | Face | 2 | 63.41 | 19.65 | 73.12 | 24.11 | 15.89 | 50.00 | 75.27 |
| Enformer | Face | 3 | 69.21 | 26.76 | 46.98 | 19.77 | 11.35 | 76.47 | 44.23 |
| Enformer | Face | 4 | 62.37 | 21.05 | 78.39 | 24.56 | 17.50 | 41.18 | 81.87 |
| Enformer | Face | 5 | 67.40 | 23.71 | 74.87 | 25.37 | 17.00 | 50.00 | 77.20 |
| Enformer | Face | Mean (95% CI) | 66.36 (62.29–70.43) | 24.29 (18.93–29.65) | 71.91 (53.52–90.30) | 24.84 (20.14–29.54) | 17.68 (10.75–24.61) | 50.59 (31.02–70.15) | 73.90 (51.98–95.82) |
| Enformer | Limb | 1 | 78.47 | 30.81 | 83.67 | 36.89 | 30.16 | 47.50 | 87.71 |
| Enformer | Limb | 2 | 78.51 | 38.24 | 76.63 | 37.58 | 25.69 | 70.00 | 77.37 |
| Enformer | Limb | 3 | 81.93 | 52.31 | 78.64 | 42.18 | 28.97 | 77.50 | 78.77 |
| Enformer | Limb | 4 | 80.89 | 49.43 | 82.66 | 42.98 | 32.10 | 65.00 | 84.64 |
| Enformer | Limb | 5 | 82.56 | 32.09 | 84.67 | 41.90 | 33.85 | 55.00 | 87.99 |
| Enformer | Limb | Mean (95% CI) | 80.47 (78.11–82.84) | 40.58 (28.33–52.82) | 81.26 (76.97–85.54) | 40.31 (36.78–43.83) | 30.15 (26.29–34.02) | 63.00 (48.21–77.79) | 83.30 (77.12–89.47) |
| DNABERT-2 | Forebrain | 1 | 73.23 | 40.72 | 70.78 | 43.69 | 32.61 | 66.18 | 71.73 |
| DNABERT-2 | Forebrain | 2 | 73.55 | 36.18 | 70.78 | 43.69 | 32.61 | 66.18 | 71.73 |
| DNABERT-2 | Forebrain | 3 | 69.76 | 37.87 | 74.56 | 44.20 | 35.40 | 58.82 | 77.81 |
| DNABERT-2 | Forebrain | 4 | 72.15 | 37.74 | 62.47 | 39.68 | 27.37 | 72.06 | 60.49 |
| DNABERT-2 | Forebrain | 5 | 60.35 | 25.91 | 54.16 | 33.58 | 22.33 | 67.65 | 51.37 |
| DNABERT-2 | Forebrain | Mean (95% CI) | 69.81 (62.98–76.63) | 35.68 (28.60–42.77) | 66.55 (56.34–76.75) | 40.97 (35.36–46.57) | 30.06 (23.60–36.53) | 66.18 (60.26–72.09) | 66.63 (53.49–79.76) |
| DNABERT-2 | Midbrain | 1 | 65.55 | 23.80 | 63.48 | 34.39 | 23.31 | 65.52 | 63.13 |
| DNABERT-2 | Midbrain | 2 | 65.81 | 26.54 | 66.75 | 33.33 | 23.57 | 56.90 | 68.44 |
| DNABERT-2 | Midbrain | 3 | 63.57 | 23.07 | 56.93 | 29.63 | 19.46 | 62.07 | 56.05 |
| DNABERT-2 | Midbrain | 4 | 67.35 | 27.64 | 67.25 | 35.64 | 25.00 | 62.07 | 68.14 |
| DNABERT-2 | Midbrain | 5 | 62.44 | 26.83 | 65.49 | 30.46 | 21.58 | 51.72 | 67.85 |
| DNABERT-2 | Midbrain | Mean (95% CI) | 64.94 (62.53–67.36) | 25.58 (23.08–28.07) | 63.98 (58.76–69.20) | 32.69 (29.50–35.88) | 22.59 (19.94–25.23) | 59.66 (52.95–66.36) | 64.72 (58.12–71.32) |
| DNABERT-2 | Hindbrain | 1 | 71.31 | 28.28 | 66.25 | 33.66 | 22.67 | 65.38 | 66.38 |
| DNABERT-2 | Hindbrain | 2 | 74.19 | 29.31 | 69.02 | 39.41 | 26.49 | 76.92 | 67.83 |
| DNABERT-2 | Hindbrain | 3 | 67.05 | 24.25 | 64.23 | 33.02 | 21.88 | 67.31 | 63.77 |
| DNABERT-2 | Hindbrain | 4 | 73.31 | 28.94 | 71.28 | 37.36 | 26.15 | 65.38 | 72.17 |
| DNABERT-2 | Hindbrain | 5 | 74.83 | 35.19 | 76.07 | 41.72 | 30.63 | 65.38 | 77.68 |
| DNABERT-2 | Hindbrain | Mean (95% CI) | 72.14 (68.24–76.03) | 29.19 (24.33–34.06) | 69.37 (63.65–75.09) | 37.03 (32.42–41.65) | 25.56 (21.22–29.90) | 68.08 (61.85–74.30) | 69.57 (62.78–76.35) |
| DNABERT-2 | Neural_Tube | 1 | 62.29 | 17.21 | 59.95 | 26.05 | 16.37 | 63.64 | 59.49 |
| DNABERT-2 | Neural_Tube | 2 | 50.84 | 11.92 | 49.37 | 19.92 | 12.08 | 56.82 | 48.44 |
| DNABERT-2 | Neural_Tube | 3 | 63.42 | 17.01 | 55.16 | 25.21 | 15.46 | 68.18 | 53.54 |
| DNABERT-2 | Neural_Tube | 4 | 66.19 | 16.30 | 65.24 | 31.00 | 19.87 | 70.45 | 64.59 |
| DNABERT-2 | Neural_Tube | 5 | 64.25 | 18.05 | 58.69 | 25.45 | 15.91 | 63.64 | 58.07 |
| DNABERT-2 | Neural_Tube | Mean (95% CI) | 61.40 (53.86–68.94) | 16.10 (13.10–19.10) | 57.68 (50.37–64.99) | 25.53 (20.65–30.41) | 15.94 (12.49–19.38) | 64.55 (58.05–71.04) | 56.83 (49.22–64.43) |
| DNABERT-2 | Heart | 1 | 63.30 | 17.86 | 60.96 | 24.39 | 14.97 | 65.79 | 60.45 |
| DNABERT-2 | Heart | 2 | 64.49 | 14.32 | 57.18 | 26.09 | 15.62 | 78.95 | 54.87 |
| DNABERT-2 | Heart | 3 | 70.17 | 21.02 | 64.74 | 25.53 | 16.00 | 63.16 | 64.90 |
| DNABERT-2 | Heart | 4 | 70.63 | 20.98 | 62.72 | 28.85 | 17.65 | 78.95 | 61.00 |
| DNABERT-2 | Heart | 5 | 57.17 | 10.85 | 53.15 | 21.19 | 12.63 | 65.79 | 51.81 |
| DNABERT-2 | Heart | Mean (95% CI) | 65.15 (58.27–72.03) | 17.01 (11.54–22.48) | 59.75 (54.01–65.48) | 25.21 (21.75–28.66) | 15.37 (13.11–17.64) | 70.53 (60.89–80.16) | 58.61 (52.13–65.09) |
| DNABERT-2 | Face | 1 | 50.68 | 10.12 | 58.79 | 16.33 | 9.88 | 47.06 | 59.89 |
| DNABERT-2 | Face | 2 | 60.99 | 15.41 | 61.06 | 19.69 | 11.95 | 55.88 | 61.54 |
| DNABERT-2 | Face | 3 | 68.79 | 21.44 | 60.30 | 24.76 | 14.77 | 76.47 | 58.79 |
| DNABERT-2 | Face | 4 | 64.02 | 14.37 | 54.52 | 21.65 | 12.69 | 73.53 | 52.75 |
| DNABERT-2 | Face | 5 | 68.44 | 21.18 | 69.35 | 27.38 | 17.16 | 67.65 | 69.51 |
| DNABERT-2 | Face | Mean (95% CI) | 62.58 (53.39–71.77) | 16.50 (10.53–22.48) | 60.80 (54.09–67.51) | 21.96 (16.61–27.31) | 13.29 (9.83–16.75) | 64.12 (48.75–79.48) | 60.49 (53.00–67.99) |
| DNABERT-2 | Limb | 1 | 67.54 | 21.07 | 63.57 | 27.14 | 16.98 | 67.50 | 63.13 |
| DNABERT-2 | Limb | 2 | 48.25 | 10.99 | 44.22 | 17.78 | 10.43 | 60.00 | 42.46 |
| DNABERT-2 | Limb | 3 | 59.01 | 17.99 | 61.31 | 23.76 | 14.81 | 60.00 | 61.45 |
| DNABERT-2 | Limb | 4 | 57.77 | 12.03 | 60.80 | 24.27 | 15.06 | 62.50 | 60.61 |
| DNABERT-2 | Limb | 5 | 61.79 | 14.53 | 57.54 | 24.89 | 15.14 | 70.00 | 56.15 |
| DNABERT-2 | Limb | Mean (95% CI) | 58.87 (50.15–67.60) | 15.32 (10.12–20.53) | 57.49 (47.90–67.08) | 23.57 (19.24–27.89) | 14.49 (11.48–17.49) | 64.00 (58.36–69.64) | 56.76 (46.33–67.19) |
| HyenaDNA-32K | Forebrain | 1 | 67.44 | 30.13 | 61.71 | 37.70 | 26.14 | 67.65 | 60.49 |
| HyenaDNA-32K | Forebrain | 2 | 64.08 | 25.81 | 58.19 | 34.65 | 23.66 | 64.71 | 56.84 |
| HyenaDNA-32K | Forebrain | 3 | 70.58 | 36.79 | 64.48 | 40.51 | 28.40 | 70.59 | 63.22 |
| HyenaDNA-32K | Forebrain | 4 | 70.21 | 32.47 | 65.99 | 43.51 | 30.41 | 76.47 | 63.83 |
| HyenaDNA-32K | Forebrain | 5 | 69.48 | 35.37 | 61.71 | 37.70 | 26.14 | 67.65 | 60.49 |
| HyenaDNA-32K | Forebrain | Mean (95% CI) | 68.36 (65.03–71.69) | 32.12 (26.70–37.54) | 62.42 (58.70–66.14) | 38.82 (34.66–42.97) | 26.95 (23.77–30.13) | 69.41 (63.87–74.95) | 60.97 (57.53–64.42) |
| HyenaDNA-32K | Midbrain | 1 | 62.50 | 20.52 | 60.45 | 31.44 | 21.05 | 62.07 | 60.18 |
| HyenaDNA-32K | Midbrain | 2 | 59.90 | 19.76 | 56.17 | 29.27 | 19.15 | 62.07 | 55.16 |
| HyenaDNA-32K | Midbrain | 3 | 64.08 | 22.58 | 61.96 | 32.29 | 21.82 | 62.07 | 61.95 |
| HyenaDNA-32K | Midbrain | 4 | 61.31 | 22.53 | 60.96 | 31.11 | 20.96 | 60.34 | 61.06 |
| HyenaDNA-32K | Midbrain | 5 | 56.86 | 17.86 | 60.96 | 29.22 | 19.88 | 55.17 | 61.95 |
| HyenaDNA-32K | Midbrain | Mean (95% CI) | 60.93 (57.52–64.34) | 20.65 (18.18–23.12) | 60.10 (57.29–62.91) | 30.67 (28.97–32.36) | 20.57 (19.26–21.88) | 60.34 (56.64–64.05) | 60.06 (56.54–63.58) |
| HyenaDNA-32K | Hindbrain | 1 | 56.54 | 18.27 | 65.74 | 26.09 | 18.18 | 46.15 | 68.70 |
| HyenaDNA-32K | Hindbrain | 2 | 58.51 | 18.05 | 54.41 | 27.89 | 17.59 | 67.31 | 52.46 |
| HyenaDNA-32K | Hindbrain | 3 | 56.42 | 18.05 | 59.19 | 23.58 | 15.62 | 48.08 | 60.87 |
| HyenaDNA-32K | Hindbrain | 4 | 59.59 | 19.27 | 60.20 | 28.18 | 18.45 | 59.62 | 60.29 |
| HyenaDNA-32K | Hindbrain | 5 | 56.67 | 17.20 | 69.77 | 26.83 | 19.64 | 42.31 | 73.91 |
| HyenaDNA-32K | Hindbrain | Mean (95% CI) | 57.55 (55.78–59.32) | 18.17 (17.25–19.08) | 61.86 (54.44–69.29) | 26.51 (24.23–28.80) | 17.90 (16.07–19.73) | 52.69 (39.77–65.62) | 63.25 (52.97–73.53) |
| HyenaDNA-32K | Neural_Tube | 1 | 63.89 | 16.77 | 57.93 | 27.07 | 16.76 | 70.45 | 56.37 |
| HyenaDNA-32K | Neural_Tube | 2 | 65.59 | 16.62 | 63.22 | 30.48 | 19.28 | 72.73 | 62.04 |
| HyenaDNA-32K | Neural_Tube | 3 | 64.71 | 15.60 | 62.47 | 30.05 | 18.93 | 72.73 | 61.19 |
| HyenaDNA-32K | Neural_Tube | 4 | 62.61 | 15.93 | 59.70 | 26.61 | 16.67 | 65.91 | 58.92 |
| HyenaDNA-32K | Neural_Tube | 5 | 64.60 | 17.21 | 60.45 | 27.65 | 17.34 | 68.18 | 59.49 |
| HyenaDNA-32K | Neural_Tube | Mean (95% CI) | 64.28 (62.90–65.66) | 16.43 (15.62–17.24) | 60.76 (58.11–63.40) | 28.37 (26.17–30.57) | 17.80 (16.27–19.32) | 70.00 (66.32–73.68) | 59.60 (56.87–62.33) |
| HyenaDNA-32K | Heart | 1 | 75.66 | 22.64 | 68.26 | 31.52 | 19.86 | 76.32 | 67.41 |
| HyenaDNA-32K | Heart | 2 | 73.58 | 24.12 | 69.77 | 29.41 | 18.94 | 65.79 | 70.19 |
| HyenaDNA-32K | Heart | 3 | 73.03 | 23.49 | 73.05 | 30.07 | 20.00 | 60.53 | 74.37 |
| HyenaDNA-32K | Heart | 4 | 77.75 | 27.30 | 75.82 | 33.33 | 22.64 | 63.16 | 77.16 |
| HyenaDNA-32K | Heart | 5 | 75.77 | 22.68 | 72.54 | 32.30 | 21.14 | 68.42 | 72.98 |
| HyenaDNA-32K | Heart | Mean (95% CI) | 75.16 (72.81–77.51) | 24.05 (21.66–26.43) | 71.89 (68.22–75.55) | 31.33 (29.34–33.32) | 20.52 (18.75–22.28) | 66.84 (59.32–74.36) | 72.42 (67.75–77.10) |
| HyenaDNA-32K | Face | 1 | 47.44 | 8.91 | 63.57 | 16.18 | 10.07 | 41.18 | 65.66 |
| HyenaDNA-32K | Face | 2 | 55.25 | 9.92 | 58.29 | 18.63 | 11.18 | 55.88 | 58.52 |
| HyenaDNA-32K | Face | 3 | 51.69 | 8.97 | 56.28 | 17.92 | 10.67 | 55.88 | 56.32 |
| HyenaDNA-32K | Face | 4 | 55.09 | 9.82 | 49.75 | 18.70 | 10.85 | 67.65 | 48.08 |
| HyenaDNA-32K | Face | 5 | 59.90 | 14.62 | 65.33 | 20.69 | 12.86 | 52.94 | 66.48 |
| HyenaDNA-32K | Face | Mean (95% CI) | 53.87 (48.12–59.63) | 10.45 (7.50–13.40) | 58.64 (50.95–66.34) | 18.43 (16.41–20.44) | 11.13 (9.82–12.43) | 54.71 (42.99–66.43) | 59.01 (49.66–68.37) |
| HyenaDNA-32K | Limb | 1 | 55.45 | 15.41 | 51.26 | 21.14 | 12.62 | 65.00 | 49.72 |
| HyenaDNA-32K | Limb | 2 | 54.54 | 12.89 | 51.26 | 21.14 | 12.62 | 65.00 | 49.72 |
| HyenaDNA-32K | Limb | 3 | 53.49 | 13.87 | 44.47 | 20.79 | 12.13 | 72.50 | 41.34 |
| HyenaDNA-32K | Limb | 4 | 56.37 | 11.82 | 51.51 | 20.58 | 12.32 | 62.50 | 50.28 |
| HyenaDNA-32K | Limb | 5 | 53.55 | 11.91 | 53.77 | 21.37 | 12.89 | 62.50 | 52.79 |
| HyenaDNA-32K | Limb | Mean (95% CI) | 54.68 (53.14–56.22) | 13.18 (11.32–15.04) | 50.45 (46.10–54.81) | 21.00 (20.61–21.39) | 12.52 (12.15–12.88) | 65.50 (60.40–70.60) | 48.77 (43.38–54.16) |
| HyenaDNA-450K | Forebrain | 1 | 67.33 | 28.34 | 61.96 | 39.36 | 27.07 | 72.06 | 59.88 |
| HyenaDNA-450K | Forebrain | 2 | 69.92 | 34.26 | 71.28 | 40.62 | 31.45 | 57.35 | 74.16 |
| HyenaDNA-450K | Forebrain | 3 | 69.79 | 31.61 | 70.03 | 40.80 | 30.83 | 60.29 | 72.04 |
| HyenaDNA-450K | Forebrain | 4 | 66.54 | 28.47 | 69.77 | 42.31 | 31.43 | 64.71 | 70.82 |
| HyenaDNA-450K | Forebrain | 5 | 67.94 | 30.62 | 72.80 | 42.55 | 33.33 | 58.82 | 75.68 |
| HyenaDNA-450K | Forebrain | Mean (95% CI) | 68.30 (66.44–70.17) | 30.66 (27.62–33.70) | 69.17 (63.95–74.39) | 41.13 (39.49–42.76) | 30.82 (27.97–33.68) | 62.65 (55.27–70.02) | 70.52 (62.77–78.26) |
| HyenaDNA-450K | Midbrain | 1 | 61.12 | 20.44 | 49.87 | 30.66 | 19.21 | 75.86 | 45.43 |
| HyenaDNA-450K | Midbrain | 2 | 57.76 | 19.84 | 63.22 | 29.81 | 20.67 | 53.45 | 64.90 |
| HyenaDNA-450K | Midbrain | 3 | 64.32 | 28.82 | 57.68 | 32.80 | 21.35 | 70.69 | 55.46 |
| HyenaDNA-450K | Midbrain | 4 | 65.40 | 25.84 | 56.68 | 32.28 | 20.92 | 70.69 | 54.28 |
| HyenaDNA-450K | Midbrain | 5 | 57.51 | 20.81 | 67.25 | 30.85 | 22.31 | 50.00 | 70.21 |
| HyenaDNA-450K | Midbrain | Mean (95% CI) | 61.22 (56.71–65.73) | 23.15 (18.22–28.08) | 58.94 (50.70–67.19) | 31.28 (29.75–32.81) | 20.89 (19.49–22.29) | 64.14 (49.75–78.53) | 58.05 (46.03–70.07) |
| HyenaDNA-450K | Hindbrain | 1 | 59.11 | 17.99 | 64.48 | 26.18 | 17.99 | 48.08 | 66.96 |
| HyenaDNA-450K | Hindbrain | 2 | 61.71 | 23.69 | 59.70 | 30.43 | 19.66 | 67.31 | 58.55 |
| HyenaDNA-450K | Hindbrain | 3 | 60.64 | 19.27 | 65.49 | 27.51 | 18.98 | 50.00 | 67.83 |
| HyenaDNA-450K | Hindbrain | 4 | 58.68 | 17.40 | 54.66 | 26.83 | 17.01 | 63.46 | 53.33 |
| HyenaDNA-450K | Hindbrain | 5 | 60.23 | 20.77 | 64.74 | 30.00 | 20.27 | 57.69 | 65.80 |
| HyenaDNA-450K | Hindbrain | Mean (95% CI) | 60.07 (58.57–61.58) | 19.82 (16.70–22.95) | 61.81 (56.10–67.53) | 28.19 (25.81–30.57) | 18.78 (17.16–20.40) | 57.31 (46.98–67.63) | 62.49 (54.67–70.31) |
| HyenaDNA-450K | Neural_Tube | 1 | 57.40 | 14.36 | 66.50 | 25.70 | 17.04 | 52.27 | 68.27 |
| HyenaDNA-450K | Neural_Tube | 2 | 62.57 | 18.20 | 59.45 | 26.48 | 16.57 | 65.91 | 58.64 |
| HyenaDNA-450K | Neural_Tube | 3 | 61.30 | 15.06 | 61.96 | 26.34 | 16.77 | 61.36 | 62.04 |
| HyenaDNA-450K | Neural_Tube | 4 | 61.09 | 15.05 | 58.19 | 24.55 | 15.34 | 61.36 | 57.79 |
| HyenaDNA-450K | Neural_Tube | 5 | 63.69 | 19.29 | 59.45 | 27.15 | 16.95 | 68.18 | 58.36 |
| HyenaDNA-450K | Neural_Tube | Mean (95% CI) | 61.21 (58.26–64.16) | 16.39 (13.66–19.12) | 61.11 (57.00–65.22) | 26.04 (24.82–27.26) | 16.53 (15.68–17.39) | 61.82 (54.25–69.39) | 61.02 (55.58–66.46) |
| HyenaDNA-450K | Heart | 1 | 77.42 | 22.43 | 66.25 | 31.63 | 19.62 | 81.58 | 64.62 |
| HyenaDNA-450K | Heart | 2 | 70.79 | 19.95 | 59.45 | 27.80 | 16.76 | 81.58 | 57.10 |
| HyenaDNA-450K | Heart | 3 | 79.52 | 26.10 | 67.00 | 30.69 | 19.21 | 76.32 | 66.02 |
| HyenaDNA-450K | Heart | 4 | 75.49 | 22.41 | 67.00 | 29.95 | 18.79 | 73.68 | 66.30 |
| HyenaDNA-450K | Heart | 5 | 76.54 | 21.60 | 66.25 | 31.63 | 19.62 | 81.58 | 64.62 |
| HyenaDNA-450K | Heart | Mean (95% CI) | 75.95 (71.92–79.98) | 22.50 (19.70–25.30) | 65.19 (61.18–69.20) | 30.34 (28.37–32.31) | 18.80 (17.32–20.28) | 78.95 (74.33–83.57) | 63.73 (59.03–68.43) |
| HyenaDNA-450K | Face | 1 | 58.60 | 12.38 | 65.33 | 20.69 | 12.86 | 52.94 | 66.48 |
| HyenaDNA-450K | Face | 2 | 58.79 | 13.54 | 65.83 | 22.73 | 14.08 | 58.82 | 66.48 |
| HyenaDNA-450K | Face | 3 | 59.93 | 11.28 | 49.25 | 20.47 | 11.82 | 76.47 | 46.70 |
| HyenaDNA-450K | Face | 4 | 62.21 | 14.81 | 62.81 | 22.11 | 13.46 | 61.76 | 62.91 |
| HyenaDNA-450K | Face | 5 | 57.85 | 11.49 | 61.56 | 20.73 | 12.58 | 58.82 | 61.81 |
| HyenaDNA-450K | Face | Mean (95% CI) | 59.47 (57.36–61.59) | 12.70 (10.87–14.53) | 60.95 (52.54–69.37) | 21.34 (20.09–22.60) | 12.96 (11.89–14.03) | 61.76 (50.81–72.72) | 60.88 (50.70–71.06) |
| HyenaDNA-450K | Limb | 1 | 62.40 | 17.62 | 59.80 | 24.53 | 15.12 | 65.00 | 59.22 |
| HyenaDNA-450K | Limb | 2 | 59.90 | 15.13 | 64.82 | 23.08 | 14.79 | 52.50 | 66.20 |
| HyenaDNA-450K | Limb | 3 | 53.92 | 12.44 | 61.81 | 20.00 | 12.67 | 47.50 | 63.41 |
| HyenaDNA-450K | Limb | 4 | 59.73 | 14.05 | 67.59 | 22.75 | 14.96 | 47.50 | 69.83 |
| HyenaDNA-450K | Limb | 5 | 54.63 | 12.20 | 52.26 | 20.83 | 12.50 | 62.50 | 51.12 |
| HyenaDNA-450K | Limb | Mean (95% CI) | 58.12 (53.56–62.67) | 14.29 (11.54–17.04) | 61.26 (54.01–68.50) | 22.24 (19.98–24.49) | 14.01 (12.39–15.63) | 55.00 (44.70–65.30) | 61.96 (53.02–70.89) |
| HyenaDNA-1M | Forebrain | 1 | 67.98 | 30.40 | 68.77 | 39.81 | 29.71 | 60.29 | 70.52 |
| HyenaDNA-1M | Forebrain | 2 | 68.50 | 29.94 | 65.74 | 40.35 | 28.75 | 67.65 | 65.35 |
| HyenaDNA-1M | Forebrain | 3 | 55.58 | 21.93 | 50.63 | 30.50 | 20.09 | 63.24 | 48.02 |
| HyenaDNA-1M | Forebrain | 4 | 67.23 | 29.92 | 65.24 | 38.94 | 27.85 | 64.71 | 65.35 |
| HyenaDNA-1M | Forebrain | 5 | 68.04 | 28.86 | 62.72 | 37.29 | 26.19 | 64.71 | 62.31 |
| HyenaDNA-1M | Forebrain | Mean (95% CI) | 65.47 (58.58–72.35) | 28.21 (23.79–32.63) | 62.62 (53.88–71.36) | 37.38 (32.39–42.36) | 26.52 (21.78–31.26) | 64.12 (60.80–67.43) | 62.31 (51.74–72.88) |
| HyenaDNA-1M | Midbrain | 1 | 51.06 | 17.81 | 46.10 | 24.65 | 15.49 | 60.34 | 43.66 |
| HyenaDNA-1M | Midbrain | 2 | 57.72 | 19.32 | 58.94 | 30.64 | 20.34 | 62.07 | 58.41 |
| HyenaDNA-1M | Midbrain | 3 | 60.51 | 21.19 | 60.96 | 31.11 | 20.96 | 60.34 | 61.06 |
| HyenaDNA-1M | Midbrain | 4 | 63.40 | 22.40 | 66.75 | 31.25 | 22.39 | 51.72 | 69.32 |
| HyenaDNA-1M | Midbrain | 5 | 61.48 | 24.12 | 59.70 | 29.20 | 19.64 | 56.90 | 60.18 |
| HyenaDNA-1M | Midbrain | Mean (95% CI) | 58.84 (52.87–64.80) | 20.97 (17.88–24.05) | 58.49 (49.08–67.90) | 29.37 (25.94–32.80) | 19.76 (16.54–22.99) | 58.28 (53.16–63.39) | 58.53 (46.96–70.09) |
| HyenaDNA-1M | Hindbrain | 1 | 63.12 | 23.28 | 63.48 | 31.28 | 20.75 | 63.46 | 63.48 |
| HyenaDNA-1M | Hindbrain | 2 | 51.97 | 16.80 | 67.76 | 23.81 | 17.24 | 38.46 | 72.17 |
| HyenaDNA-1M | Hindbrain | 3 | 61.14 | 20.38 | 64.99 | 27.98 | 19.15 | 51.92 | 66.96 |
| HyenaDNA-1M | Hindbrain | 4 | 70.27 | 26.30 | 59.70 | 32.77 | 20.97 | 75.00 | 57.39 |
| HyenaDNA-1M | Hindbrain | 5 | 53.01 | 18.14 | 47.10 | 22.22 | 13.76 | 57.69 | 45.51 |
| HyenaDNA-1M | Hindbrain | Mean (95% CI) | 59.90 (50.49–69.31) | 20.98 (16.19–25.77) | 60.60 (50.56–70.65) | 27.61 (21.93–33.29) | 18.37 (14.67–22.08) | 57.31 (40.47–74.14) | 61.10 (48.39–73.82) |
| HyenaDNA-1M | Neural_Tube | 1 | 56.86 | 13.46 | 54.16 | 23.53 | 14.43 | 63.64 | 52.97 |
| HyenaDNA-1M | Neural_Tube | 2 | 59.49 | 14.00 | 55.16 | 24.58 | 15.10 | 65.91 | 53.82 |
| HyenaDNA-1M | Neural_Tube | 3 | 61.76 | 13.60 | 50.63 | 26.32 | 15.77 | 79.55 | 47.03 |
| HyenaDNA-1M | Neural_Tube | 4 | 54.94 | 14.98 | 49.87 | 21.96 | 13.27 | 63.64 | 48.16 |
| HyenaDNA-1M | Neural_Tube | 5 | 58.51 | 14.42 | 57.43 | 22.83 | 14.29 | 56.82 | 57.51 |
| HyenaDNA-1M | Neural_Tube | Mean (95% CI) | 58.31 (55.10–61.53) | 14.09 (13.32–14.87) | 53.45 (49.52–57.38) | 23.84 (21.75–25.93) | 14.57 (13.41–15.73) | 65.91 (55.54–76.28) | 51.90 (46.56–57.24) |
| HyenaDNA-1M | Heart | 1 | 68.27 | 19.49 | 62.22 | 26.47 | 16.27 | 71.05 | 61.28 |
| HyenaDNA-1M | Heart | 2 | 72.23 | 20.58 | 67.51 | 30.27 | 19.05 | 73.68 | 66.85 |
| HyenaDNA-1M | Heart | 3 | 75.45 | 22.95 | 69.52 | 30.86 | 19.71 | 71.05 | 69.36 |
| HyenaDNA-1M | Heart | 4 | 67.38 | 16.30 | 68.01 | 27.43 | 17.52 | 63.16 | 68.52 |
| HyenaDNA-1M | Heart | 5 | 73.86 | 24.14 | 67.25 | 30.85 | 19.33 | 76.32 | 66.30 |
| HyenaDNA-1M | Heart | Mean (95% CI) | 71.44 (67.09–75.79) | 20.70 (16.88–24.51) | 66.90 (63.47–70.33) | 29.18 (26.60–31.75) | 18.37 (16.58–20.17) | 71.05 (64.94–77.17) | 66.46 (62.55–70.37) |
| HyenaDNA-1M | Face | 1 | 59.53 | 12.84 | 53.52 | 18.50 | 10.88 | 61.76 | 52.75 |
| HyenaDNA-1M | Face | 2 | 62.79 | 12.18 | 64.32 | 22.83 | 14.00 | 61.76 | 64.56 |
| HyenaDNA-1M | Face | 3 | 59.29 | 11.45 | 52.51 | 20.25 | 11.82 | 70.59 | 50.82 |
| HyenaDNA-1M | Face | 4 | 57.78 | 13.11 | 49.25 | 20.47 | 11.82 | 76.47 | 46.70 |
| HyenaDNA-1M | Face | 5 | 58.30 | 13.65 | 67.09 | 20.61 | 12.98 | 50.00 | 68.68 |
| HyenaDNA-1M | Face | Mean (95% CI) | 59.54 (57.11–61.96) | 12.64 (11.59–13.70) | 57.34 (47.58–67.10) | 20.53 (18.62–22.44) | 12.30 (10.80–13.80) | 64.12 (51.63–76.61) | 56.70 (45.00–68.41) |
| HyenaDNA-1M | Limb | 1 | 59.41 | 16.99 | 68.09 | 24.85 | 16.28 | 52.50 | 69.83 |
| HyenaDNA-1M | Limb | 2 | 52.21 | 13.53 | 54.02 | 20.09 | 12.17 | 57.50 | 53.63 |
| HyenaDNA-1M | Limb | 3 | 59.98 | 13.56 | 61.06 | 23.65 | 14.72 | 60.00 | 61.17 |
| HyenaDNA-1M | Limb | 4 | 52.68 | 11.10 | 52.76 | 21.01 | 12.63 | 62.50 | 51.68 |
| HyenaDNA-1M | Limb | 5 | 54.11 | 12.78 | 51.51 | 21.86 | 13.04 | 67.50 | 49.72 |
| HyenaDNA-1M | Limb | Mean (95% CI) | 55.68 (51.04–60.32) | 13.59 (10.93–16.26) | 57.49 (48.81–66.16) | 22.29 (19.88–24.70) | 13.77 (11.65–15.88) | 60.00 (53.06–66.94) | 57.21 (46.92–67.49) |
| DeepSTARR | Forebrain | 1 | 58.50 | 21.39 | 52.64 | 30.37 | 20.30 | 60.29 | 51.06 |
| DeepSTARR | Forebrain | 2 | 54.09 | 19.50 | 59.19 | 30.17 | 21.34 | 51.47 | 60.79 |
| DeepSTARR | Forebrain | 3 | 58.35 | 26.90 | 57.18 | 29.75 | 20.69 | 52.94 | 58.05 |
| DeepSTARR | Forebrain | 4 | 57.07 | 20.69 | 53.15 | 32.61 | 21.63 | 66.18 | 50.46 |
| DeepSTARR | Forebrain | 5 | 52.08 | 17.58 | 59.19 | 28.95 | 20.62 | 48.53 | 61.40 |
| DeepSTARR | Forebrain | Mean (95% CI) | 56.02 (52.51–59.53) | 21.21 (16.87–25.55) | 56.27 (52.31–60.24) | 30.37 (28.67–32.07) | 20.92 (20.23–21.60) | 55.88 (46.94–64.83) | 56.35 (49.82–62.89) |
| DeepSTARR | Midbrain | 1 | 51.23 | 15.97 | 47.61 | 25.71 | 16.22 | 62.07 | 45.13 |
| DeepSTARR | Midbrain | 2 | 50.02 | 14.31 | 59.95 | 23.19 | 16.11 | 41.38 | 63.13 |
| DeepSTARR | Midbrain | 3 | 53.36 | 19.28 | 62.22 | 25.74 | 18.06 | 44.83 | 65.19 |
| DeepSTARR | Midbrain | 4 | 49.48 | 14.64 | 50.88 | 24.12 | 15.58 | 53.45 | 50.44 |
| DeepSTARR | Midbrain | 5 | 50.07 | 16.19 | 64.23 | 24.47 | 17.69 | 39.66 | 68.44 |
| DeepSTARR | Midbrain | Mean (95% CI) | 50.83 (48.91–52.76) | 16.08 (13.63–18.52) | 56.98 (47.90–66.06) | 24.65 (23.29–26.00) | 16.73 (15.39–18.07) | 48.28 (36.65–59.90) | 58.47 (45.92–71.01) |
| DeepSTARR | Hindbrain | 1 | 53.58 | 14.12 | 52.90 | 24.90 | 15.74 | 59.62 | 51.88 |
| DeepSTARR | Hindbrain | 2 | 51.06 | 13.56 | 56.68 | 23.21 | 15.12 | 50.00 | 57.68 |
| DeepSTARR | Hindbrain | 3 | 45.32 | 14.90 | 58.69 | 18.00 | 12.16 | 34.62 | 62.32 |
| DeepSTARR | Hindbrain | 4 | 51.15 | 16.53 | 57.68 | 23.64 | 15.48 | 50.00 | 58.84 |
| DeepSTARR | Hindbrain | 5 | 45.15 | 11.70 | 51.89 | 20.75 | 13.23 | 48.08 | 52.46 |
| DeepSTARR | Hindbrain | Mean (95% CI) | 49.25 (44.53–53.98) | 14.16 (11.96–16.36) | 55.57 (51.84–59.30) | 22.10 (18.69–25.50) | 14.34 (12.40–16.29) | 48.46 (37.34–59.58) | 56.64 (51.15–62.13) |
| DeepSTARR | Neural_Tube | 1 | 52.11 | 11.60 | 60.71 | 20.41 | 13.16 | 45.45 | 62.61 |
| DeepSTARR | Neural_Tube | 2 | 51.21 | 13.87 | 44.84 | 19.19 | 11.45 | 59.09 | 43.06 |
| DeepSTARR | Neural_Tube | 3 | 62.32 | 17.87 | 61.71 | 24.75 | 15.82 | 56.82 | 62.32 |
| DeepSTARR | Neural_Tube | 4 | 56.17 | 15.52 | 58.19 | 23.85 | 14.94 | 59.09 | 58.07 |
| DeepSTARR | Neural_Tube | 5 | 53.14 | 12.01 | 60.96 | 22.11 | 14.19 | 50.00 | 62.32 |
| DeepSTARR | Neural_Tube | Mean (95% CI) | 54.99 (49.40–60.58) | 14.17 (10.95–17.39) | 57.28 (48.49–66.07) | 22.06 (19.19–24.94) | 13.91 (11.82–16.01) | 54.09 (46.52–61.66) | 57.68 (47.26–68.09) |
| DeepSTARR | Heart | 1 | 54.44 | 13.20 | 67.76 | 20.99 | 13.71 | 44.74 | 70.19 |
| DeepSTARR | Heart | 2 | 55.98 | 14.61 | 67.76 | 21.95 | 14.29 | 47.37 | 69.92 |
| DeepSTARR | Heart | 3 | 60.37 | 13.45 | 57.43 | 22.12 | 13.41 | 63.16 | 56.82 |
| DeepSTARR | Heart | 4 | 67.20 | 14.89 | 63.22 | 26.26 | 16.25 | 68.42 | 62.67 |
| DeepSTARR | Heart | 5 | 58.60 | 15.67 | 50.63 | 19.67 | 11.65 | 63.16 | 49.30 |
| DeepSTARR | Heart | Mean (95% CI) | 59.32 (53.15–65.48) | 14.36 (13.09–15.64) | 61.36 (52.24–70.48) | 22.20 (19.13–25.27) | 13.86 (11.80–15.92) | 57.37 (44.22–70.52) | 61.78 (50.71–72.85) |
| DeepSTARR | Face | 1 | 51.68 | 14.17 | 57.79 | 17.65 | 10.59 | 52.94 | 58.24 |
| DeepSTARR | Face | 2 | 67.15 | 18.15 | 59.80 | 21.57 | 12.94 | 64.71 | 59.34 |
| DeepSTARR | Face | 3 | 51.87 | 8.78 | 47.74 | 17.46 | 10.09 | 64.71 | 46.15 |
| DeepSTARR | Face | 4 | 54.20 | 8.90 | 48.74 | 20.31 | 11.71 | 76.47 | 46.15 |
| DeepSTARR | Face | 5 | 57.94 | 10.46 | 66.83 | 20.48 | 12.88 | 50.00 | 68.41 |
| DeepSTARR | Face | Mean (95% CI) | 56.57 (48.58–64.56) | 12.09 (7.09–17.09) | 56.18 (46.25–66.11) | 19.49 (17.21–21.77) | 11.64 (10.03–13.25) | 61.76 (48.60–74.93) | 55.66 (43.82–67.49) |
| DeepSTARR | Limb | 1 | 57.03 | 16.29 | 66.83 | 24.14 | 15.67 | 52.50 | 68.44 |
| DeepSTARR | Limb | 2 | 59.87 | 14.46 | 53.02 | 22.41 | 13.43 | 67.50 | 51.40 |
| DeepSTARR | Limb | 3 | 47.25 | 10.42 | 62.31 | 18.48 | 11.81 | 42.50 | 64.53 |
| DeepSTARR | Limb | 4 | 57.41 | 12.93 | 57.29 | 21.30 | 13.07 | 57.50 | 57.26 |
| DeepSTARR | Limb | 5 | 49.77 | 10.63 | 49.50 | 18.62 | 11.11 | 57.50 | 48.60 |
| DeepSTARR | Limb | Mean (95% CI) | 54.27 (47.51–61.02) | 12.95 (9.83–16.07) | 57.79 (49.14–66.44) | 20.99 (17.95–24.02) | 13.02 (10.84–15.20) | 55.50 (44.22–66.78) | 58.04 (47.58–68.51) |

**Supplementary Table 3 | Production tissue oracles: checkpoint selection and held-out test performance.** Performance and deployment details for the seven tissue-specific Enformer classifiers used as REGAL reward oracles and genome-wide scanners.

| **Tissue** | **Selection_metric** | **Seed** | **Production_checkpoint** | **Test_positives** | **Test_negatives** | **Test_positive_to_negative_ratio** | **Test_AUROC** | **Test_AUPRC** | **Decision_threshold** | **Accuracy** | **Sensitivity** | **Specificity** |
| --- | --- | --- | --- | --- | --- | --- | --- | --- | --- | --- | --- | --- |
| Forebrain | Validation AUPRC | 999 | best_pr_s999.pth | 68 | 329 | 1:4.8 | 0.805 | 0.575 | 0.0208 | 0.793 | 0.735 | 0.805 |
| Midbrain | Validation AUPRC | 999 | best_pr_s999.pth | 58 | 339 | 1:5.8 | 0.735 | 0.422 | 0.0125 | 0.733 | 0.638 | 0.749 |
| Hindbrain | Validation AUROC | 123 | best_roc_s123.pth | 52 | 345 | 1:6.6 | 0.779 | 0.388 | 0.1755 | 0.743 | 0.712 | 0.748 |
| Neural_Tube | Validation AUPRC | 42 | best_pr_s42.pth | 44 | 353 | 1:8.0 | 0.720 | 0.275 | 0.3917 | 0.796 | 0.636 | 0.816 |
| Heart | Validation AUPRC | 456 | best_pr_s456.pth | 38 | 359 | 1:9.4 | 0.788 | 0.394 | 0.3152 | 0.856 | 0.579 | 0.886 |
| Face | Validation AUPRC | 42 | best_pr_s42.pth | 34 | 364 | 1:10.7 | 0.730 | 0.273 | 0.4924 | 0.706 | 0.706 | 0.706 |
| Limb | Validation AUPRC | 999 | best_pr_s999.pth | 40 | 358 | 1:8.9 | 0.864 | 0.527 | 0.6367 | 0.849 | 0.725 | 0.863 |

**Supplementary Table 4 | ENCODE ChromHMM chromatin-state annotations used for the enhancer-state enrichment analysis.**

| **Tissue** | **ENCODE File ID** | **ChromHMM Regions** |
| --- | --- | --- |
| Face | ENCFF128LXT | 549,885 |
| Heart | ENCFF497ATP | 713,406 |
| Forebrain | ENCFF749VMS | 532,682 |
| Hindbrain | ENCFF730BWG | 492,973 |
| Limb | ENCFF722AXW | 574,732 |
| Midbrain | ENCFF343BKP | 553,731 |
| Neural_Tube | ENCFF872QGN | 510,658 |

**Supplementary Table 5 | Components of the four REGAL design campaigns.** The four interchangeable components of the REGAL framework, namely the motif vocabulary, the reward oracle, the background pool, and the target set, together with the window of the composite sequence each oracle reads, as instantiated for the mouse, *Drosophila*, human cell-line, and zebrafish campaigns.

| **Campaign** | **Motif library** | **Reward oracle** | **Background pool** | **Target set** |
| --- | --- | --- | --- | --- |
| Mouse | De novo TF-MoDISco vocabulary, 637 motifs | Per-tissue Enformer classifier | Real VISTA-negative sequences for the matching tissue (500 sampled per run) | Six mouse tissues: forebrain, midbrain, hindbrain, neural tube, heart, limb |
| *Drosophila* | jaspar_ffs_v1, 943 motifs | DeepSTARR_embryo | Per-tissue background pools of tissue-inactive DeepSTARR genomic tiles (1,000 sequences each, stored at 1,536 bp) | Central nervous system, gut, muscle |
| Human cell lines | jaspar_vertebrates_2024, 879 PWMs | Malinois | Gosai et al. 2024 MPRA low-activity oligos (\|log_2_FC\| < 0.1 and lfcSE < 0.3) | K562, HepG2, SK-N-SH |
| Zebrafish | zebrafish_swissregulon_jaspar_merged_2026, 1,246 motifs | DeepZebrafish | Per-cell-type, per-timepoint 1,536-bp pools | Cardiac muscle, endothelial, fast muscle, slow muscle, notochord |

**Supplementary Table 6 | Training objectives and optimization settings for REGAL design campaigns.** Summary of the reward objectives, off-target penalties, random seeds, training iterations, and campaign-specific optimizer settings for the four REGAL design campaigns.

| **Campaign** | **Seed** | **Iterations** | **Objective** | **Off-target definition** | **Weighting / special rule** |
| --- | --- | --- | --- | --- | --- |
| Mouse | 1–19, 42, 123, 999, 2024, 31337 | 1,500 | $R=\max\left( d_{\mathrm{target}} , 0 \right)-\sum_{i} w_{i}d_{i}$ | Neural targets: heart and limb; heart and limb targets: each other and all neural tissues. | Neural: heart and limb, $w=1.5$; heart↔limb, $w=1.0$; neural tissues for heart/limb, $w=0.25$. Neural siblings were unpenalized. |
| *Drosophila* | 1–19, 42, 123, 999, 2024, 31337 | 3,000 | $R=d_{\mathrm{target}}-\alpha P_{\mathrm{off}}$ | All remaining embryonic tissue outputs; for CNS targets, the brain output was excluded because it is nested within the CNS output. | $\alpha=0.5$ for CNS and gut; $\alpha=1.0$ for muscle. |
| Human cell lines | 1, 2, 3, 42, 123, 999, 2024, 31337 | 1,500 | $R=d_{\mathrm{target}}-\alpha\max\left( d_{\mathrm{off}} \right)$ | The two non-target Malinois cell-line outputs. | $\alpha=1.0$. |
| Zebrafish | 42, 123, 456 | 5,000 | $R=d_{\mathrm{target}}-\alpha\text{ }\mathrm{mean}\left( \mathrm{top}\text{ }10\text{ }d_{\mathrm{off}} \right)$ | All 639 off-target heads, excluding neutral heads within the same group. | $\alpha=1.0$ for endothelial and slow muscle; $\alpha=2.0$ for cardiac muscle, fast muscle and notochord. |

**Supplementary Table 7 | Oligos used in this study.** Primers and adapters used for STARR-seq library preparation, sequencing, and H11 transgene genotyping, grouped by function.

| **name** | **sequence_5to3** |
| --- | --- |
| Fwd_universal_STARR | CATCTCCGAGCCCACGAGACTCGACGAATTCGGCCGG |
| Rev_N504_STARR | ACATCTGACGCTGCCGACGATCTACTCTTGCATGCTCTAGATCAATCTAATTC |
| Tn5_1 | TCGTCGGCAGCGTCAGATGTGTATAAGAGACAG |
| Tn5_2_ME_comp | GTCTCGTGGGCTCGGAGATGTGTATAAGAGACAG |
| TN5MEREV | /5Phos/CTGTCTCTTATACACATCT |
| Fwd_atac-starr_tag | TCGTCGGCAGCGTCAGATG |
| Rev_atac-starr_tag | GTCTCGTGGGCTCGGAGATG |
| STARR_GSP | CTCATCAATGTATCTTATCATGTCTG |
| Illumina_P5_primer | AATGATACGGCGACCACCGAGATCTACAC |
| Illumina_P7_primer | CAAGCAGAAGACGGCATACGAGAT |
| Nextera_i7_N701_Ad2.1 | CAAGCAGAAGACGGCATACGAGATTCGCCTTAGTCTCGTGGGCTCGGAGATGTG |
| Nextera_i7_N702_Ad2.2 | CAAGCAGAAGACGGCATACGAGATCTAGTACGGTCTCGTGGGCTCGGAGATGTG |
| Nextera_i7_N703_Ad2.3 | CAAGCAGAAGACGGCATACGAGATTTCTGCCTGTCTCGTGGGCTCGGAGATGTG |
| Nextera_i7_N704_Ad2.4 | CAAGCAGAAGACGGCATACGAGATGCTCAGGAGTCTCGTGGGCTCGGAGATGTG |
| Nextera_i7_N705_Ad2.5 | CAAGCAGAAGACGGCATACGAGATAGGAGTCCGTCTCGTGGGCTCGGAGATGTG |
| Nextera_i5_N504_Ad1.4 | AATGATACGGCGACCACCGAGATCTACACAGAGTAGATCGTCGGCAGCGTCAGATGTGTAT |
| Nextera_i5_N505_Ad1.5 | AATGATACGGCGACCACCGAGATCTACACGTAAGGAGTCGTCGGCAGCGTCAGATGTGTAT |
| Nextera_i5_N506_Ad1.6 | AATGATACGGCGACCACCGAGATCTACACACTGCATATCGTCGGCAGCGTCAGATGTGTAT |
| Nextera_i5_N507_Ad1.7 | AATGATACGGCGACCACCGAGATCTACACAAGGAGTATCGTCGGCAGCGTCAGATGTGTAT |
| Nextera_i5_N508_Ad1.8 | AATGATACGGCGACCACCGAGATCTACACCTAAGCCTTCGTCGGCAGCGTCAGATGTGTAT |
| H11-5prime-F | AGCTGGTGAGTTTCTGACTGG |
| H11-5prime-R | CTGGAGAGCTTGTGAGACAGG |
| H11-3prime-F | TGCATTCTAGTTGTGGTTTGTCCA |
| H11-3prime-R | ACCTTTGCTCTTGGGGCTTAGA |

**Supplementary Table 8 | Replicate correlation for the mouse *in vitro* STARR-seq designed library.** All-versus-all Pearson correlations among the 12 input-DNA and reporter-RNA libraries from three replicates each in N2a and HL-1 cells. Correlations were calculated from log₁₀(count + 1)-transformed per-oligo counts across the 9,500-sequence library.

| **Sample** | **N2a DNA R1** | **N2a DNA R2** | **N2a DNA R3** | **N2a RNA R1** | **N2a RNA R2** | **N2a RNA R3** | **HL-1 DNA R1** | **HL-1 DNA R2** | **HL-1 DNA R3** | **HL-1 RNA R1** | **HL-1 RNA R2** | **HL-1 RNA R3** |
| --- | --- | --- | --- | --- | --- | --- | --- | --- | --- | --- | --- | --- |
| N2a DNA R1 | 1.0000 | 0.9887 | 0.9885 | 0.8498 | 0.8450 | 0.8522 | 0.9858 | 0.9845 | 0.9841 | 0.6858 | 0.6705 | 0.6660 |
| N2a DNA R2 | 0.9887 | 1.0000 | 0.9904 | 0.8513 | 0.8456 | 0.8545 | 0.9865 | 0.9847 | 0.9857 | 0.6940 | 0.6793 | 0.6762 |
| N2a DNA R3 | 0.9885 | 0.9904 | 1.0000 | 0.8522 | 0.8472 | 0.8540 | 0.9856 | 0.9851 | 0.9849 | 0.6915 | 0.6758 | 0.6734 |
| N2a RNA R1 | 0.8498 | 0.8513 | 0.8522 | 1.0000 | 0.9836 | 0.9877 | 0.8531 | 0.8521 | 0.8529 | 0.7039 | 0.6928 | 0.6892 |
| N2a RNA R2 | 0.8450 | 0.8456 | 0.8472 | 0.9836 | 1.0000 | 0.9848 | 0.8460 | 0.8465 | 0.8464 | 0.6916 | 0.6788 | 0.6742 |
| N2a RNA R3 | 0.8522 | 0.8545 | 0.8540 | 0.9877 | 0.9848 | 1.0000 | 0.8553 | 0.8529 | 0.8570 | 0.7054 | 0.6946 | 0.6901 |
| HL-1 DNA R1 | 0.9858 | 0.9865 | 0.9856 | 0.8531 | 0.8460 | 0.8553 | 1.0000 | 0.9854 | 0.9875 | 0.6931 | 0.6786 | 0.6737 |
| HL-1 DNA R2 | 0.9845 | 0.9847 | 0.9851 | 0.8521 | 0.8465 | 0.8529 | 0.9854 | 1.0000 | 0.9841 | 0.6850 | 0.6694 | 0.6656 |
| HL-1 DNA R3 | 0.9841 | 0.9857 | 0.9849 | 0.8529 | 0.8464 | 0.8570 | 0.9875 | 0.9841 | 1.0000 | 0.6972 | 0.6818 | 0.6774 |
| HL-1 RNA R1 | 0.6858 | 0.6940 | 0.6915 | 0.7039 | 0.6916 | 0.7054 | 0.6931 | 0.6850 | 0.6972 | 1.0000 | 0.9733 | 0.9707 |
| HL-1 RNA R2 | 0.6705 | 0.6793 | 0.6758 | 0.6928 | 0.6788 | 0.6946 | 0.6786 | 0.6694 | 0.6818 | 0.9733 | 1.0000 | 0.9725 |
| HL-1 RNA R3 | 0.6660 | 0.6762 | 0.6734 | 0.6892 | 0.6742 | 0.6901 | 0.6737 | 0.6656 | 0.6774 | 0.9707 | 0.9725 | 1.0000 |

**Supplementary Table 9 | Replicate correlation for the mouse *in vitro* STARR-seq motif-ablation and position-shuffling library.** All-versus-all Pearson correlations among the 12 input-DNA and reporter-RNA libraries from three replicates each in N2a and HL-1 cells, calculated from log₁₀(count + 1)-transformed per-oligo counts across the 3,198 motif-ablation and position-shuffling oligos.

| **Sample** | **N2a DNA R1** | **N2a DNA R2** | **N2a DNA R3** | **N2a RNA R1** | **N2a RNA R2** | **N2a RNA R3** | **HL-1 DNA R1** | **HL-1 DNA R2** | **HL-1 DNA R3** | **HL-1 RNA R1** | **HL-1 RNA R2** | **HL-1 RNA R3** |
| --- | --- | --- | --- | --- | --- | --- | --- | --- | --- | --- | --- | --- |
| N2a DNA R1 | 1.0000 | 0.9874 | 0.9861 | 0.7534 | 0.7548 | 0.7596 | 0.9788 | 0.9825 | 0.9762 | 0.5207 | 0.5155 | 0.5065 |
| N2a DNA R2 | 0.9874 | 1.0000 | 0.9883 | 0.7530 | 0.7536 | 0.7599 | 0.9811 | 0.9816 | 0.9814 | 0.5323 | 0.5258 | 0.5184 |
| N2a DNA R3 | 0.9861 | 0.9883 | 1.0000 | 0.7480 | 0.7517 | 0.7581 | 0.9760 | 0.9787 | 0.9797 | 0.5253 | 0.5176 | 0.5103 |
| N2a RNA R1 | 0.7534 | 0.7530 | 0.7480 | 1.0000 | 0.9902 | 0.9918 | 0.7634 | 0.7697 | 0.7645 | 0.4931 | 0.4938 | 0.4814 |
| N2a RNA R2 | 0.7548 | 0.7536 | 0.7517 | 0.9902 | 1.0000 | 0.9910 | 0.7618 | 0.7709 | 0.7645 | 0.4854 | 0.4844 | 0.4733 |
| N2a RNA R3 | 0.7596 | 0.7599 | 0.7581 | 0.9918 | 0.9910 | 1.0000 | 0.7670 | 0.7740 | 0.7711 | 0.4990 | 0.4991 | 0.4893 |
| HL-1 DNA R1 | 0.9788 | 0.9811 | 0.9760 | 0.7634 | 0.7618 | 0.7670 | 1.0000 | 0.9808 | 0.9819 | 0.5282 | 0.5213 | 0.5122 |
| HL-1 DNA R2 | 0.9825 | 0.9816 | 0.9787 | 0.7697 | 0.7709 | 0.7740 | 0.9808 | 1.0000 | 0.9797 | 0.5225 | 0.5173 | 0.5068 |
| HL-1 DNA R3 | 0.9762 | 0.9814 | 0.9797 | 0.7645 | 0.7645 | 0.7711 | 0.9819 | 0.9797 | 1.0000 | 0.5339 | 0.5289 | 0.5195 |
| HL-1 RNA R1 | 0.5207 | 0.5323 | 0.5253 | 0.4931 | 0.4854 | 0.4990 | 0.5282 | 0.5225 | 0.5339 | 1.0000 | 0.9780 | 0.9780 |
| HL-1 RNA R2 | 0.5155 | 0.5258 | 0.5176 | 0.4938 | 0.4844 | 0.4991 | 0.5213 | 0.5173 | 0.5289 | 0.9780 | 1.0000 | 0.9793 |
| HL-1 RNA R3 | 0.5065 | 0.5184 | 0.5103 | 0.4814 | 0.4733 | 0.4893 | 0.5122 | 0.5068 | 0.5195 | 0.9780 | 0.9793 | 1.0000 |

**Supplementary Table 10 | Replicate correlation for the human *in vitro* STARR-seq library.** All-versus-all Pearson correlations among the 18 input-DNA and reporter-RNA libraries from three replicates each in K562, HepG2 and SK-N-SH cells, calculated from log₁₀(count + 1)-transformed per-oligo counts.

| **Sample** | **K562 DNA R1** | **K562 DNA R2** | **K562 DNA R3** | **K562 RNA R1** | **K562 RNA R2** | **K562 RNA R3** | **HepG2 DNA R1** | **HepG2 DNA R2** | **HepG2 DNA R3** | **HepG2 RNA R1** | **HepG2 RNA R2** | **HepG2 RNA R3** | **SK-N-SH DNA R1** | **SK-N-SH DNA R2** | **SK-N-SH DNA R3** | **SK-N-SH RNA R1** | **SK-N-SH RNA R2** | **SK-N-SH RNA R3** |
| --- | --- | --- | --- | --- | --- | --- | --- | --- | --- | --- | --- | --- | --- | --- | --- | --- | --- | --- |
| K562 DNA R1 | 1 | 0.9953 | 0.9934 | 0.8112 | 0.7988 | 0.7929 | 0.9789 | 0.9804 | 0.9775 | 0.7287 | 0.7289 | 0.7295 | 0.9864 | 0.9947 | 0.9858 | 0.6446 | 0.6419 | 0.6144 |
| K562 DNA R2 | 0.9953 | 1 | 0.9927 | 0.808 | 0.7962 | 0.7906 | 0.9787 | 0.98 | 0.9777 | 0.7271 | 0.7266 | 0.728 | 0.9852 | 0.9951 | 0.9844 | 0.6419 | 0.6398 | 0.6122 |
| K562 DNA R3 | 0.9934 | 0.9927 | 1 | 0.8154 | 0.8027 | 0.7973 | 0.9775 | 0.9805 | 0.9771 | 0.7282 | 0.7292 | 0.7287 | 0.9839 | 0.9916 | 0.9843 | 0.6451 | 0.6414 | 0.6136 |
| K562 RNA R1 | 0.8112 | 0.808 | 0.8154 | 1 | 0.9948 | 0.9938 | 0.8216 | 0.8246 | 0.8219 | 0.633 | 0.635 | 0.6323 | 0.8283 | 0.8095 | 0.8294 | 0.6263 | 0.6239 | 0.6118 |
| K562 RNA R2 | 0.7988 | 0.7962 | 0.8027 | 0.9948 | 1 | 0.9948 | 0.8081 | 0.8105 | 0.808 | 0.621 | 0.6224 | 0.6202 | 0.8163 | 0.797 | 0.8176 | 0.6176 | 0.6166 | 0.6051 |
| K562 RNA R3 | 0.7929 | 0.7906 | 0.7973 | 0.9938 | 0.9948 | 1 | 0.8021 | 0.8051 | 0.8017 | 0.6147 | 0.6169 | 0.6134 | 0.8099 | 0.7909 | 0.8121 | 0.6136 | 0.6119 | 0.6013 |
| HepG2 DNA R1 | 0.9789 | 0.9787 | 0.9775 | 0.8216 | 0.8081 | 0.8021 | 1 | 0.9965 | 0.9963 | 0.7649 | 0.7651 | 0.7647 | 0.9842 | 0.9802 | 0.9863 | 0.6486 | 0.6474 | 0.6286 |
| HepG2 DNA R2 | 0.9804 | 0.98 | 0.9805 | 0.8246 | 0.8105 | 0.8051 | 0.9965 | 1 | 0.9966 | 0.7629 | 0.7642 | 0.7636 | 0.9841 | 0.9817 | 0.9872 | 0.6508 | 0.6496 | 0.6298 |
| HepG2 DNA R3 | 0.9775 | 0.9777 | 0.9771 | 0.8219 | 0.808 | 0.8017 | 0.9963 | 0.9966 | 1 | 0.7678 | 0.7683 | 0.7698 | 0.9834 | 0.9793 | 0.9863 | 0.6492 | 0.6485 | 0.6293 |
| HepG2 RNA R1 | 0.7287 | 0.7271 | 0.7282 | 0.633 | 0.621 | 0.6147 | 0.7649 | 0.7629 | 0.7678 | 1 | 0.9967 | 0.9969 | 0.7626 | 0.7306 | 0.7633 | 0.801 | 0.8041 | 0.8066 |
| HepG2 RNA R2 | 0.7289 | 0.7266 | 0.7292 | 0.635 | 0.6224 | 0.6169 | 0.7651 | 0.7642 | 0.7683 | 0.9967 | 1 | 0.997 | 0.7627 | 0.7298 | 0.7635 | 0.8008 | 0.8037 | 0.807 |
| HepG2 RNA R3 | 0.7295 | 0.728 | 0.7287 | 0.6323 | 0.6202 | 0.6134 | 0.7647 | 0.7636 | 0.7698 | 0.9969 | 0.997 | 1 | 0.7639 | 0.7315 | 0.764 | 0.7991 | 0.8024 | 0.8049 |
| SK-N-SH DNA R1 | 0.9864 | 0.9852 | 0.9839 | 0.8283 | 0.8163 | 0.8099 | 0.9842 | 0.9841 | 0.9834 | 0.7626 | 0.7627 | 0.7639 | 1 | 0.9863 | 0.9945 | 0.658 | 0.6562 | 0.6374 |
| SK-N-SH DNA R2 | 0.9947 | 0.9951 | 0.9916 | 0.8095 | 0.797 | 0.7909 | 0.9802 | 0.9817 | 0.9793 | 0.7306 | 0.7298 | 0.7315 | 0.9863 | 1 | 0.9848 | 0.6437 | 0.642 | 0.6138 |
| SK-N-SH DNA R3 | 0.9858 | 0.9844 | 0.9843 | 0.8294 | 0.8176 | 0.8121 | 0.9863 | 0.9872 | 0.9863 | 0.7633 | 0.7635 | 0.764 | 0.9945 | 0.9848 | 1 | 0.6567 | 0.6554 | 0.6374 |
| SK-N-SH RNA R1 | 0.6446 | 0.6419 | 0.6451 | 0.6263 | 0.6176 | 0.6136 | 0.6486 | 0.6508 | 0.6492 | 0.801 | 0.8008 | 0.7991 | 0.658 | 0.6437 | 0.6567 | 1 | 0.996 | 0.9914 |
| SK-N-SH RNA R2 | 0.6419 | 0.6398 | 0.6414 | 0.6239 | 0.6166 | 0.6119 | 0.6474 | 0.6496 | 0.6485 | 0.8041 | 0.8037 | 0.8024 | 0.6562 | 0.642 | 0.6554 | 0.996 | 1 | 0.9933 |
| SK-N-SH RNA R3 | 0.6144 | 0.6122 | 0.6136 | 0.6118 | 0.6051 | 0.6013 | 0.6286 | 0.6298 | 0.6293 | 0.8066 | 0.807 | 0.8049 | 0.6374 | 0.6138 | 0.6374 | 0.9914 | 0.9933 | 1 |
